# The Genetic History of the South Caucasus from the Bronze to the Early Middle Ages: 5000 years of genetic continuity despite high mobility

**DOI:** 10.1101/2024.06.11.597880

**Authors:** Eirini Skourtanioti, Xiaowen Jia, Nino Tavartkiladze, Liana Bitadze, Ramaz Shengelia, Nikoloz Tushabramishvili, Gunnar U. Neumann, Raffaela Angelina Bianco, Angela Mötsch, Kay Prüfer, Thiseas C. Lamnidis, Luca Traverso, Claudia Sagona, Luka Papac, Wolfgang Haak, David Reich, Sturla Ellingvåg, Philipp W. Stockhammer, Johannes Krause, Harald Ringbauer

**Affiliations:** Department of Archaeogenetics, Max Planck Institute for Evolutionary Anthropology, Leipzig, Germany; Max Planck Harvard Research Center for the Archaeoscience of the Ancient Mediterranean (MHAAM), Leipzig, Germany; Anthropological Laboratory, Ivane Javakhishvili Institute of History and Ethnology, Tbilisi, Georgia; Centre for Archaeological Researches, Ilia State University, Tbilisi, Georgia; Tbilisi State Medical University, Tbilisi, Georgia; Explico - Historical Research Foundation, Norway; Department of Genetics, Harvard Medical School, Boston, MA 02115, USA; Department of Human Evolutionary Biology, Harvard University, Cambridge, MA 02138, USA; Institute for Pre- and Protohistoric Archaeology and Archaeology of the Roman Provinces, Ludwig Maximilian University, Munich, Germany; School of Historical and Philosophical Studies, University of Melbourne, Victoria 3010, Australia

**Author notes:** These authors contributed equally (Eirini Skourtanioti, Xiaowen Jia).

## Abstract

Archaeological and archaeogenetic studies have highlighted the pivotal role of the Caucasus region throughout prehistory, serving as a central hub for cultural, technological, and linguistic innovations. However, despite its dynamic history, the critical area between the Greater and Lesser Caucasus mountain ranges, mainly corresponding to modern-day Georgia, has received limited attention. Here, we generated an ancient DNA time transect consisting of 219 individuals with genome-wide data from 47 sites in this region, supplemented by 97 new radiocarbon dates. Spanning from the Early Bronze Age 5000 years ago to the so-called ‘Migration Period’ that followed the fall of the Western Roman Empire, we document a largely persisting local gene pool that continuously assimilated migrants from Anatolia/Levant and the populations of the adjacent Eurasian steppe. More specifically, we observe these admixture events as early as the Middle Bronze Age. Starting with Late Antiquity (late first century AD), we also detect an increasing number of individuals with more southern ancestry, more frequently associated with urban centers – landmarks of the early Christianization in eastern Georgia. Finally, in the Early Medieval Period starting 400 AD, we observe genetic outlier individuals with ancestry from the Central Eurasian steppe, with artificial cranial deformations (ACD) in several cases. At the same time, we reveal that many individuals with ACD descended from native South Caucasus groups, indicating that the local population likely adopted this cultural practice.

## Introduction

Situated at the crossroads of Eastern Europe and Southwest Asia, the South Caucasus has long been recognized as a seminal center of cultural and technological innovation throughout prehistory ^1–4^. Despite being surrounded by formidable geographical barriers, such as the Greater Caucasus Mountain Range, the area constituted an isthmus connecting various civilizations ^1–3^. As early as the Bronze Age (BA; starting c. mid-4^th^ millennium BC), the distinct traditions of the cultural complexes of Maykop and Kura-Araxes suggest a remarkable degree of cultural connectivity across the area south and north of this mountain range ^4,5^.

Previous ancient DNA (aDNA) studies have revealed that the Caucasus formed no lasting barrier to human movement. The populations of the North Caucasus inhabiting the so-called piedmont region were genetically connected to the south since the Eneolithic ^6^, while the prehistoric populations of the South Caucasus and Anatolia extensively mixed since the 7^th^ millennium BC ^7^. Conversely, the impact of mobility from BA steppe pastoralists into the South Caucasus has been attested using aDNA from the Middle/Late BA (c. 1900-1200 BC) onwards in Armenia ^8^- and also in eight genomes from two Late BA (LBA) sites in Georgia ^9^.

Right after the formation of stratified societies in the Middle BA, the rise of political authorities in the LBA and Early Iron Age has been postulated ^10,11^. Beginning around 800 BC and coinciding with the linguistic diversification of proto-Karto/Georgian-Zan (a distinct language family from Indo-European) into Georgian and Zan^12^, the social and economic division of the region of present-day Georgia deepened, culminating in the formation of the first states of Colchis in the Iron Age, and the Kingdom of Iberia in Early Antiquity, respectively ^12^. Soon, Colchis was involved in the conflict against the Urartian and subsequent Median empires and finally fell under the rule of the Persian Achaemenid Empire ^13^. No later than the 6^th^ century BC, foreign contacts through the Greek colonies were established on the Black Sea coast ^14^.

On the eastern side of Georgia, the kingdom of Iberia was founded in the aftermath of Alexander’s conquest and, together with Colchis, were under the suzerainty of the Seleucid empire. Around 65 BC, the Romans conquered Colchis from the Kingdom of Pontus ^15^ and the Kingdom of Iberia, exposing it to the Graeco-Roman world and ultimately claiming both regions under the principate ^14^. The early Christianization of Iberia started in the 4^th^ century AD ^13^ and remained strong considering the fluctuating political boundaries elsewhere on Rome’s eastern frontier ^16^. Since the 3^rd^ century, initially Romans and Sassanians, and later Byzantines and Arabs, vehemently competed for Iberia and Colchis (later called Lazica) until the great medieval Kingdom of Georgia was unified in 1008 ^17^.

The South Caucasus has been a melting pot of diverse ethnic groups from as far away as Inner Asia and Eastern Europe, including Sarmatians and the Alans ^18,19^ and later the Huns during the migration period of the Early Middle Ages ^20,21^. These nomadic people of the steppes practiced certain types of artificial cranial deformations (ACD) ^22^. This practice also appears in the region of Georgia, evidencing influence from the north across the Caucasus in different periods ^39,40^.

It remains unclear how the dynamic history of the South Caucasus region drove human mobility at the individual and group levels. Here, we generated an ancient genome-wide dataset of 219 individuals from 47 sites in Georgia. Importantly, we also produced 97 same-sample radiocarbon dates that confirm that sampling spans *c.* 4,500 years continuously from the EBA to the Early-High Middle Ages. This extensive time transect fills several critical gaps in archaeogenetic sampling for the area between the Greater and Lesser Caucasus, thus far represented by only eight Bronze Age ^9^ and two Upper Palaeolithic genomes (‘Caucasus hunter-gatherers’) ^23^.

This new data enabled us to trace individual and group-level ancestry patterns through time. Our findings reveal a high degree of genetic continuity, along with lasting mixture events since the MBA with steppe-related and Anatolian/Levantine groups. From the early Christianization of the Iberia kingdom, we observe a high rate of individuals with ancestry from Anatolia or the Levant. Notably, we find that most of the individuals with ACD descended from native South Caucasus groups, highlighting a cultural practice adopted by the local population. Finally, by screening for identical-by-descent (IBD) segments, we establish higher levels of biological relatedness among individuals from rural areas buried a few kilometers apart, compared to large urban centers like the ancient capital of the Iberia kingdom. By unraveling the intricate tapestry of population movements and cultural exchanges, our results contribute to a better understanding of the region’s rich and diverse heritage.

## Results

### Compiling and authenticating the ancient DNA dataset

We converted drilled bone and tooth powder from 359 samples of human remains into single-stranded genomic libraries, estimated endogenous DNA preservation by shotgun-sequencing a portion of the libraries (n=96), and proceeded with enrichment for 1,233,0123 variable sites (SNPs) in the human genome (‘1240k’ panel) for 343 samples (**Tables S1-S3**). After quality control, we removed libraries with low sequencing coverage on the targeted positions (<20,000 SNPs covered; see Methods) and substantial contamination rates (>10%) and merged libraries from genetically identical individuals. After this QC, we retained a dataset of 219 unique individuals with newly generated genome-wide data (**Methods, Table S1**). Where factors such as female sex, coverage, mt/nu ratio, and terminal deamination (aDNA damage) prevented a reliable autosomal contamination estimate, we kept these cases in our dataset but excluded them from our group-based analyses and interpreted the other results cautiously.

Our final dataset originates from 47 sites across present-day Georgia (**SI Material 1**), 22 located close to Mtskheta and Tbilisi, the former and last capital of the Kingdom of Iberia, respectively (**Figure 1A**). By unifying the sampling range from present-day Armenia to the North Caucasus and the adjacent steppe (**Figure 1B**), our results synthesize into a larger picture of the population dynamics in the area from prehistory to historical periods. Several of the earliest individuals are associated with the Kura-Araxes culture (the site of Kiketi), a Bronze Age cultural complex that extended throughout the Southern Caucasus. Despite its importance and many unresolved consequential questions, such as its linguistic makeup, published ancient DNA only covers the southern and eastern end of this range ^6,8^. Finally, the most recent transect in our dataset (Early Middle Ages) is a unique addition to the Caucasus region that allows us to extend our understanding of the impact of Christianization and the subsequent ‘Migration Period’ on human mobility and population genetics beyond Europe ^24–27^. For some analysis, we partitioned the dataset based on time periods (see Methods; *Data compilation*). We note that a discretization of the dynamic past of the South Caucasus is challenging due to diverse cultural and archaeological contexts. Therefore, our grouping is based solely on calendar dates, with the main goal of increasing the statistical power of time series analysis.

**Figure 1.**
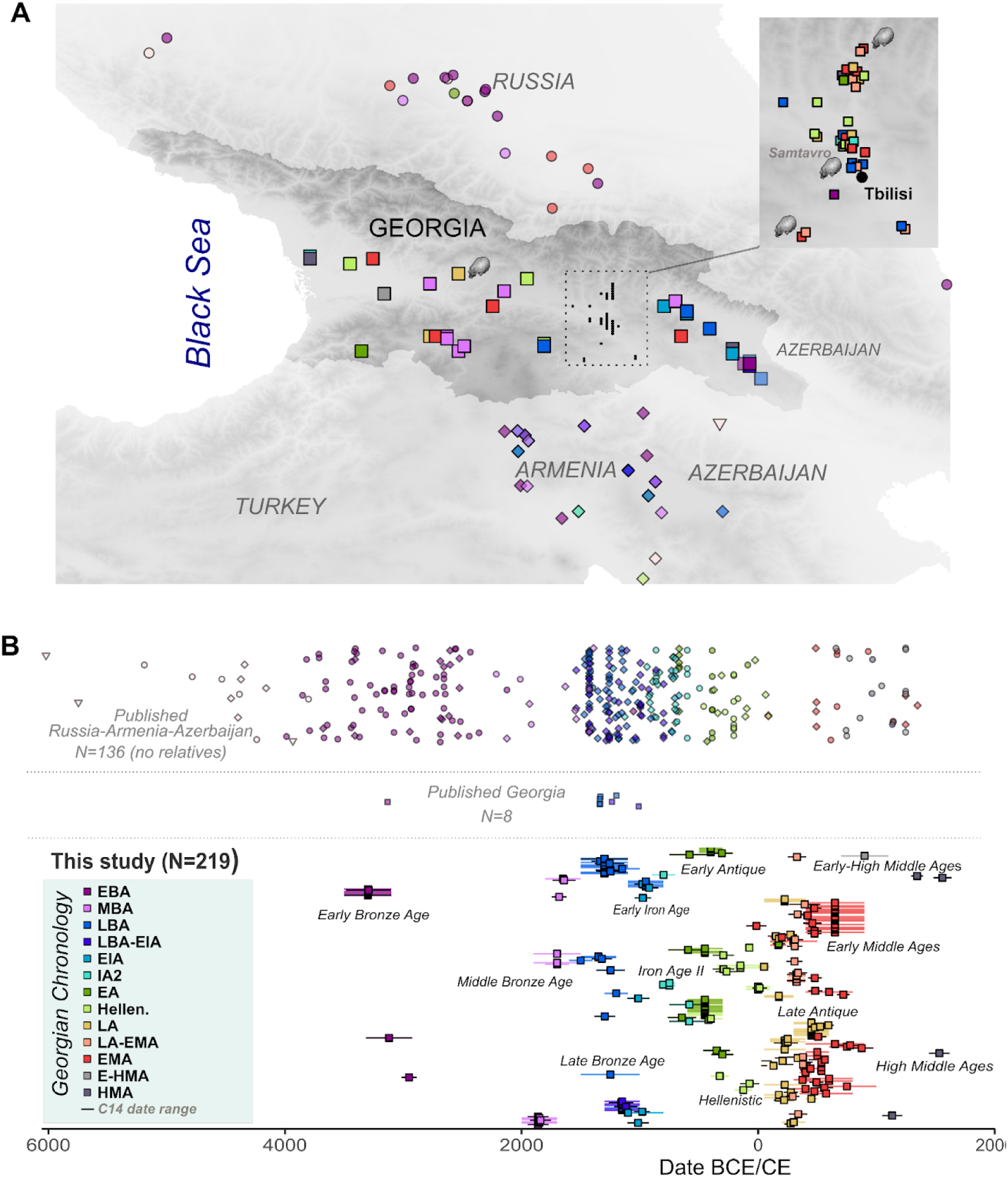
Spatiotemporal information of new and published data analyzed in this study. **A.** We annotate the geographic location of new genomic data from Georgia in squares, with solid colors corresponding to their chronological period. Published data from Georgia and surrounding territories (Armenia, Azerbaijan, Russia, and Turkey) are also indicated in transparent colors. We also highlight locations with artificially elongated skulls**. B.** Date and archaeological period information as an average for published data and as a range for the new samples. ^14^C date ranges are annotated in black lines. Abbreviations of the periods discussed are provided.

### Persisting local gene pool and limited gene flow in the South Caucasus

To explore genetic ancestry, we first assessed the genome-wide data by projecting the time-transect and other relevant ancient DNA datasets onto the first two axes of a principal component analysis that we constructed from 77 present-day populations from Western Eurasia (**Figure 2A**), including several modern groups from the Caucasus (see Methods). We find that more than 90% of the individuals join the same cluster outlined by previous studies presenting Eneolithic to Early Iron Age individuals from the southern and northern Caucasus (excluding the steppe environment to the north) ^6,9,28,29^. Notably, this ancient pc1-pc2 cluster also overlaps with modern populations from the southern Caucasus. None of the individuals from the Bronze to Iron Age phases fall into the distinct ‘Caucasus-Steppe BA’ cluster or a cline of mixed ancestry from the steppe to the southern Caucasus that was described previously ^6^. This signal indicates that this pc1-pc2 cloud comprehensively represents the genetic variability of the local South Caucasus ancestry.

**Figure 2.**
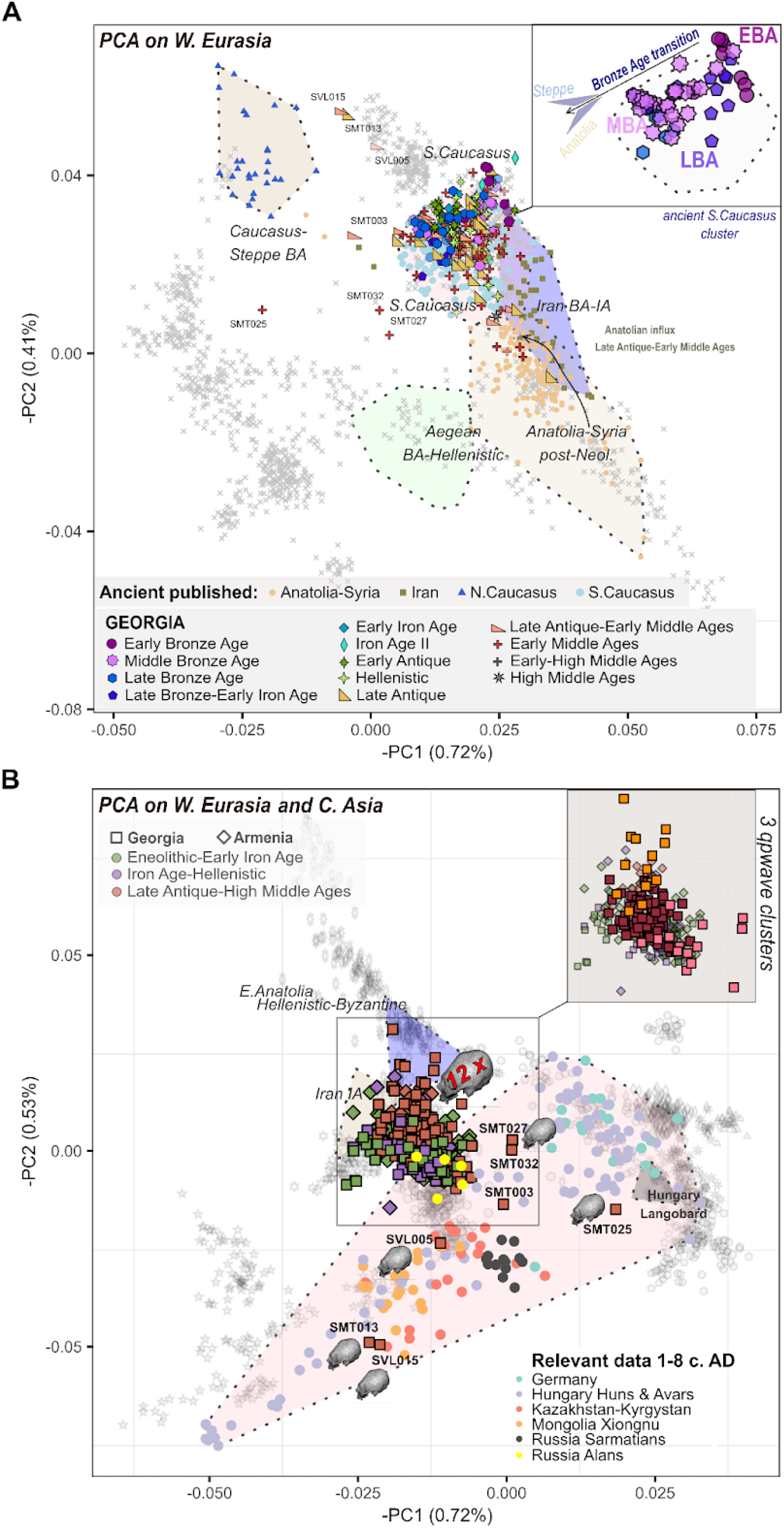
Principal component analysis in two different datasets. **A.** PCA on 1300 present-day individuals from W. Eurasia. The dataset of 221 from Georgia is projected with the ‘*lsqproject*’ parameter of smartpca and annotated by period (color and shape). Fill color transparency is applied for individuals with low coverage and/or questionable quality (e.g., contamination rates). Other relevant ancient datasets are projected as well. **B.** PCA on 1486 present-day individuals from W. Eurasia and C. Asia. Published ancient groups are plotted when their members have similar PC1 and PC2 coordinates and are dated close to the ancestry outliers from Georgia. Individuals with artificial cranial deformations are annotated with respective icons. PC1 and PC2 variability in post-Hellenistic Georgia is also detected with cluster analysis with pairwise *qpWave* tests that indicate three different clusters at the plot’s upper-right corner.

Within this cluster of local ancestry, however, we observe a temporal substructure throughout the BA, with EBA individuals occupying higher pc1 and pc2 values compared to M/LBA individuals (**Figure 2A**). Moreover, several individuals are placed outside this main cluster. They all date from the Late Antique and Early Middle Ages (n=17) and can be further grouped into two components: a. those on a continuum of increasing genetic affinity to Anatolia/Levant (n=10), and b. those with diverse ancestry backgrounds (n=7, referred to throughout the text as ‘ancestry outliers’). Remarkably, four out of seven of the ancestry outliers are individuals whose preserved skulls attest to the practice of ACD (**Figure 2B**).

To better understand the geographic origin of the ancestry outliers (or of their recent ancestors), we computed another PCA that also included present-day individuals from Central Asian populations (see Methods) and projected the ancient ones as before. This PCA shows that the two individuals SMT013 and SVL015 (c. 3^rd^ and 4^th^ centuries AD, respectively) descended from Central Asian populations (**Figure 2B**), a conclusion further supported by another PCA on all available Eurasian populations (**Supplementary Figure 1A**). Projecting only ancient groups with similar pc1 and pc2 coordinates and dates to our ancestry outliers reveals a gradient of ancestries from European to Central/Northeast Asian, which is also visible in individuals from the Pannonian Basin starting with the invasion of the Huns and later the Avars ^30,31^. This observation suggests that the ancestry outliers, who were all buried in two urban contexts within a short distance of each other (Samtavro and Samshvilde), may also be linked to related contexts.

To further evaluate the ancestry patterns in the PCA, we performed explicit admixture modeling using *qpAdm/qpWave* to address four main objectives: 1. deconstruct ancestry as broad contributions from earlier Southwest Asian and East European populations and compare the composition across time periods (distal modeling) 2. explicitly model differences across periods using as sources populations that could have been donors (proximal modeling), 3. quantify genetic variability among individuals, and 4. investigate alternative models (admixture) for the ancestry outliers.

For the distal modeling, we found that all groups from Georgia and Armenia can be modeled with sources from the earliest sources from Neolithic Anatolia, the hunter-gatherers from Georgia (CHG), Chalcolithic Iran (west; Seh Gabi), Pre-Pottery Neolithic (PPN) Levant, and hunter-gatherers from Eastern Europe (EEHG) (**Supplementary Figure 2A**). After corroborating previous findings that CHG and Chalcolithic Iran ancestries are competing sources that are challenging to distinguish ^8^, we combined them into one source (**Figure 3A**). We then modeled the ancient populations of Georgia and Armenia as temporal groups based on chronology. This analysis shows low levels of ancestry from EEHG already during the EBA (7.7±2.4% and 6.3±1.9%, respectively, where ± denotes one standard deviation), while per-individual models had less power to detect it ^8^. However, it is only from the Middle Bronze and the Early Iron Ages that EEHG-related ancestry substantially increases to 13±1%, as does the coefficient from Neolithic Anatolia in Georgia (from 16±2.5% to c. 30±1.2% during the Late Antique and Early Middle Ages). Notably, in parallel to the increasing EEHG-related ancestry in the MBA, we also document the first appearance of males with the Y-haplogroup R1b-Z2103 (**Supplementary Figure 4, Table S1**), a lineage strongly associated with the Yamnaya, Catacomb and North Caucasus culture groups and their expansions such as the Altai Afanasievo ^28,32,33^.

**Figure 3.**
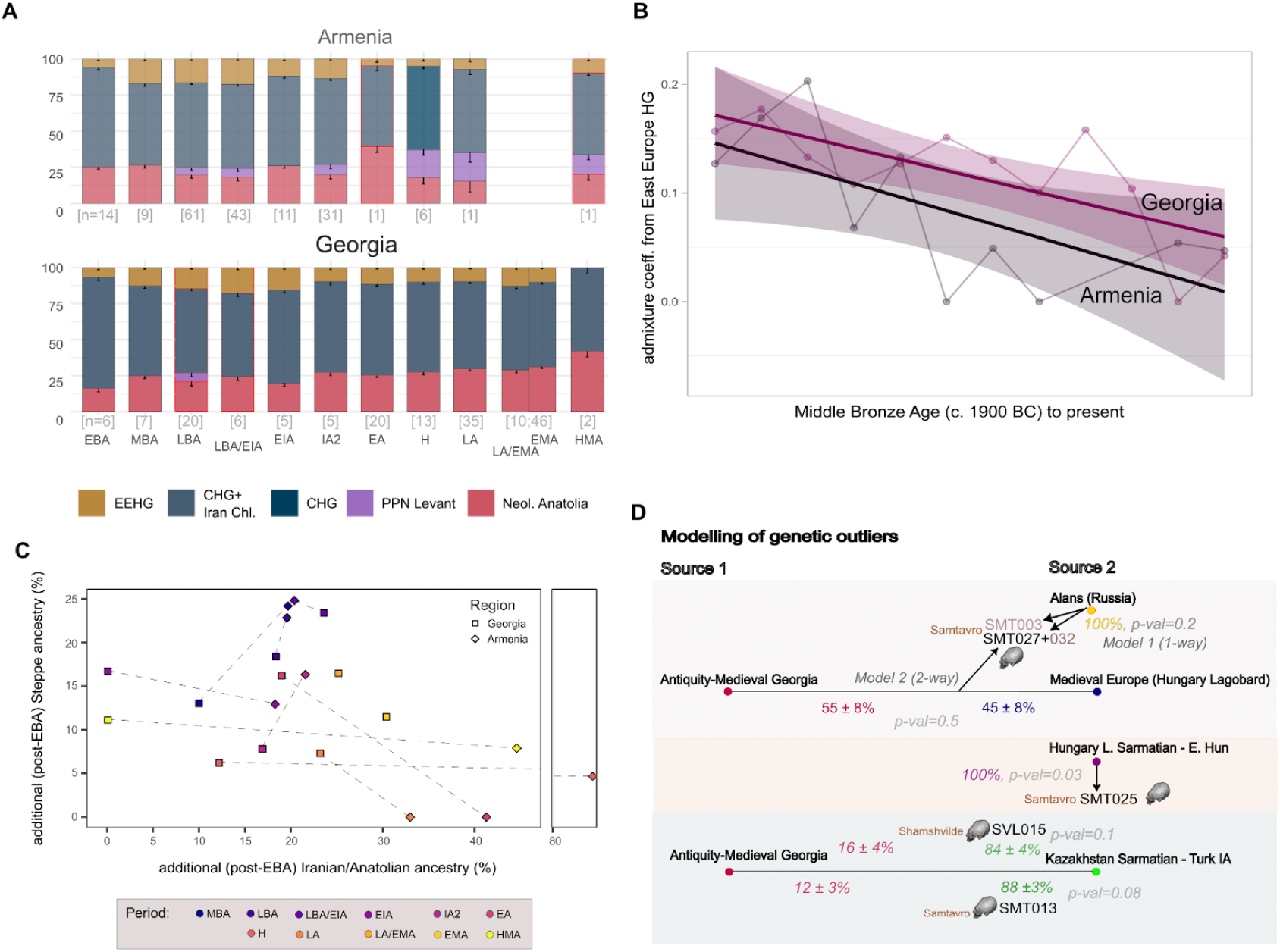
Admixture modeling with *qpWave/qpAdm*. **A.** *qpAdm* coefficients with -1SE are estimated for every period in Georgia and respectively in Armenia, using distal sources from Anatolia/Levant, Iran, the Caucasus (hunter-gatherers) and the W. Eurasian Steppe (Eastern hunter-gatherers). For some groups, adequate models are reached with CHG instead of Chl. Iran and vice-versa therefore, the two groups were merged into one source. Red outlines in the bar plots indicate models of marginal fit (0.01≤p-value<0.05). Periods are in two-to-three-letter abbreviations. **B.** Linear regression for EEHG-related ancestry in both Armenia and Georgia indicates a decrease in this ancestry since the Middle Bronze Age (MBA). **C.** Proximal *qpAdm* models in post-Early Bronze Age (EBA) S. Caucasus. EBA Georgia is used as the main source. With few exceptions, groups from all the periods require additional ancestry both from BA Steppe (here modeled with ‘Steppe_Catacomb’) and Anatolia (modeled here with ‘Turkey_Ikiztepe_LateC’). In the case of Hellenistic Armenia, Anatolia and Iran (‘Iran_Hasanlu_IA’) can adequately model the target with only minor components from EBA Armenia and Steppe. **D.** Adequate one-to three-source mixture models with *qpAdm* for the genetic outliers (p-value ≥ 0.05). Different possible combinations were tested informed by PC coordinates, IBD analysis, historical hypotheses, and the assumption that some of these outliers might still derive part of their ancestry locally (see also **Table S8**).

Starting in the Iron Age, EEHG ancestry continuously declined -more substantially in Armenia than in Georgia-but reached equally low levels in modern-day Armenians and Georgians (**Figure 3B, Supplementary Figure 2A**). Using EBA individuals from the South Caucasus (Armenia and Georgia) to represent the local ancestry, we were able to model all post-EBA groups with additional ancestry from a steppe population north of the Caucasus (‘Russia_Steppe_Catacomb’) and a northeast Anatolian population (‘Turkey_Ikiztepe_LateC’). In the M/LBA, the proportion of the steppe-related ancestry are slightly higher than those from Anatolia, but overall, these models indicate that the observed shift in pc1 and pc2 values from EBA to M/LBA results from two different admixture events from opposite geographical directions and of comparable magnitude. From the Hellenistic period onwards, our models show more extensive gene flow from the south, particularly into Armenia (**Figure 3C)**, as also seen in the increasing Levantine component in the distal modeling. In this line, we also find that Levantine sources (‘Lebanon_Hellenistic.SG’ and ‘Lebanon_ERoman.SG’) rather than Anatolian/Iranian adequately model Hellenistic and Late Antique Armenia (**Table S4**).

We next explored whether genetic diversity spiked in specific periods, which would evidence recent admixture events or high individual mobility. Using *qpWave*, we find that in every period, some individuals cannot be modeled as being cladal with each other (i.e., genetically similar with respect to a set of reference populations), with an overall proportion of non-cladal models with p-value<0.01 of 10% or more (**Supplementary Figure 2B**). This proportion is highest for the periods from the Late Antique to the Early Middle Ages, reaching statistical significance in the transitional group when compared to the earliest phases (e.g., Bronze Age groups). Together with the ancestry outliers - not included in these tests - and individuals with increasing affinity to Anatolia, these results highlight that human mobility intensified starting in the Late Antique period. The increased diversity in Y further corroborates this idea (**Supplementary Figure 4**). We illustrate the results of the *qpWave* tests as a clustered heatmap of the p-values for pairs from the Hellenistic period to the Early Middle Ages (**Supplementary Figure 3A**). Two clusters, each consisting of 14 individuals, separate from the remaining 108 and correspond to a substructure also partially captured in the second PCA (**Figure 2B**). The first cluster (pink square symbols) consists of individuals with a higher affinity to north Caucasus/steppe groups, as indicated by *F_4_-statistics* (**Table S5**). Conversely, the second cluster (orange symbols) suggests that the high affinity to Anatolia/Levant in some individuals already visualized in the first PCA could extend to a cline with mixing southern ancestry into the local gene pool. We note that in both cases, the use of the term ‘cluster’ refers to its members being similarly distinct from the other individuals, but they cannot be treated as genetically homogeneous groups.

Overall, the results of PCA and *qpAdm/qpWave* delineate admixture events that were continuously assimilated into a persisting local gene pool, preserving the overall population structure in the South Caucasus since the Bronze Age with only subtle shifts on the whole-population level.

### Connectivity within the South Caucasus and beyond

Following the evidence from *qpAdm* for ‘northern’ and ‘southern’ gene flow into both Georgia and Armenia, we screened for Identical by Descent (IBD) sharing among ancient genomes using the software *ancIBD* (see Methods). First, we show that the average IBD sharing rate within Georgia and Armenia and between them overlap and continuously decay with geographical distance at the same rate (**Figure 4A**). This continuous “isolation-by-distance” pattern ^34^ suggests that - despite the different political and territorial influences between present-day Armenia and Georgia in the past - the populations mixed at a rate proportional to their geographical distance. By screening IBD sharing of all pairs containing one ancient Georgian individual from our dataset and another individual buried up to 1500 km away, we quantify the connections with the rest of Southwest Asia (‘South’) and Eastern Europe to Central Asia (‘North’) (**Figure 4B**). At shorter geographical distances (i.e., 100-240 km), we observe an increased rate of short IBD sharing with the south (i.e., Armenia), corroborating increased population connectivity within the South Caucasus compared to across the Caucasus, which therefore likely served as a barrier to mobility. However, at larger distances, rates of IBD sharing equalize between south and north, reflecting the global admixture proportions estimated with *qpAdm* and the rates of outliers whose ancestry is traced on either side. These signals indicate that the Caucasus mountain range was not a lasting barrier to long-distance mobility.

**Figure 4.**
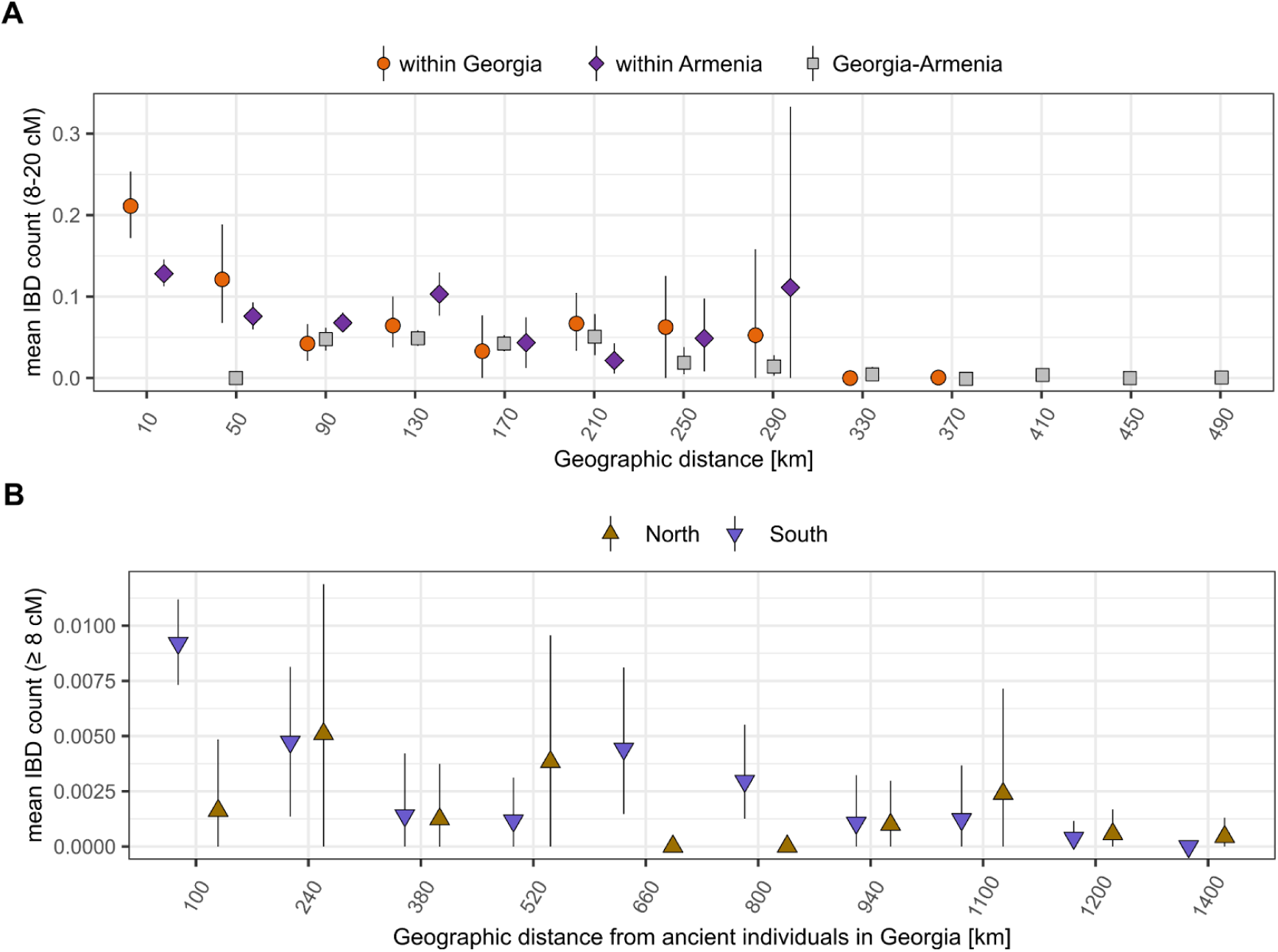
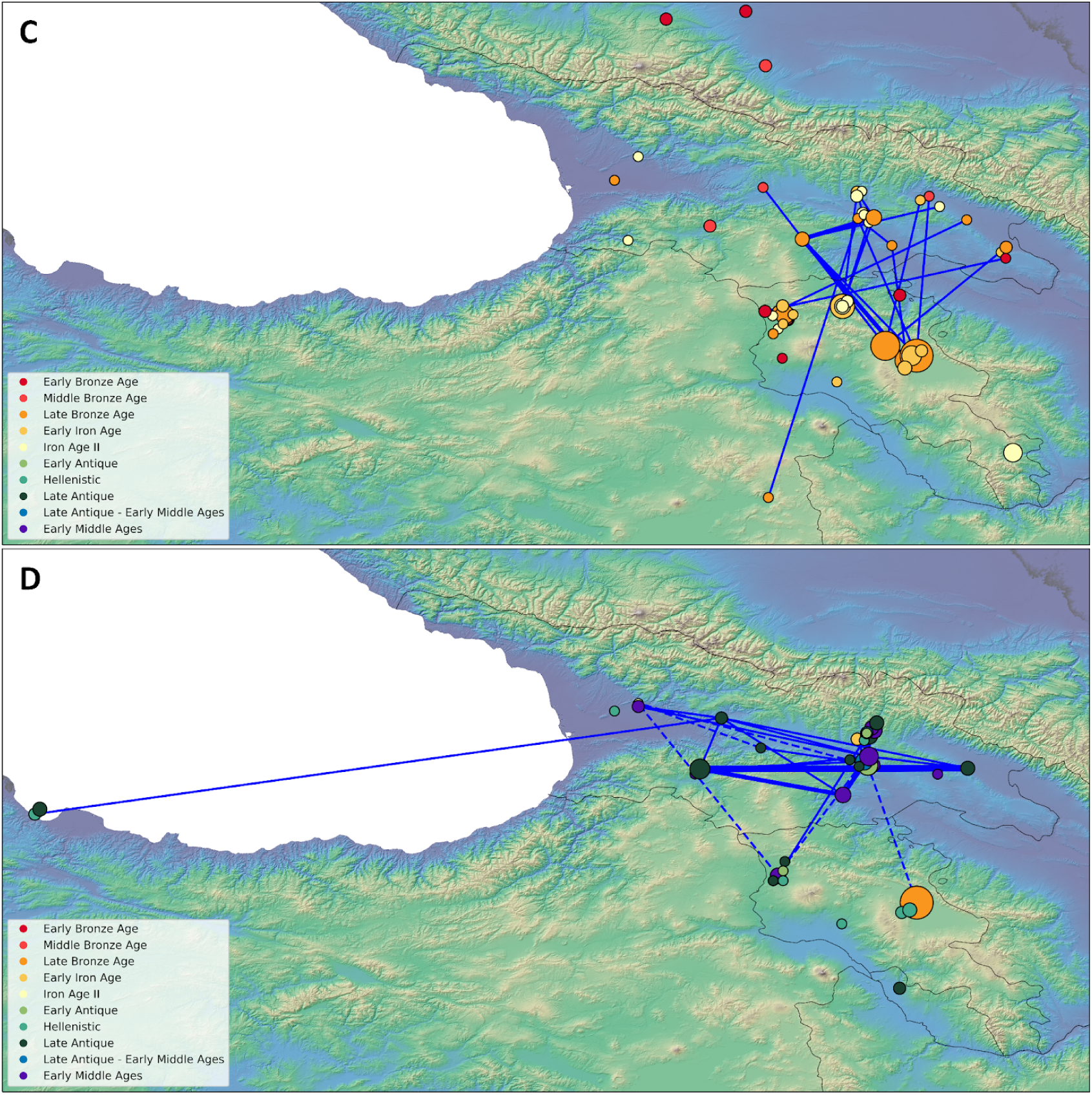
Spatial distribution of IBD connections for Georgia and Armenia. **A.** Sum of IBD counts for all pairs including Georgia and Armenia (*c.* 20,000) with IBD sum ≤ 200cM and date difference of up to 1000 years plotted in bins of increasing geographic distance. **B**. Distant connection between ancient individuals from Georgia and populations North and South of the Greater Caucasus presented as the mean count of IBD ≥ 8cM and geographical distance from *c.* 100 to 1500 km. Pairs with a date difference of more than 1000 years were removed, retaining a total of *c.* 35,000 pairs. The 95% confidence interval for every bin was calculated with the embedded bootstrap function ‘mean_cl_boot’ of ‘stat_summary’ in R (ggplot). **C.** Map of the inter-site IBD connections (≥12cM) for Georgia in early periods (Early Bronze Age to Iron Age II). The size of the symbols indicates the sample size of each site, while the line thickness represents the number of IBD pairs between two sites. **D.** Map of the inter-site IBD connections (≥12cM) for Georgia in late periods (Early Antique to Early Middle Ages). Dashed lines indicate the IBD connections between early and late periods. The IBD connections that were identified between the three closely located sites, Aragvispiri, Dzinvali, and Nedzikhi, have been filtered out.

### Varying population dynamics between rural settlements and urban centers

We then analyzed IBD sharing ≥12cM long, which evidences recent genealogical links because IBD segments of this length originate from relatives connected within several hundred years ^35^. We observed that such sharing was common across all periods and distances (**Figure 4C and 4D**). However, we detected remarkable local patterns. The dense sampling in eastern Georgia - the area corresponding to the eastern territory of the Kingdom of Iberia - and the distribution of IBD connections in the Late Antique and Early Middle Ages allows us to investigate two groups of sites with contrasting niches of different social organization. We present a network analysis of the IBD segments connecting Samtavro, Samshvilde, and Fiqris Gora, named in order of importance as urban centers and pilgrimage sites, while the sites of Nedzikhi, Dzinvali, and Aragvispiri are all associated with rural settlements located within a very short distance from each other (**Figure 5A**).

**Figure 5.**
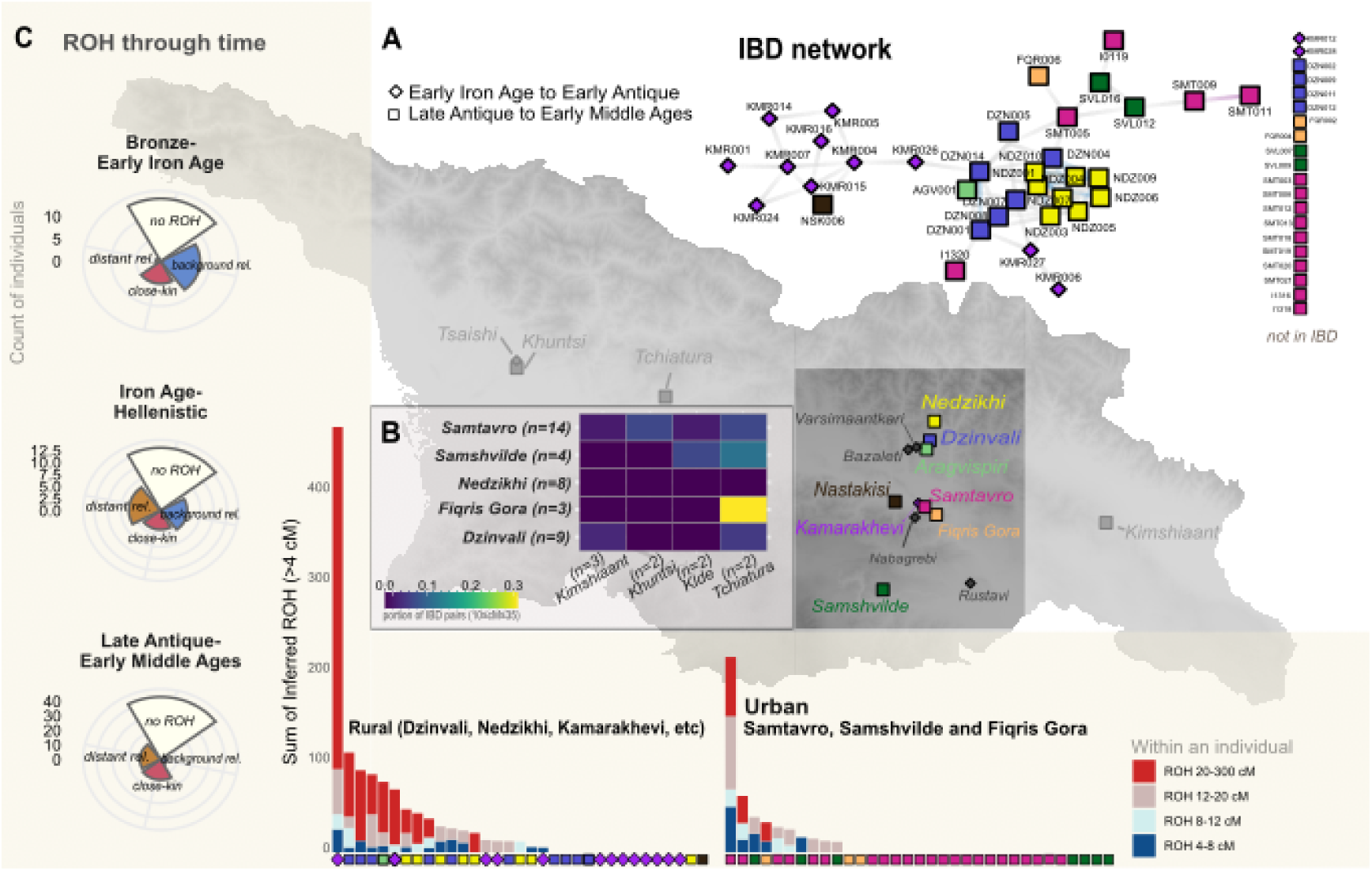
ROH and IBD network analyses. **A.** Network of IBD connections in Eastern Georgia where sampling is particularly dense. In the earlier period (Iron Age to early Antiquity), only individuals from the site of Kamarakhevi exhibit a high rate of intra-IBD connections within and also with sites across Georgia as shown in the panel. **B**. During the Late Antique and Early Middle Ages, there was an asymmetry in IBD distributions caused by some sites where all individuals from within and between sites associated with settlements of rural activity are distantly related. In contrast, nearby sites with urbanism like Samtavro exhibit minimal IBD within but higher with more distant sites as shown in panel B. **C**. Runs of homozygosity from the sites in the IBD network suggest a correlation of IBD connections with consanguinity revealing a contrast of more endogamic rural communities (with both ample long ROH and intra-site IBD sharing) living side-by-side with large cosmopolitan populations from urban centers in the Late Antique/Early Middle Ages (with little long ROH and intra-site IBD sharing). However, the overall levels of consanguinity have been stable over time (leftmost panel).

Samtavro, located in the old capital and the urban site Mtskheta, shows the lowest rate of internal IBD sharing, with only 4 of 14 individuals connected via IBD. Overall, intrasite genetic relatedness estimated with *ancIBD* and *BreadR* is exceptionally low in Samtavro, with only two pairs of parent-offspring and no pairs of distant relatives up to the sixth degree (**Figure 5A, Tables S1 and S6**). Remarkably, SMT005, the individual with the most intersite connections, is also the only one from Samatvro with a signal of consanguinity (sum ROH>200cM; **Figure 5C; Table S7)**. In addition to the high genetic diversity within the site shown with PCA, *qpWave* clustering, and the lack of internal biological relatedness, Samtavro displays the highest rate of IBD sharing with four sites at larger geographic distances (**Figure 5B**). Conversely, Dzinvali and especially Nedzikhi’s IBD sharing already declined at a short geographic radius. Taken together, these signals suggest that the urban scale of Samtavro was also reflected in the organization of its necropolis, whereas in nearby settlements, people maintained a social structure of smaller and, to some extent, endogamic communities.

We observe signals of markedly different social organizations outside the urban sites as early as the Iron Age. In the site of Kamarakhevi, located next to cosmopolitan Samtavro, as many as nine of 11 individuals share IBD between them (**Figure 5A**). One of them presents one of the highest amounts of long ROH in the ancient DNA record, equivalent to parents related as either first or second-degree relatives. Continuing into the Late Antique and Early Middle Ages, all eight individuals from the rural site Nedzikhi show pairwise IBD connections ranging from the second to the tenth degree. We also show that four individuals from Nedzikhi exhibit short to intermediate runs of homozygosity, consistent with a population with substantial background relatedness. The remaining three have long ROH equivalent to close-kin unions (e.g., first and second cousins), indicating that all individuals sampled from this site represent an endogamous community. At rural Dzinvali - coeval and at a close distance to Nedzikhi -, the absence of ROH in half of the individuals suggests that the group was part of a larger mating pool or an endogenous community with new members joining. The *qpWave* and PCA results (**Supplementary Figure 3**) further support this idea: Individuals DZN002, DZN009, and DZN011, who are not related to anyone via IBD sharing, cluster separately from the Late Antique group due to genetic affinities from either Anatolia or possibly north of the Caucasus. The same applies to FQR002 from urban Fiqris Gora, one of the individuals with the highest affinity to Anatolia, which contrasts with the IBD-connected and consanguineous individual FQR006.

Overall, the levels of consanguinity, defined as the proportion of close (up to first cousins) and distant (second cousins or equivalent) paternal relationships, remain constant through time at a rate of ∼25-30% (**Figure 5C**), which is elevated compared to the ancient DNA record ^36^. Precisely, we observe consanguinity in 16 out of 125 individuals analyzed for long ROH (Methods). The frequency in Samtavro is lower (1/25); however, the frequency of individuals consistent with distant parental relatedness (5/20) is comparable to the levels in the overall ancient post-Iron Age population (18/81). The decline in short ROH from a maximum in the Bronze Age pastoralists to a minimum in the Late Antique and Early Middle Ages shows a general trend towards locally larger population sizes, particularly in urban areas, while smaller communities continued to exist.

### Insights into Artificial Cranial Deformations (ACD) from aDNA

Within our dataset, 17 individuals from Samtavro, Samshvilde, Nedzikhi, and Tserovani had artificially elongated skulls. Artificial Cranial Deformations (ACD), a form of skull modification, has occurred independently throughout the world and as early as the Early Neolithic in Southwest Asia, where it is thought to have been widespread ^37,38^. In Georgia, the earliest documented ACD cases date to the Late Bronze Age - Early Iron Age transition ^39^, but the frequency substantially increased in the Mtskheta area during the Migration Period of Europe (4^th^ - 7^th^ century AD) ^40^. In particular, craniological material from early medieval Georgia suggests a frequency of c. 20% (126/579 preserved skulls), and even higher in the Samtavro cemetery (c. 30%; 43/132) ^39^ Furthermore, within one type of ACD, Georgian skulls show higher diversity than those associated with the Huns, challenging the widespread idea that the practice spread due to Hunnic influence ^40^.

Here, we associate information about genetic ancestry and ACD typology on 16 adult individuals from the end of the Late Antique period (late 3^rd^ century AD) to the Early Middle Ages. According to our PCA and IBD network analyses (**Figures 2B and 5A**), the ACD cases from Nedzikhi, which comprise a third of our dataset, occurred among individuals from a close mating network within the local population. Distantly linked to this network, an individual from Samtavro with ACD (SMT009) is the mother or daughter of another individual with ACD (SMT010). Both have ancestry profiles from the South Caucasus (called ‘local’ hereafter). Although they were subjected to different elongation techniques, the evidence of lineal kinship suggests that the practice was common among the local population and passed on through genealogical transmission.

Despite the majority of individuals with ACD (12 out of 17 individuals) carrying ‘local’ genetic ancestry, four ACD cases of tabular (n=3) and circular (n=1) “Hunnic” type are clear ancestry outliers in our dataset. The genetic backgrounds of these individuals are very diverse, ranging from Central/Eastern Europe (SMT025), Central/Inner Asia (SMT013 and SVL005), and East Europe/Central Asia (SVL015). Their ^14^C date span is 220-525 cal AD (**Table S1**), which partially predates the widespread presence of Huns in Eastern Europe (*c.* 370-460 AD). More specifically, the tool *mobest*, using PCA and spatiotemporal coordinates as input (see **Methods**), estimates higher similarity probabilities for the Carpathian Basin for SMT025, the Caucasus - including part of the Pontic-Caspian steppe - for SVL005, present-day Kazakhstan, Mongolia, and southeast China for SVL015 and SMT013 (**Supplementary Figure 1C, 1D and 1E**). For ACD individual SMT027 *mobest* outputs the highest probabilities for a genetic origin within the Caucasus. In addition, SMT027 -along with SMT032 and SMT003- occupy the extreme end of the *qpWave* first cluster, plotted closer to individuals from the Alan and Sarmatian groups of Russia but also Eastern Europe (late Sarmatian to Hunnic period).

Informed by these analyses, we then formally modeled the ancestry of each genetically non-local ACD individual using *qpAdm*. Low-coverage SMT027 clusters with SMT032 (no ACD), which shows the same placement on the PCA. Together, they could be modeled as a one-source model from the pastoralist group of Alans in the North Caucasus (7^th^ cent. AD) with p-value=0.02 (**Figure 3D, Table S8**). An alternative model of a two-way mixture from LA-EMA Georgia and individuals from the Carpathian Basin from the Late Sarmatian or Langobard period is also accepted; however, no decay of local ancestry due to recent admixture could be fitted with the DATES software. For individual SMT025, adequate models (p-value≥0.05) include c. 70% ancestry related to the Carpathian Basin (i.e., Langobards) admixed with a source from the Kazakh-Pontic Steppe (e.g., Sarmatians), although adding local ancestry from Georgia (LA-EMA; c. 20%) further improved the fit of the model. Consistently, a simple one-source model - although marginally rejected with p-value=0.035 - was achieved with the group from the Late Sarmatian period in Hungary (Danube-Tisza interfluve region, 4^th^-5^th^ centuries), which was shown to be distinct from others in the area due to its higher affinity with populations from the Pontic Steppe ^31^. Individuals SVL015 (262-525 cal AD) and SMT013 (218-328 cal AD) exhibit a genetic profile common in the central and eastern steppe that has spread as far as the Carpathian Basin in the Avar period ^30^. Coeval groups from the central steppe such as ‘Kazakhstan_Berel_IA’ and ‘Kyrgyzstan_TianShan_Hun.SG’ fit neither as one-source models nor in combination with a local source (Georgia LA-EMA). The exception is the medieval group from east Kazakhstan, which serves as a majority source (c. 85%) in a two-way model with LA-EMA Georgia. The mitochondrial haplogroup H of SVL015 also supports West Eurasian ancestry. For the earliest individual, SMT013, who carries the East Asian mitochondrial haplogroup D4e1, this model can be attributed to a recent admixture event of 10±4 generations before her time (estimated with DATES). Furthermore, IBD analysis distantly connects SMT013 with one individual from Hungary from the Early Avar Period (SZOD1-187; 25cM in total) and two more from the Hun Period (KMT-2785, VZ-12673; 14 and 13cM, respectively), all of which are genetically associated with the Eastern Steppe ^30^ (**Supplementary Figure 1A and 1B**). These IBD connections suggest that SMT013 originates from the transcontinental mating network that reached Eastern Europe shortly after.

## Discussion

This study presents the first large archaeogenetic dataset from present-day Georgia. Extending previous data^6,8,41,42^, this temporally continuous genomic time transect allowed us to provide comprehensive insights into the population history of the South Caucasus.

Our comparative models for Georgia and Armenia during the Early Bronze Age demonstrate that previous mixture events between metapopulations related to the CHG and the farmers of Anatolia resulted in a homogeneous population throughout the South Caucasus not only specific to present-day Armenia. The trace amounts of EHG-related ancestry observed in EBA Armenia and Georgia indicate that it did not disappear after the Chalcolithic period when it was observed in Armenia ^43^.

We find that EHG ancestry resurged again in the Middle Bronze Age. Two lines of evidence suggest that this increase can be associated with specific populations, namely the pastoralist groups from the steppe adjacent to the North Caucasus, which are 1) fitting sources in our proximal modeling (**Figure 3C**) and also sources for the newly arriving Y haplogroup Z2103. Notably, we observe no geographically stratified signal of EHG (steppe ancestry) in the Late Bronze/Early Iron Age that could indicate an eastwards or westwards migration route across the Caucasus (**Table S9**), but one limitation is that our dataset underrepresents the western end of Georgia. The admixture timing and the model fit of the group ‘Steppe_Catacomb’ correlate with deteriorating climatic conditions in the Caucasus-Steppe region, which led to the breakdown of stock specialization and, ultimately, its abandonment at the end of the 2^nd^ millennium BC ^44,45^. In addition to such ‘push’ factors,, the farming and metallurgical societies of the South Caucasus that had developed since the 6^th^ millennium BC and could have attracted groups with similar economies. It has also been suggested that steppe mobility into the South Caucasus played a role in language innovation in Indo-European-speaking Armenia ^8^. However, recent linguistic evidence also supports a northward expansion of Indo-European languages from a region south of Caucasus postdating the divergence of the Armenian language. In the case of Georgia, Kartvelian, a distinct non-Indo-European language family that has existed since approximately 13,000 years ago ^47^, persists until today. It further diversified after the MBA/LBA admixture, and the described gene flows may have contributed to its proposed structural relationship with Indo-European languages ^48,49^.

The simultaneous rise in Anatolian/Levantine ancestry in the Middle Bronze Age indicates a separate and parallel admixture event. This gene flow likely reflects a continuous demographic process of prolonged contact between the South Caucasus and Anatolia, as previously documented ^6,7^.

Together, the two MBA admixture events mentioned above have lastingly shaped the gene pool of the South Caucasus, as evidenced by the PCA, *qpAdm* modeling, and uniparentally-inherited markers, which reached present-day population diversity levels ^50^ shortly after the Bronze Age. It is worth noting that pairwise comparisons by period reveal differences, particularly between the latest and earlier groups (see **Supplementary Figure 2A**). However, as the proximal modeling indicates, these differences can be explained by a decreasing Steppe-Anatolian/Iranian ancestry ratio over time (see **Figure 3C**). Given that Armenia and, to some extent, Iran are the southernmost enclaves of steppe ancestry in southwest Asia ^8^, this genetic signal is interpreted as an increase in gene flow from Anatolia and Iran. The historical relevance of this finding can be proposed at the base of the continuous political influence exerted on the region by the successive empires that extended across Southwest Asia, including the Median, Achaemenid, and Sasanian empires. The Iberian, Colchis, and Armenian kingdoms were among those influenced.

With half of the aDNA dataset dating to the Late Antique and Early Middle Ages (c. 50-800 AD), we were able to assess the potential influence of important historical events such as the Roman conquest, Christianization, and increased contact with nomadic tribes from the Eurasian steppe. According to a recent study ^51^, approximately 7% of historical individuals in Europe and Southwest Asia had non-local ancestry. Our results for this period broadly fit this observation, but with a higher fraction of around 15%, most of which indicates individual mobility from neighboring areas such as Anatolia and the North Caucasus, and a smaller fraction explained by trans-Eurasian mobility networks. However, these figures cannot be extrapolated to the entire area of Georgia, as our sampling concentrates on Iberian urban centers such as Mtskheta (Samtavro) and Tbilisi and their surroundings. The Greek Historian Strabo refers in his Geography (Book 11, Chapter 3) to this area as the confluence of the Aragvi River’s two branches - the Dariel Pass’s main route - which since antiquity has been considered a crucial passageway controlling the migration on either side. Although generally known as a rural area, it has the earliest evidence of Christianization and served as a defensive system for the capital. Our analyses combining haplotype sharing and ancestry modeling revealed a mixed social organization with rural endogamic communities together with genetically more diverse communities of unrelated individuals in urban sites, including the ’ancestry outliers’ buried according to the same rites as the ‘locals’ (see **SI Material 1**). Notably, the individuals with a high Anatolian/Levantine ancestry component were buried in the urban necropolis of Samtavro and mostly date to the 5^th^ century AD, which is about a century after Christianity became Iberia’s official religion.

Finally, we generated new insights into the practice of ACD in the Early Medieval Caucasus. A recent study using geometric morphometric analysis between medieval Hungary and Georgia concluded that ACD was not associated with the Huns but rather with the Sarmatians and Alans in the North Caucasus ^40^. Furthermore, due to the lack of craniometric data from infants and children with ACD (only one 7 to 9-year-old child from Samshvilde), it has not been possible to establish that such individuals were born into the community. Also, burials of individuals with ACD did not indicate sex bias, higher status, nobility, or poorer health status ^52^. Here, we show not only the local ancestry of the majority of ACD cases but also their biological relatedness, establishing that ACD has been practiced locally and was passed on to the next generation. However, a person with ACD was more likely to have migrated - directly or through recent ancestors - from across the Eurasian steppe. This observation parallels the findings from a study on Early Medieval deformed skulls from Bavaria, whose collective ancestry derived from Eastern Europe ^26^. Here, our modeling analyses of the ACD in Georgia support the hypothesis of Sarmatian and Alan origin in some cases, but with haplotype sharing we identify direct genealogical links to individuals from the Avar and Hun periods in the Carpathian basin. Remarkably, all the outlying ACD individuals, with the exception of the possibly contaminated SVL005, had their deformation carried in a different way (‘tabular’) than the one associated with the Huns (‘annular’) ^53^.

### Limitations

In this study, we demonstrate the efficacy of dense sampling in detecting subtle and transient admixture events, such as the elevated genetic affinity to Anatolia of some individuals following the Christianization of the Iberia Kingdom. However, the genetic landscape in the country’s west, corresponding to ancient Colchis, remains only resolved on a broad level due to sample underrepresentation. Our analyses do not detect significant gene flow from the Aegean region despite the enduring presence of Greek colonies along the Black Sea coast. Future research in this specific region is poised to elucidate if and how historical narratives are mirrored in population genetics.

## Supplementary Figures

**Supplementary Figure 1.**
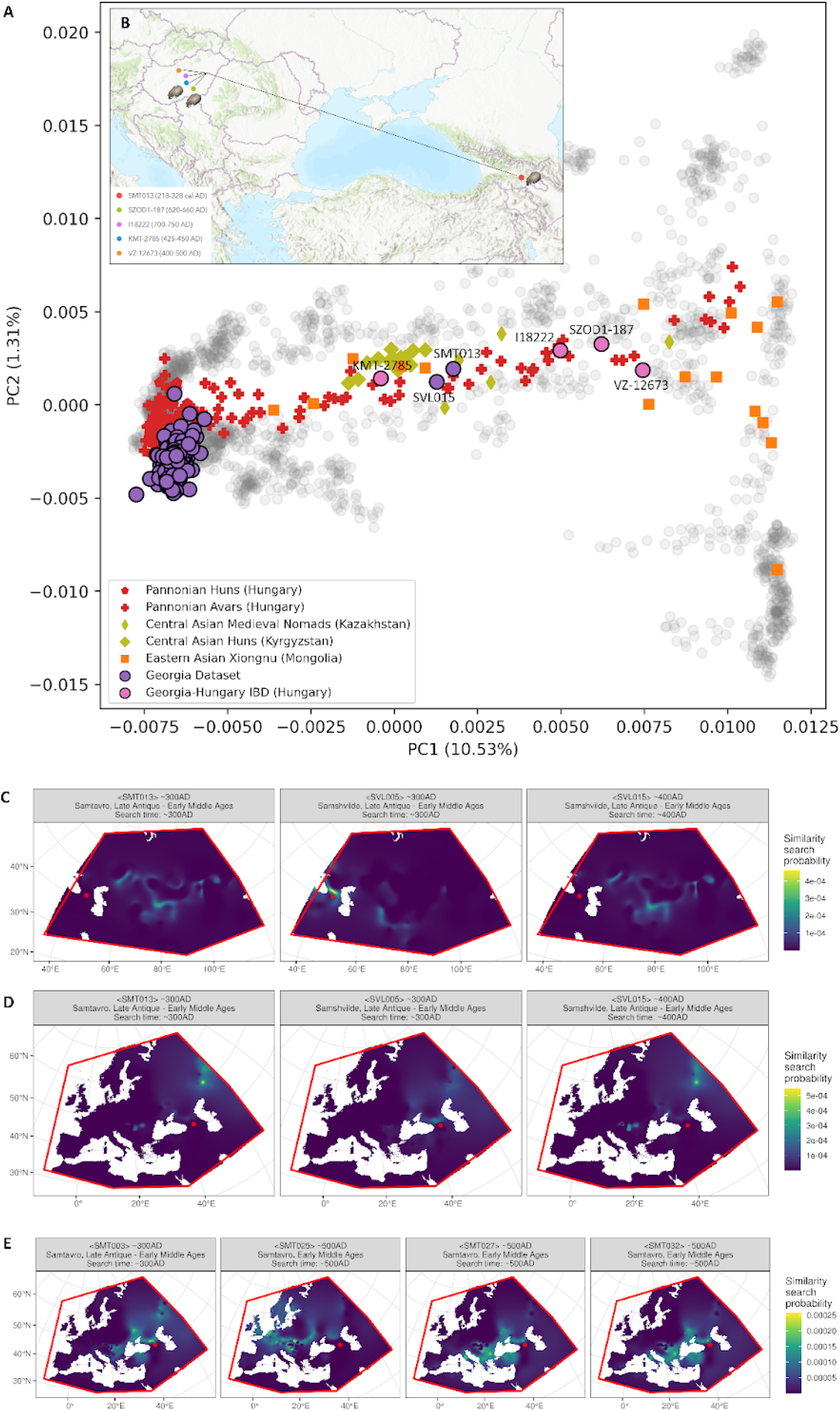
**A.** PCA on 2260 present-day individuals from entire Eurasia. We project 218 individuals from this study using the ‘*lsqproject*’ parameter of smartpca and annotate the ancestry outlier. For context, we project other relevant ancient genomes from Hungary, Central Asia, and Mongolia. Four published individuals from Hungary who share IBD with SMT013 are also annotated. **B.** Map of the IBD connections (≥12cM) between SMT013 and four Hungary individuals. Individuals with artificial cranial deformations are annotated with respective icons. **C.** Mobest search results of SMT013, SVL005, SVL015 centered on Central Asia. **D.** Mobest search results of SMT013, SVL005, SVL015 centered on Western Eurasia. **E.** Mobest search results of SMT003, SMT025, SMT027, SMT032 centered on Western Eurasia.

**Supplementary Figure 2.**
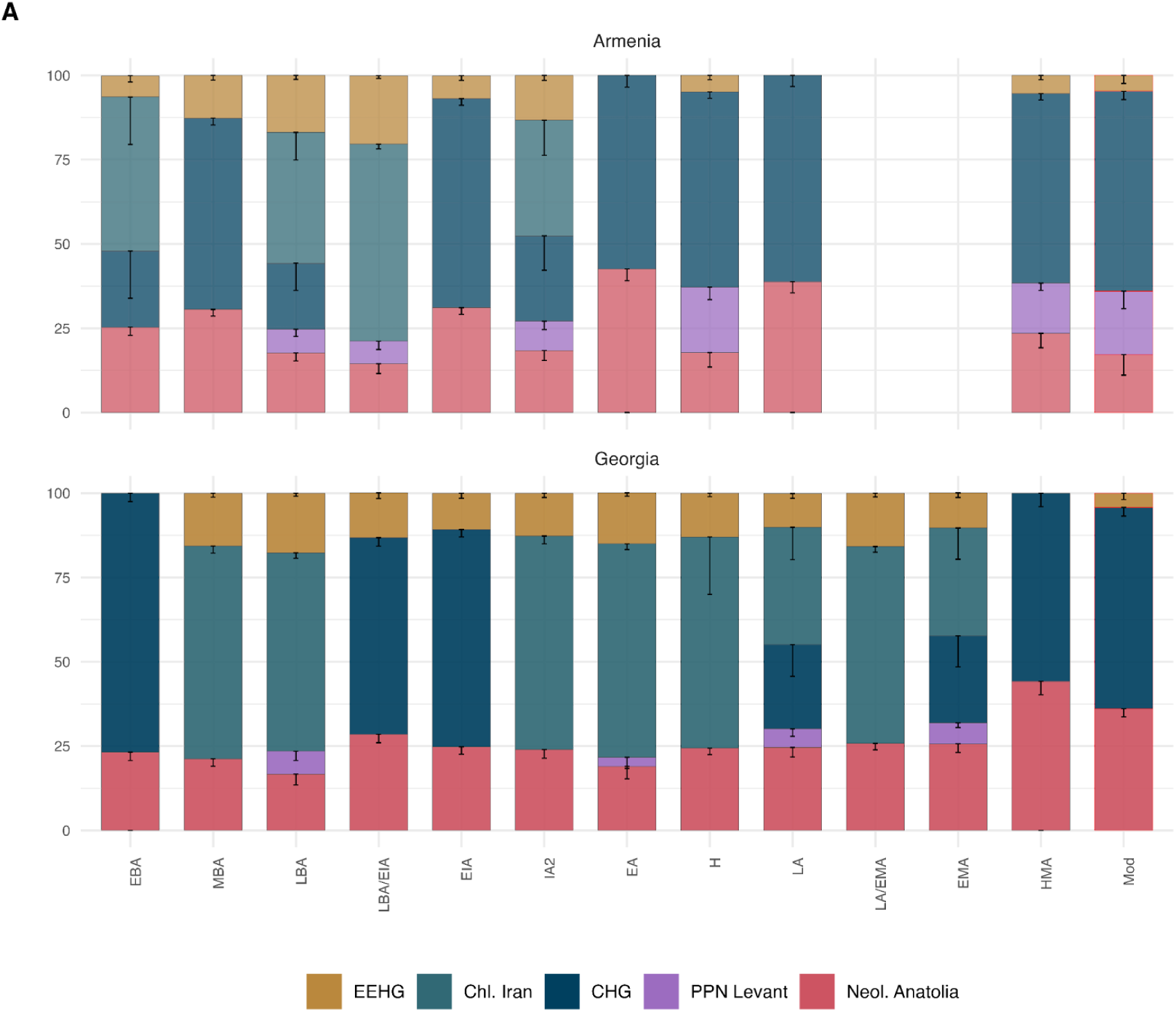
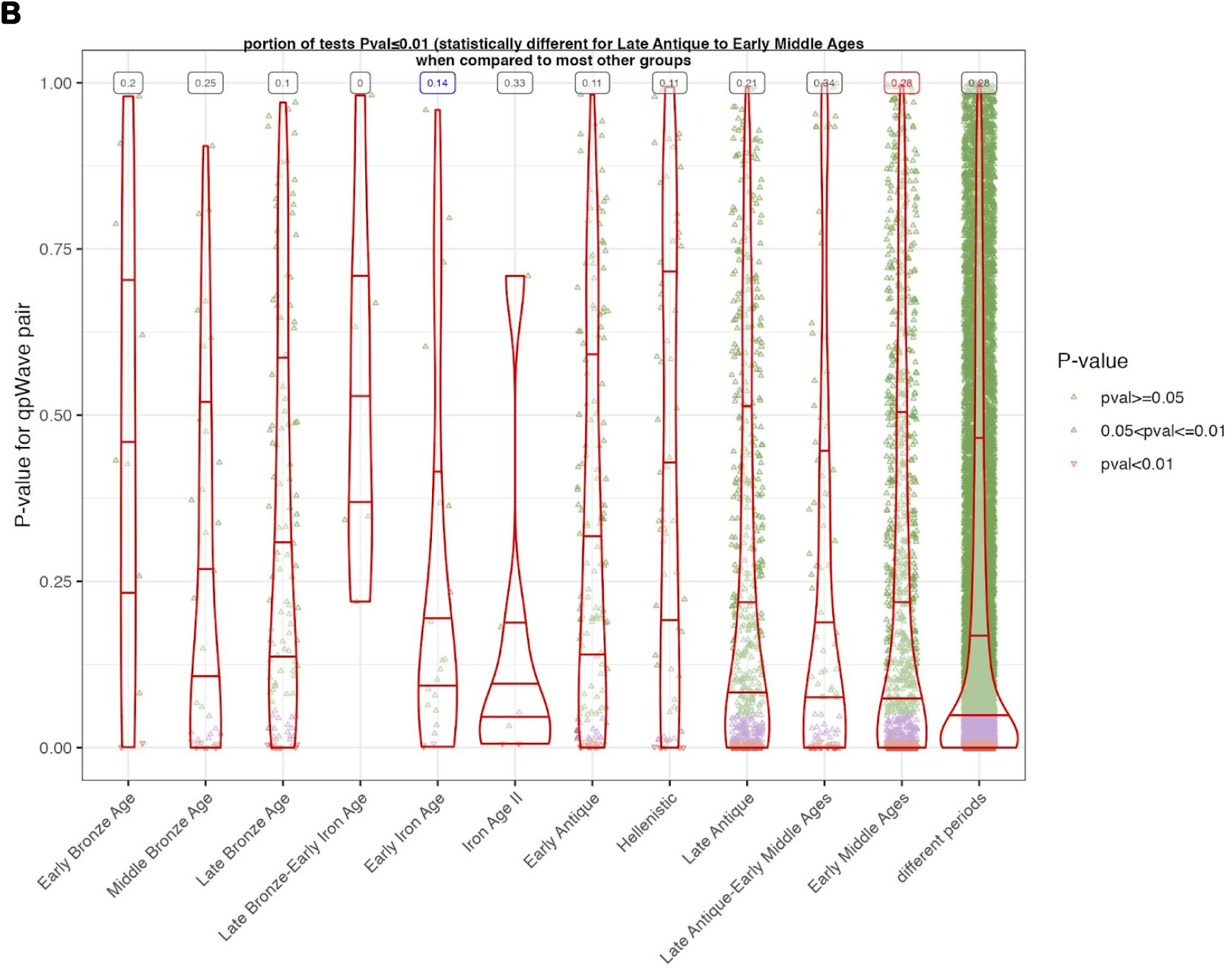
A. Distal *qpAdm* models for Georgian and Armenian groups through time. Estimated coefficients are plotted with -1SE. A red borderline is applied on the bar plots of models with low p-values. **B. Violin plots of pairwise *qpWave* for all periods with more than two individuals.** Ancestry outliers or individuals with <100,000 SNPs covered are excluded from the pair combinations. The portion of tests with p-value < 0.01 - a rough measure of genetic variability through time – is given at the top of every violin plot. Pairwise comparisons for proportions were performed with the R stats package. The p-values were adjusted using the embedded Bonferroni correction. The high proportion in the group ‘Late Antique – Early Middle Ages is statistically significant for many of the other groups, especially the earlier ones

**Supplementary Figure 3.**
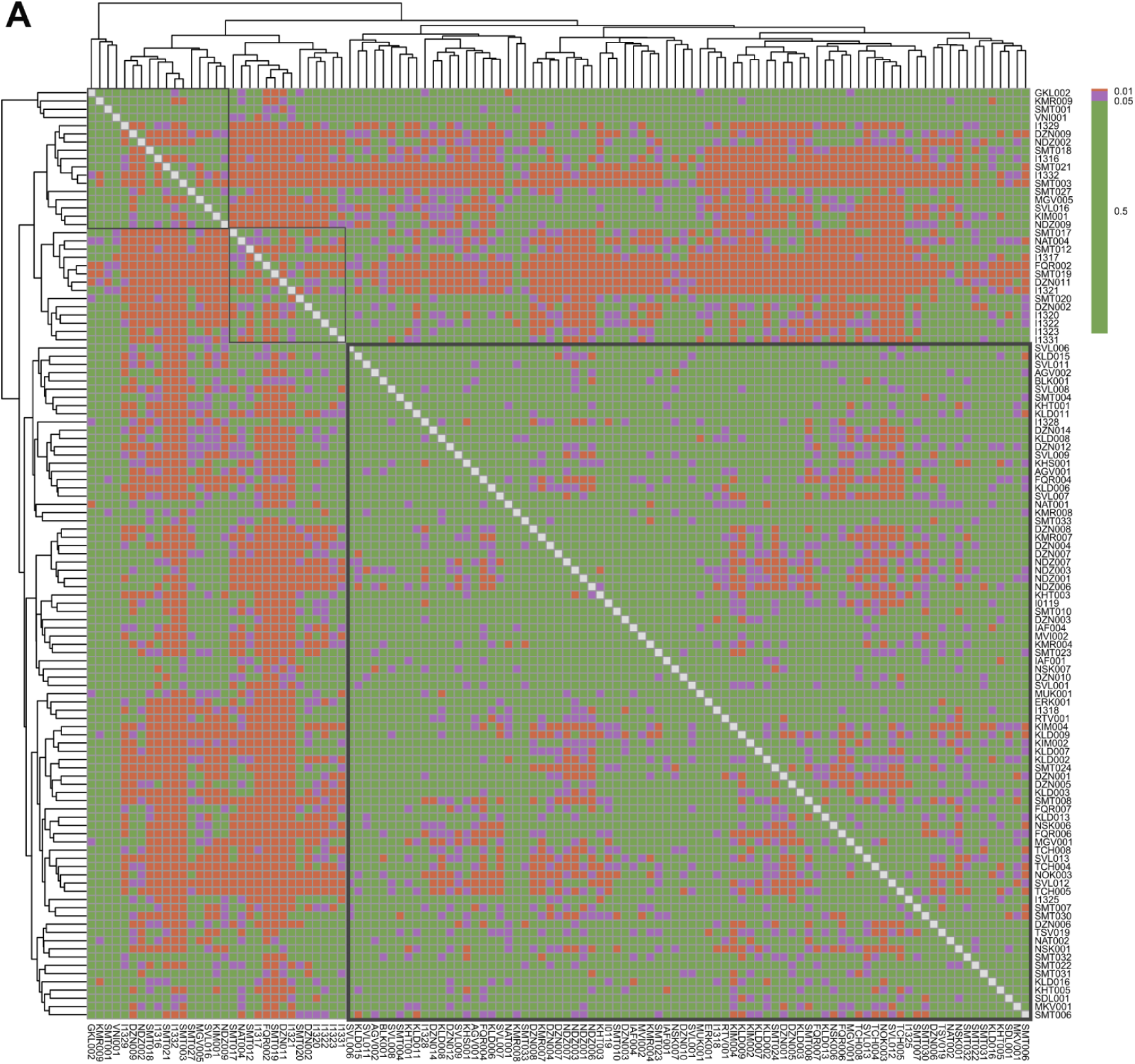
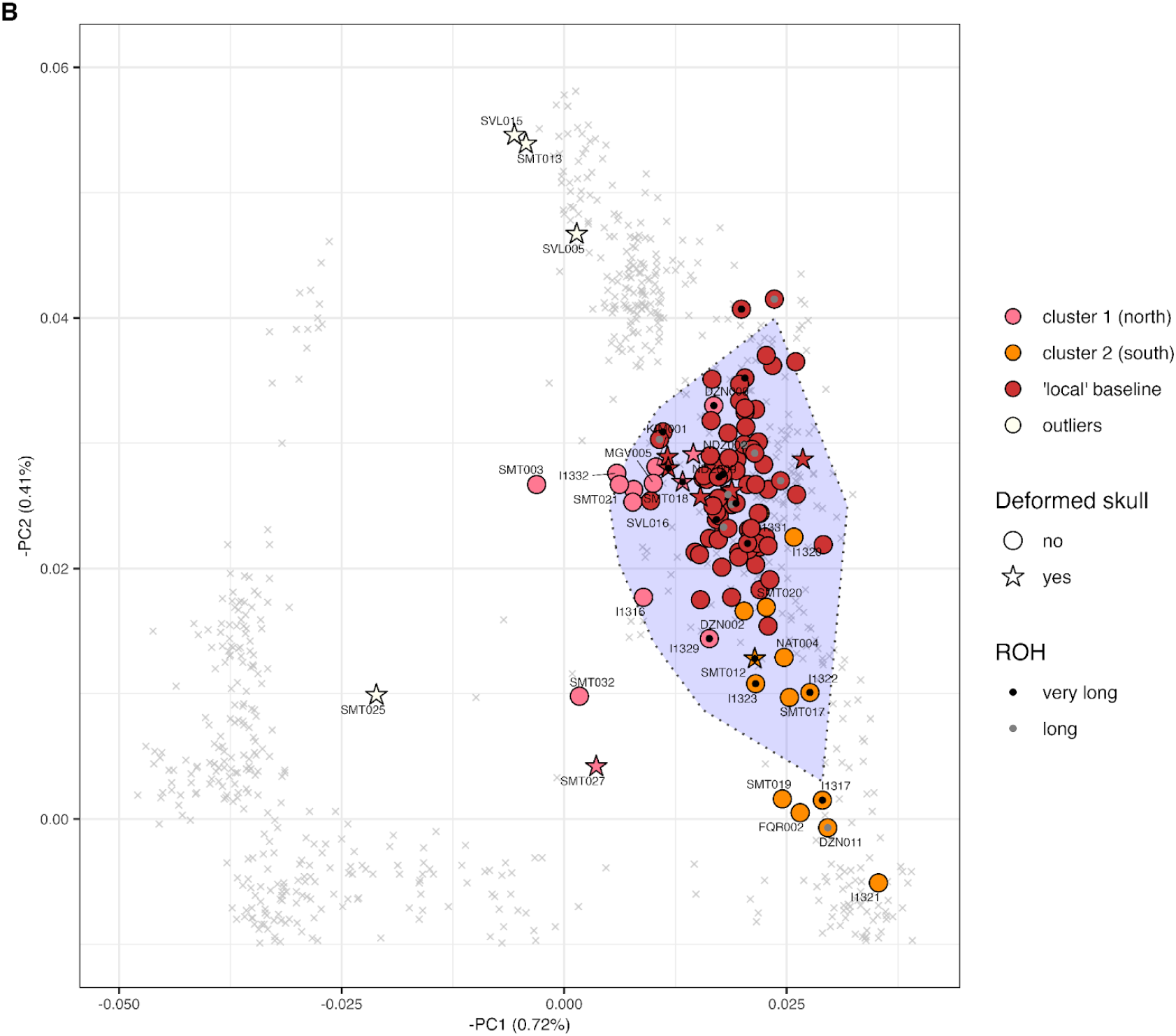
**A.** Clustered heatmap of the *qpWave* p-values for individuals from the Hellenistic period to the Early Middle Ages (HLAEMA). For visual convenience, we color-coded p-values according to the three value ranges used to interpret the tests. Three main clusters are formed, as indicated by the squared outlines. **B.** Zoom in of West Eurasian PCA focused on individuals from the Hellenistic - Early Middle Ages combined with information about *qpWave* clustering, ROH, and ACD. The range of ‘native’ South Caucasus ancestry is based on published Bronze Age data.

**Supplementary Figure 4.**
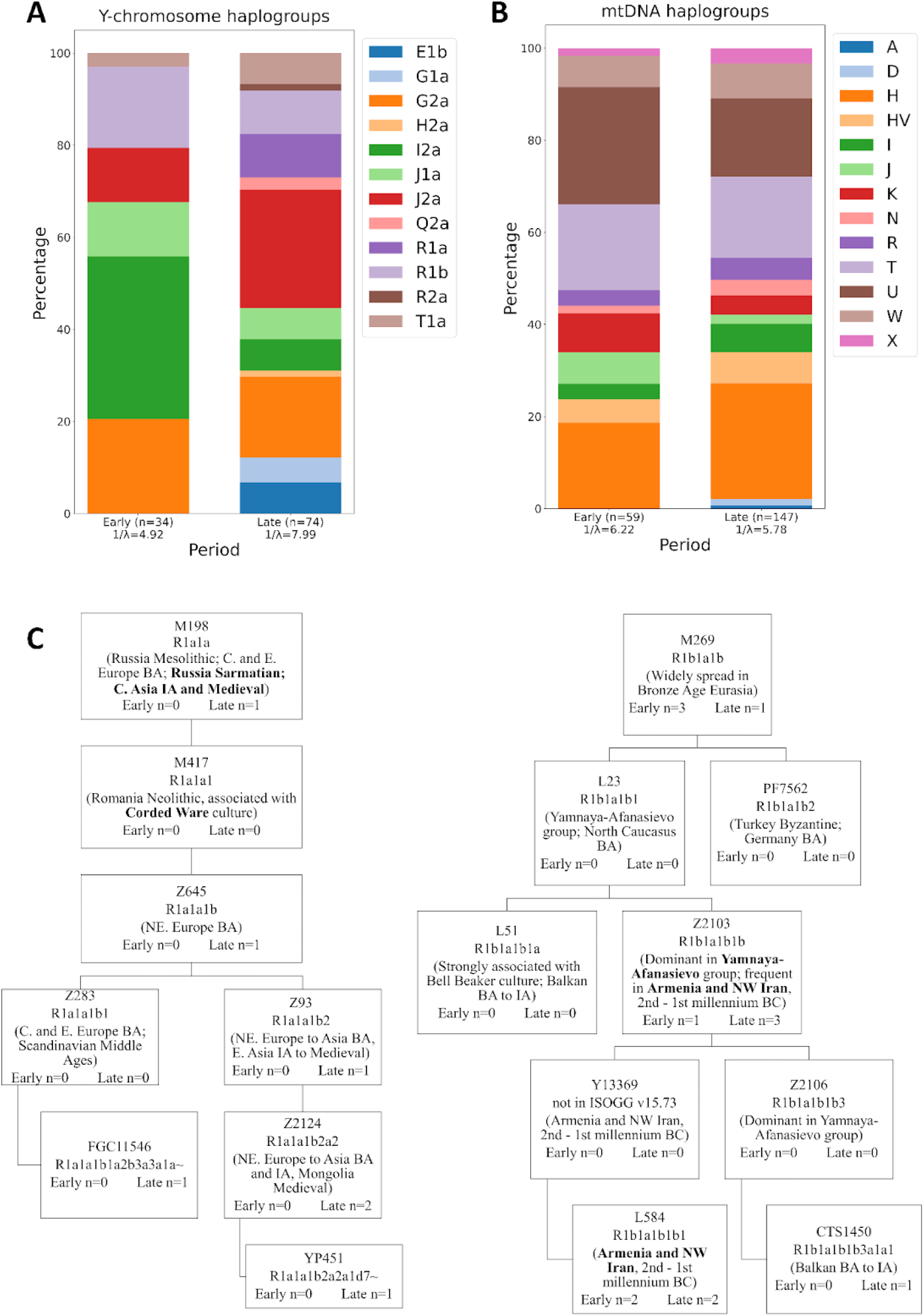
**A**. Comparison of Y-chromosome haplogroup distribution in early periods (Early Bronze Age to Iron Age II) and late periods (Early Antique to Early Middle Ages) of the time transect. The abbreviation 1/λ denotes the inverse Simpson index, a diversity measure also known as the “effective number of types” (calculated as described in **Methods**). **B.** Comparison of mitochondrial haplogroup distribution in early and late periods (as in panel a). **C.** Y-chromosome phylogenetic tree of R1a1a (R1a-M198) and R1b1a1b (R1b-M269) with temporal and spatial distribution of selected sublineages relevant for our dataset.

**Supplementary Figure 5.**
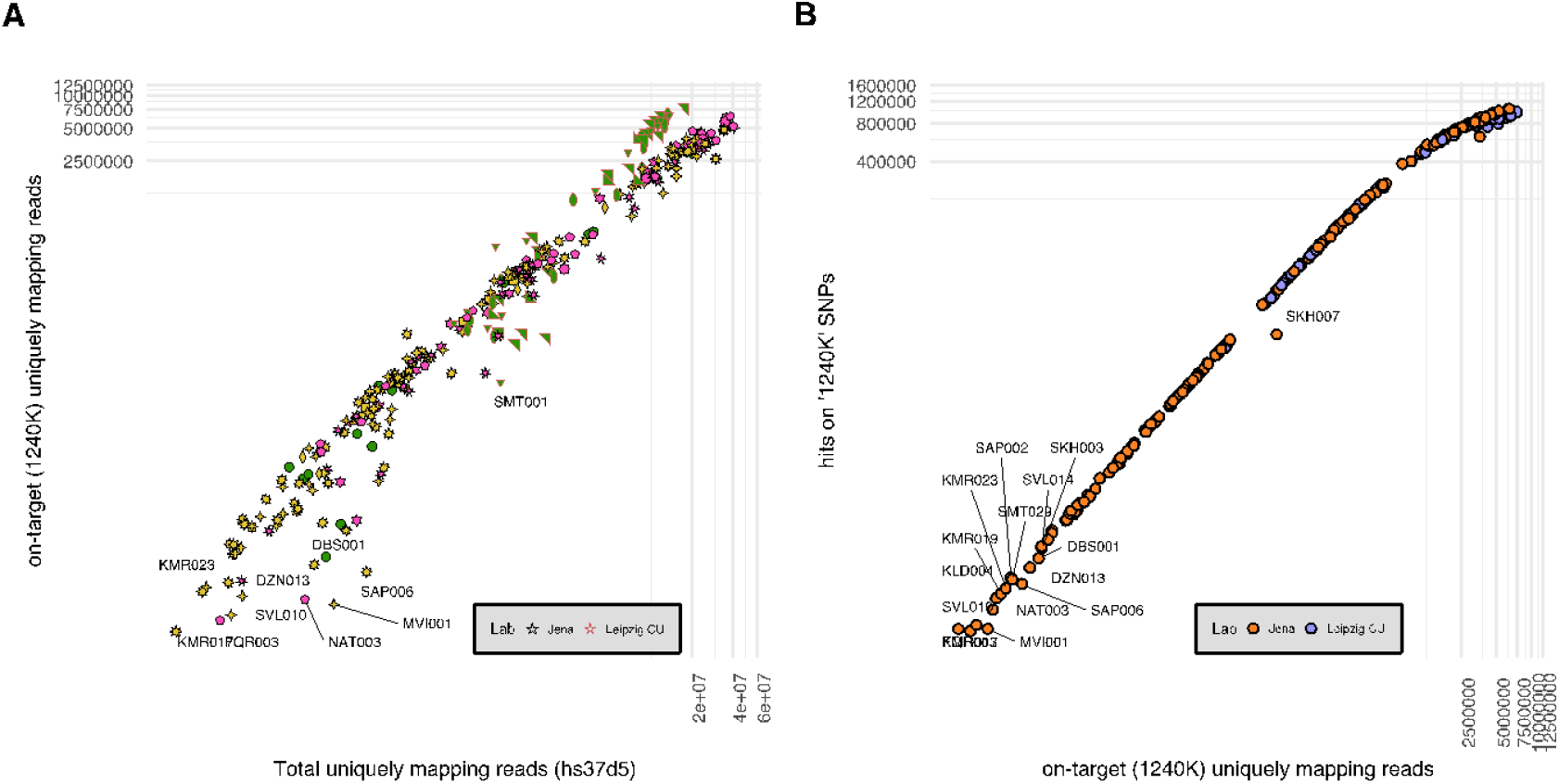
Hybridization capture (‘1240k’) efficiency and possible batch effects for genomic libraries processed in Germany. **A.** A scatterplot of unique mapping reads onto the reference genome after quality filtering steps against unique mapping reads to 1240k positions. It indicates comparable capture performance across different batches (annotated in distinct color shapes). Libraries that deviate belong to various batches and the lower range of human-mapped reads. **B**. As different probes are pooled to create 1240k array capture, this scatterplot evaluates if the coverage on 1240k positions scales up with an increase in on-target endogenous. In both plots, a log10 transformation is applied on both axes.

## Materials and Methods

### Sampling of osteological material

This study started as a pilot project to evaluate ancient DNA preservation in the osteological collections at the Anthropological Laboratory (Ivane Javakhishvili Institute of History and Ethnology). A total of 67 skeletal samples (25 petrous portions and 42 teeth) spanning from the Late Bronze Age to the Early Middle Ages were initially collected from the collection in 2021. After the processing of the batch in the lab facilities of the Department of Archaeogenetics of the Max Planck Institute for Evolutionary Anthropology (MPI-EVA), the ancient DNA molecules were determined to carry characteristic terminal deamination and to be preserved in rates from 0.1% to 67% for 61/67 of the samples (median 27%) (see ‘Sequence data processing and quality control’). Based on empirical evidence, the pilot study indicated that 90% of the samples could produce analyzable ancient DNA data after applying hybridization capture (see ‘DNA extraction and data production’). We then scaled the study up to 256 samples from the Anthropological Laboratory and 76 samples from collections of the Museum of History of Georgian Medicine, which were collected during three sampling phases in 2021-2022. In all cases, we collected the material by applying -when necessary-thermal weakening of glue based on wax and/or mechanical extraction of the temporal/petrous bone from the skull and the teeth from the mandibles/maxillas. Once the samples reached the ancient dedicated clean room facility at the Max Planck Institute of Geoanthropology (formerly MPI for the Science of Human History) in Jena, Germany, they were subject to surface decontamination strategies (UV irradiation) and were sampled using a protocol described here https://doi.org/10.17504/protocols.io.bqd8ms9w, based on a minimally invasive method for the petrous portion ^54^ and a protocol with a cut at the cementoenamel junction of the teeth and drilling from the pulp chamber that maximizes recovery of both host and pathogenic DNA (https://doi.org/10.17504/protocols.io.bqebmtan. An average of c. 50mg of bone powder was obtained per sample.

Material from 15 samples from the Late Antique-Early Medieval site of *Samtavro* was collected during a different sampling trip in 2013 at the Anthropological Laboratory and subsequently processed in the ancient DNA facilities of the Australian Centre for Ancient DNA, The University of Adelaide, Australia, and the Department of Genetics, Harvard Medical School, Boston, Massachusetts.

### C14 dating

For 98 samples, we generated radiocarbon (^14^C) dates using a MICADAS-type Accelerator Mass Spectrometry system of the Curt-Engehorn-Center Archaeometry in Mainz (Table S10). Collagen was extracted from 0.16-3gr of a sample from the same skeletal element used to produce ancient DNA, purified by ultrafiltration, and freeze-dried. A glue-decontamination step was applied beforehand for specified samples. The isotopic ratios ^14^C/^12^C and ^13^C/^12^C of the samples, blanks, control standards, and calibration standard (Oxalic Acid-II) were all measured simultaneously in the AMS. The ^14^C ages were normalized to ^13^C=-25‰ ^55^ and were calibrated using the dataset *IntCal20* and software *Oxcal* ^56^. In their majority, samples were radiocarbon-dated post-data production, and the selection strategy was as follows: 1. At least one ^14^C date for every sampled phase per site, 2. all genetic ‘outliers’, 3. All individuals with artificial cranial deformations, and 4. additional samples from sites where a previous radiocarbon date widely disagreed with archaeological dating.

### DNA extraction and data production

The steps from the DNA extraction to the library preparation and the hybridization capture and sequencing were performed in three different facilities with dedicated clean rooms: the Core Unit (CU) of MPI-EVA in Leipzig, the facility of Department Archaeogenetics (MPI-EVA) at the MPI of Geoanthropology (formerly MPI for the Science of Human History) in Jena, Germany, and the Department of Genetics, Harvard Medical School, Boston, Massachusetts.

### Germany (Core Unit of MPI-EVA, Department of Archaeogenetics – Jena)

DNA was extracted using a silica-based method that is optimized for the recovery of short DNA fragments ^57^ and following a modified version of a published protocol (https://doi.org/10.17504/protocols.io.baksicwe). First, bone powder samples were submitted to the decalcification and denaturation step, adding 1ml of extraction lysis buffer (0.45 M EDTA, pH 8.0, 0.25 mg/ml proteinase K, 0.05% Tween-20). From the resulting lysate 125-150μl were aliquoted and purified using an automated handling system (Bravo NGS Workstation B, Agilent Technologies, with silica-coated magnetic beads and binding buffer D as previously described ^58^. The final elution volume was 30μl, and the entire product was used for the library preparation. The remaining lysate was freeze-stored (-25°C) and re-aliquoted as described if more than one DNA libraries were requested per sample. An extraction blank was included with each extraction plate. An automated version of the single-stranded DNA library preparation ^59^ was applied to all DNA following a published protocol ^60^. The library preparation plate included the extraction blank as well as additional negative controls (library blanks). Library yields and the efficiency of library preparation were determined using one quantitative PCR assays ^60^. The libraries were amplified to plateau by running 35 PCR cycles and tagged with pairs of library-specific indices using AccuPrime Pfx DNA polymerase ^60^. Amplified libraries were purified using SPRI technology ^61^ as described in Gansauge et al. (2020). Libraries were enriched using a bait set targeting approximately 1.2 M SNP positions (in-solution hybridization capture) ^32,62,63^. Libraries that were prepared at the Core Unit were subject to two consecutive rounds of capture as described in Fu et al. (2013). Libraries from the first phase of sampling (pilot batch) were initially sequenced at ∼5-10M reads on a HiSeq4000 with 75 bp single-end sequencing chemistry (1 x 76 + 8 + 8 cycles). Following the evaluation of the shotgun metagenomic sequencing data for the endogenous DNA and damage patterns (see Sequence data processing and quality control), a selected number of libraries were submitted to hybridization capture after which they were sequenced again. For the remaining sampling phases, produced libraries were sequenced only after the enrichment with in-solution capture. Post-capture, a total of 376 libraries were sequenced on a HiSeq4000 with 75 bp single or paired-end sequencing chemistry (1/2 x 76 + 8 + 8 cycles) for a total of ∼15-140M reads depending on their complexity and a case-by-case evaluation.

### Boston

#### Sequence data processing and quality control

Demultiplexed sequencing data were processed with *leeHom* [v1.1.5] ^64^ for merging of paired-end data and trimming of Illumina adaptor sequences from reads shorter than 76bp. Forward, reverse, and merged reads were mapped against the human reference hs37d5 with the Burrows–Wheeler aligner (BWA; v.0.7.12) ^65^with parameters -n 0.01 -o 2 -l 16500 (disabled seeding).

The output files in Binary Alignment Map (BAM) format were subsequently processed with nf-core/eager nextflow pipeline [v2.4.6] ^66,67^ using the custom configuration file from MPI-EVA (https://github.com/nf-core/configs/blob/master/docs/eva.md). In more detail, running parameters were set as follows: run_bam_filtering = true, bam_mapping_quality_threshold = 0, bam_unmapped_type = ’bam’, bam_filter_minreadlength = 30, run_mtnucratio = true, damage_calculation_tool = ’*mapdamage*’, mapdamage_downsample = 100000, bamutils_clip_single_stranded_none_udg, genotyping_tool = ’*pileupcaller*’, pileupcaller_min_map_quality = 25, pileupcaller_min_base_quality = 30, run_sexdeterrmine = true, run_nuclear_contamination = true. Bam files from same-sample/individual libraries were processed separately for quality control estimates like male contamination and deamination rates but were merged, sorted, and deduplicated before sex determination, samtools mpileup, and finally extraction of pseudo-haploid genotypes with pileupCaller (https://github.com/stschiff/sequenceTools). To evaluate the contamination status of an individual/library, three additional methods were applied on bam files with mapping quality 30: *hapCon* ^68^, *Schmutzi* (Renaud et al., 2015), and *AuthentiCT* ^69^. By using a haplotype copying approach related to imputation, *hapCon* estimates contamination on ancient DNA data of male individuals with a lower coverage threshold and less uncertainty than similar male contamination tools like *ANGSD* ^70^ (embedded in nf-core/eager). *Schmutzi* estimates sample contamination from mitochondrial reads, but estimates were shown to be unreliable for Mt/Nu ratio>200 ^71^. *AuthentiCT* harnesses information from the damage distribution and provides an alternative when the other three methods are not accessible. However, we noticed a negative correlation between *AuthentiCT* estimates and deamination rates, while the latter is also known to correlate with sample age {Dabney, Jesse et al, 2013; Lindahl, Tomas, 1993}. With a timespan from a few hundred to 5000 years, this correlation is shown in our dataset as well (Table S1). Therefore, when we were relying only on *AuthentiCT*, we marked as ‘include_contaminated’ samples with a high contamination value (≥10%) and terminal 5’ deamination rate of 30% or lower (Table S1). In the same way, we also flagged libraries/individuals with reliable contamination estimation from the other methods when the upper boundary was close to 10%. In addition, for a potential XXY (Klinefelter; TSV006), we ignored the high contamination rate with *hapCon*, but we also marked it as ‘include_contaminated’ because of *AuthentiCT*. In total, we sorted 21 cases on this status, which we maintained in our dataset and distinctly annotated them in analyses such as PCA, and pairwise *qpWave* matrix (see ‘Principal Component Analysis’ and ‘Admixtools’), but excluded them from group-based analyses. Furthermore, we completely excluded individuals as contaminated, for which the SNP coverage on 1250k positions did not exceed 60,000 (c. 5%). A hundred twelve (112) libraries were completely excluded from our dataset as of low quality with less than 25,000 SNPs covered and contamination rates either very high or impossible to estimate. Finally, as part of the quality control, we analyzed all pairs for close relatedness (see ‘Close biological relatedness’), including previously published data from Georgia ^41^, and determined 12 cases of identical individuals/twins. With no definitive evidence of twins (e.g., same skeletal element), we concluded that these were all cases of sample mix-ups in the bone collection or duplicates created while registering samples with very similar archaeological IDs in the ancient DNA lab. However, in three cases of identical pairs, the samples came from different sites and it has not been possible to ascertain the original location. We present and make available the data of these individuals making the distinction of ‘unresolved_origin’. Accordingly, we provide the data from two individuals from the site of Tsaishi but exclude them from group or temporally ordered analyses, since two radiocarbon dates and IBD analyses on three different samples from the same site indicated a mixed layer with material from the Iron Age and the high Middle Ages.

With approximately 30% of the libraries excluded due to very low SNP coverage, we investigated the possibility of batch effects in the efficiency of the hybridization protocol. We evaluated the complexity of libraries post-capture by looking into the number of the uniquely mapping reads on the SNP positions against all the uniquely mapping reads on the human reference (**Supplementary Figure 5**). For successful capture essays, these two numbers are expected to be highly correlated with a relatively fixed coefficient across different capture plates/batches. We noticed that the few libraries that deviated from this expectation a. belonged to different capture plates and b. had ≤10,000 unique reads mapped on hs37d5, suggesting outlying cases of libraries with low preservation of DNA rather than problems during hybridization.

Furthermore, we also noted that libraries that were submitted for two rounds of capture (Leipzig CU) on average yielded more SNPs (covered at least once) than the one-round captures performed in Jena. We caution, however, that the current data between the two labs are not comparable to establish whether libraries at the lower range of preservation would have yielded a higher number of SNPs after two rounds of capture and 20-40M reads instead of one round and sequencing at a higher depth (i.e., ≥40).

### Data compilation

We present a dataset of genome-wide data from 219 individuals from present-day Georgia that includes a total of 26 individuals flagged as follows: ‘possibly contaminated’ (n=21), ‘unresolved origin’ (n=3), and ‘uncertain date’ (n=2). We used the Poseidon framework (https://www.poseidon-adna.org/#/) ^72^ to combine the per-individual genotype outputs generated by nf-core/eager into one eigenstrat file with an associated annotation file. We manually enriched the annotation file, filling in information about biological relatedness, contamination rates estimated by all the methods applied herein, and grouping assignments. We applied the same steps to a recently published dataset that included eight individuals from Georgia ^41^. More specifically, downloaded bam files from European Nucleotide Archive (ENA) were processed with nf-core/eager under the same configuration as for the main dataset, applying bamutils_clip_double_stranded_none_udg_right = 10), after which a Poseidon package was created. Transversions-only versions of the genotypes were created and examined with Principal Component Analysis (see below), which suggested that residual damage might introduce biases in some of the downstream comparative analyses. Therefore, we co-analyzed only the individuals with enough data on transversions (≥20,000; geo015_tv, gur017_tv) in PCA and Admixtools. In addition, we excluded individual geo29 from the Bronze Age site of Didnauri, for which we processed a different sample with c. 900K SNPs. The two datasets from Georgia were merged with the Allen Ancient DNA Resource (AADR) [v54.1] ^73,74^ and a recently published dataset from the Aegean ^75^. In total, approximately 250 individuals from Armenia, Azerbaijan, Russia, and Eastern Turkey post-dating the Neolithic period ^6,28,29,41,42,51,76^ were of high relevance to our study and were included in our analyses.

We have applied two grouping systems for the downstream analyses: the site-period-specific (‘Site_Period’) and the broader period-specific (‘Georgia_Period’). In both cases, information about contamination, relatedness, and ancestry outliers is appended with a corresponding suffix (e.g., ‘_contam’, ‘_rel’, etc.). Regarding the period assignment, we apply 12 consecutive and transitional periods in Georgian prehistory and history [Early Bronze Age (n=7), Middle Bronze Age (19^th^-16^th^ c. BA; n=8), Late Bronze Age (15^th^-13^th^ c. BC; n=21), Late Bronze Age-Early Iron Age (13^th^-12^th^ c. BC; n=6), Early Iron Age (12^th^-9^th^ c. BC; n=5), Iron Age II (8^th^-7^th^ c. BC; n=9), Early Antique (6^th^-4^th^ c. BC; n=16), Hellenistic (3^rd^-1^st^ c. BC; n=12), Late Antique (1^st^ c. BC-3^rd^ c. AD; n=35), Late Antique-Early Middle Ages (3^rd^-4^th^ c. AD; n=9), Early Middle Ages (4^th^-10^th^ c. AD; n=48), and High Middle Ages (11^th^-15^th^ c. AD; n=2)]. In the course of radiocarbon dating, some discrepancies in the range of ±200-300 years with the archaeological dating – which is mostly based on burial goods - arose. Whenever possible, radiocarbon dating, relatedness information from IBD analyses, and stratigraphic association of burials were combined to update the period assignments for burials without ^14^C date information.

#### Close biological relatedness

To estimate close genetic relatedness among all ancient individuals from Georgia [current study and Koptekin et al. (2023)], we calculated the pairwise mismatch rate (PMR) from n overlapping SNPs in the autosomal portion of the eigenstrat file. Accordingly, standard errors were output as √pmr(1-pmr)/nSNPs. The median of c. 0.25 was calculated from all c. 25k pairs and served as a point estimate of the baseline of unrelatedness in our dataset. The expected values for twins/identical, first-degree, and second-degree relatives were defined accordingly (i.e., 0.5*0.25, 0.75*0.25, and 0.875*0.25, respectively). With Z-scores estimated as 0.25-pmr/sterr, we assessed the significance of the first and second-degree assignments. We further validated our results with BreadR (https://github.com/cran/BREADR) ^77^, which also estimates PMR after thinning to remove possible effects from LD. Assuming a binomial distribution for PMR values, every pair is returned with a posterior probability distribution for different degrees of relatedness.

#### Principal Component Analysis (PCA)

We computed PCA on three different datasets of present-day populations using smartpca [v1600] from EIGENSOFT [v7.2.1] ^78,79^ with parameters *lsqproject*: YES and *shrinkmode*: YES. The first dataset (1300 individuals) comprised modern populations from Western Eurasia genotyped on the Human Origins panel ^43,80–82^, with additional populations from the Caucasus genotyped at the same panel ^83^: Abazin.HO Armenian_Hemsheni.HO, Avar.HO, Azeri.HO, Circassian.HO, Darginian.HO, Ezid.HO, Ingushian.HO, Kabardinian.HO, Kaitag.HO, Karachai.HO, Kubachinian.HO, Kurd.HO, Lak.HO, Tabasaran.HO, Turkish_Balikesir.HO. The second dataset (1486 individuals) included the populations from the first dataset and the following populations from C. Asia which were included in the same studies, and ^84^: Balochi.HO, Brahui.HO, GujaratiA.HO, GujaratiB.HO, GujaratiC.HO, GujaratiD.HO, Iran_Zoroastrian.HO, Kalash.HO, Makrani.HO, Moldavian.HO, Pathan.HO, Sindhi_Pakistan.HO, Tajik.HO. Last, the third dataset (2260 individuals) aimed to represent the entire Eurasia, therefore groups -some of which were published in ^85,86^- were added to the population list: Aleut.HO, Altaian.HO, Altaian_Chelkan.HO, Ami.HO, Atayal.HO, Bashkir.HO, Besermyan.HO, Buryat.HO, Cambodian.HO, Chukchi.HO, Chuvash.HO, Dai.HO, Daur.HO, Dolgan.HO, Dungan.HO, Enets.HO,, Eskimo_ChaplinSireniki.HO Eskimo_Naukan.HO, Even.HO, Evenk_FarEast.HO, Evenk_Transbaikal.HO, Gagauz.HO, Han.HO, Hezhen.HO, Itelmen.HO Japanese.HO, Kalmyk.HO, Karakalpak.HO, Karelian.HO, Kazakh.HO, Ket.HO, Khakass.HO, Khakass_Kachin.HO, Khamnegan.HO, Kinh.HO, Korean.HO, Koryak.HO, Kubachinianv, Kyrgyz_Tajikistan.HO, Kyrgyz_Kyrgyzstan.H, Kyrgyz_China.HO, China_Lahu.HO, Mansi.HO, Miao.HO, Moldavia.HO, Mongol.HO, Mongola.HO, Nanai.HO, Naxi.HO, Negidal.HO, Nganasan.HO, Nivh.HO, Nogai_Stavropol.HO, Nogai_Karachay_Cherkessia.HO, Oroqen.HO, Russian_Archangelsk_Krasnoborsky.HO, Russian_Archangelsk_Leshukonsky.HO, Russian_Archangelsk_Pinezhsky.HO, Saami.HO, Selkup.HO, She.HO, Sherpa.HO, Shor_Khakassia.HO, Shor_Mountain.HO, Tatar_Kazan.HO, Tatar_Mishar.HO, Tatar_Siberian.HO, Tatar_Siberian_Zabolotniye.HO, Thai.HO, Tibetan.HO, Todzin.HO, Tofalar.HO, Tu.HO, Tubalar.HO, Tujia.HO, Turkish_Balikesir.HO, Turkmen.HO, Tuvinian.HO, Udmurt.HO, Ukrainian_North.HO, Ulchi.HO, Uzbek.HO, Veps.HO, Xibo.HO, Yakut.HO, Yi.HO, Yukagir_Tundra. All population names are reported as in AADR [v54.1]^73,74^.

#### Mobest

We implemented the Mobest [v1.0.0] ^87^ tool to perform spatiotemporal interpolation of human genetic ancestry and probabilistic similarity search of those individuals identified as genetic outliers. For the individuals labeled SMT003, SMT025, SMT027, SMT032, and SVL005, we employed Principal Component Analysis (PCA) computed on the first dataset, comprising 1300 present-day individuals. We then used PCA’s first (PC1) and second (PC2) components as genetic coordinates. We defined the research area centering around western Eurasia with a map projection utilizing the coordinate reference system EPSG: 3035. In the case of SMT013 and SVL015, a similar approach was used but was based on another PCA, this time performed on our third dataset of 2260 individuals. Again, the coordinates of PC1 and PC2 were integrated as the coordinates of an MDS analysis. In this instance, we defined the research area centering around central Asia, complemented by a map projection with the coordinate reference system EPSG: 2542. For the reference, we incorporated all the ancient samples from the current study, Koptekin et al., 2023, and those from the AADR [v54.1] database, which were situated within the bounds of our research zones.

#### Admixtools (F-statistics, qpWave, and qpAdm)

We used the modules *qpDstat*, *qpWave*, and *qpAdm* from Admixtools [v5.1] ^82^ to estimate allele sharing and model admixture events in our dataset. For *qpWave* and *qpAdm* we applied two settings of reference populations (‘right pops’): 1. Mbuti.DG, ANE (Ancestral North Eurasian; Russia_AfontovaGora3, Russia_MA1_HG.SG), Serbia_IronGates_Mesolithic, Israel_Natufian, Turkey_Epipaleolithic and Iran_GanjDareh_N, and 2. Mbuti.DG, ANE, Serbia_IronGates_Mesolithic, Levant_PPN, Turkey_N, Caucasus Hunter-Gatherers (CHG; Georgia_Kotias.SG, Georgia_Satsurblia.SG), Iran_GanjDareh_N, EEHG (Eastern European Hunter-Gatherers; Russia_EHG, Russia_Karelia_HG, Russia_Popovo_HG, Russia_Samara_HG). For the modeling of the ancestry outliers, we added group ‘China_YR_LN’ to the second setting. All the population names are reported as in the AADR [v54.1] ^74,88^. With *qpWave* we first tested that the two sets of reference populations can sufficiently differentiate the ‘left’ populations (i.e., p-value << 0.01). Because of the low-coverage genome-wide data in the group ‘Israel_Natufian’ of the first setting, we applied the parameter allsnps: YES, which uses all the SNPs available in every quadruple of f_4_-stats, instead of only the SNPs common among the populations in the entire F_4_-statistics matrix. We used the first setting to model temporal groups from Georgia and Armenia as a linear combination of ancestral -mainly Early Holocene-populations that contributed differently to the genetic make-up of Bronze Age Western Asia (distal modeling with *qpAdm* module). With the second setting, we attempted to model temporal genetic shifts since the Bronze Age period (proximal modeling). We also used it to test for cladal relationships between pairs of individuals or groups (i.e., one stream of ancestry explaining this reference set from the tested pair). Unless indicated otherwise, *qpWave* and *qpAdm* were applied to groups at the exclusion of close relatives (first to third-degree) and (possibly) contaminated individuals.

#### Runs of Homozygosity

We inferred runs of homozygosity from the genome-wide data of ancient individuals from Georgia using *hapROH* [v1.0] ^36^ using the recommended cut-off of c. >0.4x coverage on 1240k SNPs and a length cutoff of >4 centimorgan. Additionally, we also screened for long ROH on individuals at a lower coverage (c. 0.3x), which is prone to false positive rates of shorter ROH segments. We also cross-checked for consanguinity among individuals with the ‘possibly contaminated’ flag; as contamination is expected to break down ROH, individuals still exhibiting long ROH cannot carry high rates of contamination ^89^. We summarized the ROH output by classifying every individual in three ROH groups as follows: 1. background relatedness (small mating pool): ROH in the length bin 4-8cM, also individuals with a unique ROH extending up to 12cM, 2. distant parental relatedness: individuals with roh at the length bins 12-20cM and/or one ROH at 20-30cM, 3. consanguinity: all the individuals consistent with parents being related as first cousins or closer. For the latter, we also included individuals with a lower SNP cut-off at 250,000 SNPs since this was previously shown to capture cases with elevated consanguinity ^75^.

#### Imputation of ancient genomes

We used the ATLAS platform’s MLE function ^90^, specifically designed for ancient DNA, to call genotype likelihoods. For each bam file, we obtained the expected read length from the file header, which was equivalent to the sequencing cycle number. To divide the reads by their length for single-end sequencing data, we used ATLAS’s SplitMerge. Reads exceeding the expected length were grouped separately, as the mutation probability of a base due to post-mortem damage (PMD) depended on its distance from the read ends. If the DNA fragment size is greater than the sequencing length, the damage information could be ambiguous or misleading since we cannot assume that the read end aligns with the fragment end. We estimated PMD differently for reads equal to or longer than the cycle number compared to shorter reads. We estimated potential sequencing biases using ATLAS’s recal function, which considered the PMD profile from the previous steps. The recalibration patterns were built using chromosome 1 data from all the samples of one run to correct machine-specific biased calls. We performed genotype likelihood estimation on the entire SNP panel, which included about 20 million SNPs from the 1,000-genome data, via ATLAS’s call tool that considered both PMD and recal patterns. These calls were then used as inputs for imputation in *GLIMPSE* ^91^, taking the haplotypes from the 1,000-genome’s phase-3 (1KGP3) data release ^92^ as the reference.

We used *GLIMPSE* with default parameters, using HapMap phase II NCBI b37 as genetic maps. We simultaneously phased and imputed each sample, divided into per-chromosome VCFs, using the *GLIMPSE_phase* function on genomic chunks of 2,000,000 base pairs, with a buffer of 200,000 base pairs. After phasing, we concatenated the various chunks using *GLIMPSE_ligate*. To generate the final phase/imputed VCF files, with genotype posterior probabilities at each 1240k position, we utilized *GLIMPSE_sample* functions with the “solve” flag alongside bcftools [v1.3] ^93^.

#### Identical by Descent (IBD) analysis

We called Identical by Descent (IBD) segments between pairs of individuals with the tool *ancIBD* [v0.5] ^35^ using the imputed genotype likelihoods at the 1240k SNP set and the recommended cutoff for data quality (corresponding to ca. 0.75x for capture data and 0.25x for WGS data). We used the default parameters of *ancIBD*, recommended for ancient DNA data. First, we singled out cases of close and distant relatives, then we individually evaluated pairs with total IBD≥20cM for true positives by assessing date overlap in the pair, the imputed genotype likelihood scores, and the length distribution of IBDs. Finally, we visually inspected IBDs with the embedded plotting function of *ancIBD* (using the function run_plot_pair). For our measures of isolation-by-distance, we retained all pairs with IBD≥8cM and applied a threshold of imputed genotype likelihood score of >0.7 at either side of the pair.

#### DATES

We used the software *DATES* (https://github.com/MoorjaniLab/DATES_v4010) ^94^ to infer the timing of admixture on the ancestry outliers of our dataset. We ran the tool with standard parameters: binsize = 0.001, maxdist = 1 (Morgan units), and fit of decay curve from 0.45cM (lovalfit).

#### Uniparentally-inherited markers

We assigned mitochondrial haplogroups and haplotypes with Haplogrep software (v.2.1.25) ^95^, applying a quality threshold of 0.65 on >3X mitogenomes after assembling the consensus sequence (q30) with *Schmutzi*. To determine the Y chromosome haplogroups in male individuals, we first produced mpileups with samtools for 73,148 informative SNPs according to the ISOGG Y-haplotype tree from https://isogg.org/tree (Version: 15.73) and then recorded the count of derived and ancestral calls for each informative marker. We then manually inspected those counts to verify whether the presence of diagnostic SNPs for a specific haplogroup exhibited a root-to-tip trajectory or if there were anomalous jumps in the phylogeny due to residual damage and sequencing errors. We used the following formula to calculate the inverse Simpson index (1/λ): 1/λ = *N* * (*N* − 1) / ∑(*n* * (*n* − 1)), where n denotes the total number of individuals belonging to a specific haplogroup during early or late periods, and ’N’ the total number of individuals in the corresponding time period. We categorized the mitochondrial haplogroups using the one-letter scale of the ISOGG nomenclature and for Y haplogroups, we used the three-letter scale.

## Acknowledgments

This work was supported by funding from the Max Planck Society and the Max Planck-Harvard Research Center for the Archaeoscience of the ancient Mediterranean (MHAAM). We thank the members of the laboratory facilities in MPI-EVA (Core Unit) and in Jena (Department of Archaeogenetics MPI-EVA) for their support on the laboratory work. We thank the members of MHAAM, the population genetics group from MPI-EVA, and Ayshin Ghalichi, Clemens Schmid, and Guido Alberto Gnecchi Ruscone for their useful feedback.

## Notes

### Competing Interest Statement

The authors have declared no competing interest.

