## Supplementary Information Material 1 for "The Genetic History of the South Caucasus from the Bronze to the Early Middle Ages: 5000 years of genetic continuity despite high mobility"

### Material 1: Archaeological Site Descriptions

1. **Samtavro (SMT)**
2. **Tserovani (TSV)**
3. **Zhinvali (DZN)**
4. **Aragvispiri (AGV)**
5. **Nedzikhi (NDZ)**
6. **Tchiatura (TCH)**
7. **Kamarakhevi (KMR)**
8. **Nabagrebi (NBG)**
9. **Treli (TRL)**
10. **Didnauri (DDN)**
11. **Fiqris Gora (FQR)**
12. **Sapar-Kharaba (SAP)**
13. **Kiketi (KKT)**
14. **Natakhtari (NAT)**
15. **Khimshiant (KIM)**
16. **Samshvilde (SVL)**

17. **Vani (VNI)**
18. **Bazaleti (BZT)**
19. **Nastakisi (NSK)**
20. **Nokalakevi (NOK)**
21. **Khuntsi (KHT)**
22. **Atskuri (ATK)**
23. **Alaverdi (AVR)**
24. **Skhalta (SKH)**
25. **Samadlo (SDL)**
26. **Klde (KLD)**
27. **Mukhatgverdi (MUK)**
28. **Narekvavi (NRK)**
29. **Mogvtakari (MGV)**
30. **Tsaghvli (TSG)**
31. **Arboshiki (AHI)**
32. **Taribana (TRB)**
33. **Manavi (MVI)**
34. **Tsaishi (TSH)**
35. **Igoeti (IGT)**
36. **Varsimaantkari (VRS)**
37. **Erkneti (ERK)**
38. **Okrokana (OGR)**
39. **Kistauri (KTR)**
40. **Bodbe (BBD)**
41. **Bulachauri (BLK)**

**42. Mtsketijvari (MKV)**

**43. Rustavi (RTV)**

**44. Telavi (TEL)**

**45. Tetritskaro (TKA)**

**46. Lafanaantkari (IAF)**

**47. Grakliani (GKL)**

**48. Khtsisi (KHS)**

**49. Murakebi (MRA)**

**50. Ota (OTA)**

**51. Sagvarjile (SGV)**

**Please contact the corresponding author to request access to reference information on the archaeological site description.**

**Samtavro Cemetery (SMT)**

**A) General Location and Chronology:**

Samtavro (41.8464288, 44.7184942) is located on the right bank of the River Aragvi within the city of Mtskheta. It borders the Aragvi River to the east, Kodmani in the west, Monastrishkhevi in the south, and Baiatkhevi in the north. Samtavro Cemetery is multi-layered. It has existed from the Early Bronze Age until the Middle Ages (from the 2nd half of the 3rd millennium BC to the 11th-12th centuries). The remains of Late Bronze- Early Iron Age, Middle Bronze, Hellenistic, Late Antiquity, and Early Middle Age settlements and more than four thousand different types of burials from the Middle Bronze Age, Late Bronze- Early Iron Age, Antiquity, and early Middle Ages discovered on Samtavro Cemetery.

**B) Excavation history and description of burials:**

Archeological excavations at Samatavro began in the second half of the 19th century. In 1871, tombs were found on the right bank of the Aragvi River in Mtskheta during the expansion of the Georgian military road. For their study, the "Society of Lovers of Caucasian Archeology" sent its member F. Bayern, who conducted excavations from 1871-77. Intensive research on Samtavro Cemetery was carried out from 1938-41 and discontinuously until the present. In 1976, a cist grave was accidentally found, after which the archaeological research of the site resumed. In 1976, a total of 201 tombs were excavated in the Samtavro Cemetery. Among the tombs, there are 89 cist graves,  97 pit burials, nine slab-built burials, and four tile graves. Burial goods were found only in 97 graves. Thirty-eight cist graves, 49 pit graves, nine slab-built burials, and one of the four tile-graves were inventoried. The tombs are dated from the Late Antiquity to the Early Middle Ages: 34 belong to the Late Antiquity period, and 167 to the Early Middle Ages. The burials are dated according to the archaeological inventory.

The site has been studied by archaeologists Andria Afakidze, Aleksandre Kalandadze, and Vakhtang Nikolaishvili, among others. Some of the osteological material is kept at the National Museum of Georgia and another at the anthropology laboratory of the Institute of History and Ethnology. Prof. Malkhaz Abdushelishvili conducted the paleoanthropological study. Currently, most of the material stored at the anthropology laboratory was discovered during the 1975-76 excavations and dates to the Late Antique-Early Middle Ages.

Burial N2 is the slab-built burial of two individuals. One individual was buried in an extended position, and the second individual's skeleton was mixed and didn't show any position. In the tomb were found two gold rings, one gold button, an earring, three silver rings, and beads (35 jet, 14 bugles, and three crystal beads).

Burial N187 is a cist grave that includes seven individuals. Two of them were buried in an extended position, and the other one's skeleton was mixed and didn't show any position. The following burial goods were found: five glassware, one iron clothespin, one silver ring, four gold earrings, beads (34 jet beads, five glasses, two crystals, and sardine beads).

SMT004 is a child whose sex was assigned genetically to a male. The individual had cribra orbitalia and hyperostosis that indicate childhood nutritional stress, trauma, infection, and iron deficiency, likely owing to hereditary anemia.

SMT005 was macroscopically and genetically defined as female. She had a metopic suture.

SMT006 is a female individual of estimated age between 25-30 years. The skull is poorly preserved.

SMT007 is a male individual of estimated age between 30 and 35 years.

SMT008 was macroscopically and genetically defined as female (40-45 years). The skull indicated that she had artificial cranial deformation, circular type.

SMT009 is a female individual with an estimated age between 20 and 25 years. No postcranial skeleton was preserved, but the skull indicated that she had artificial cranial deformation of circular type.

SMT011 is a female individual of estimated age between 25-30 years. The individual has a metopic suture and artificial cranial deformation of oblique type.

SMT012 was macroscopically and genetically defined as female. The age at death is estimated to 60-65 years.

SMT013 was macroscopically and genetically defined as female. The age at death is estimated to 30-35 years. The skull indicated that she had artificial cranial deformation of annular type.

SMT014 is a male adult (30-35 years).

SMT018 is a male individual of the estimated age range 40-45 years. The skull is poorly preserved.

SMT019 is a male individual of the estimated age range 50-55 years. The skull is poorly preserved. The individual had metopic suture.

SMT021- is a male individual of the estimated age range between 30 and 35 years.

SMT022 is a female individual of the estimated age range of 40-45 years. On the right side of the maxilla, the individual had antemortem incisors loss (I, II) and antemortem first molar loss on the right and left side of the Mandible.

SMT023 is a male individual ofestimated age between 50 and 55 years. The individual had periodontal diseases.

SMT024 is a male individual of estimated age between 60 to 65 years. The skull is poorly preserved.

SMT025 was macroscopically and genetically defined as male. The skull indicated that he had artificial cranial deformation- of annular type. On the right mandible, the individual had ante mortem second premolar loss.

SMT030 was macroscopically and genetically defined as female. The age at death is estimated to 50-55 years. The skull is poorly preserved. The mandible indicates that the individual had periodontal diseases.

SMT033 is a female of estimated age between 60 and 65 years. The skull indicated that she had artificial cranial deformation of annular type.

**Tserovani (TSV)**

**A) General location and chronology:**

Tserovani (41.89329, 44.70024) is a village in eastern Georgia in the Mtskheta municipality of the Mtskheta-Mtianeti region. Three archaeological sites were studied in Tserovani: Tserovani I, Tserovani II, and Tserovani III. Tserovani I is located northwest of Tserovni. The inventory of various jar types, curved-inward pots, drinking vessels, and bronze leaf-like daggers date the monument to the 13th-12th centuries BC. Tserovani II is located northeast of the village, on the so-called "Kalandadze Hill." Tserovani II dates back to the 14th-13th centuries BC. Tserovani III is located in Tserovani village and dates back to the 13th-12th centuries BC, based on the curved-inward typology of a jar and a tray.

**B) Excavation history and description of burials:**

**Tserovani I** was discovered and excavated in 1977. As a result of the excavations, 30 heavily damaged pit burials were found, with the dead buried on the right or left side, in a contracted position, head oriented to the north. The inventory in the tombs mainly included jar-type vessels and biconical vessels.

**Tserovani II** was excavated in 1977-79. The studied area measured a total of 2 $km^2$. The findings were agricultural pits and burials, likely dating to the Early Bronze Age and later. The central cluster of tombs is dated to the Late Bronze Age. A total of 75 pit graves with rounded corners were excavated, and some had been plundered. Three tombs belong to the Early Bronze

Age, 64 to the Late Bronze Age, and 8 to the Early Iron Age. Burials were in their majority individual (n=38), and the rest were paired (n=12) and mass (n=4). The dead were placed in a crouched position - either on their right or left side- and their heads oriented north. In nine tombs were buried cattle, and in five tombs were sheep and goats. The tombs included burial goods such as black-burnished vessels and jars, Kakhuri-type daggers, clay horse and sheep statues (toys), bronze ritual items, and jewelry.

**Zhinvali (DZN)**

**A) General Location and Chronology**:

Zhinvali (42.109804, 44.770221) is at the narrowest point of the Aragvi Canyon. A township in Georgia, located in the Dusheti municipality, on the right bank of the Aragvi River, along the Georgian Military Road.

**B) Excavation history and description of burials:**

In 1974, in the village of Zhinvali, a multi-layered cemetery (area XXV) extending on the slope on the right bank of Aragvi was discovered. Twenty-five graves were unearthed, of which seven were empty. The burial ground consisted of usual pit tombs, pit tombs covered with stone slabs, and catacombs. Chronologically, the tombs are divided into two main chronological groups: the first group mainly dates to the 2nd-3rd centuries, and the second to the 4th-6th centuries AD. It is also possible to single out a section of tombs that belongs to the transition period from the Late Antiquity to the Early Middle Ages (3rd-4th centuries AD). The stone tombs and catacombs of the Early Middle Ages were constructed under Christian burial customs and rites.

In Late Antiquity and the transition period, the primary burial type is individual tombs. However, a burial of two individuals buried in a contracted position was also found. The orientation varies, but the graves with the head to the west or southwest prevail. Family tombs in the Early Middle Ages became common, and the dead were predominantly buried in Christian traditions. The inventory of the Late Antiquity group of tombs (N1, 5, 18, 22) includes clay vessels, jewelry, and an iron ring with a Greek inscription of 'Bakur-I' on the intaglio. Interestingly, this name is confirmed in Georgian and foreign written sources and seems widespread between the 2nd and 4th centuries.

From 1974 to 1975, the archaeological expedition of Zhinvali continued to excavate the "Nakalakari " burial ground (district III). The excavations were carried out inside and outside of two churches. The discovered tombs are similar to the previously studied tombs in terms of structure, burial rite, and inventory but date back to the 11th-14th centuries. Ninety-eight tombs from this burial ground have been studied, represented by pit tombs, cist graves, charnel houses, and tombs with built-up walls. The skeleton was found in an extended position in one pit burial 1.7 m long and 0.5 m wide. There are numerous cist graves (e.g., N159, 161, 173, 181, 201, 221, 231). Tombs with built-up walls (e.g., N 171, 178, 187, 218, 220, 228) contained three to ten individuals, and the charnel houses seven to twenty-one individuals. The burial goods included clay urns, one-eared and earless small cups, fragments of glass vessels, bracelets, buttons, beads, ring eye, arrowheads, earrings, rings, pendants, buttons, hooks, and mirrors.

Between 1977 and 1985, about 5.42 $km^2$ of ancient territory was unearthed, revealing about 558 burials dating to c. 1st century BC-8th century AD, numerous settlements of the first half of the first millennium AD, a wine cellar of the 2nd century BC, and some other monuments. The numerous graves were of various types: pit (ground) burials with stone and earth embankments and jar burials, typical of the Late Antique period. In contrast, catacombs and ground burials belong to the Early Medieval period.

The burial customs attested at the Zhinvali cemetery were Christian. However, some exceptions also occur: while in most catacombs, skeletons were buried in a stretched position, in some (n=13), the skeletons were found in a crouched posture. All the catacombs of the cemetery have been discovered at a depth of 1.80 – 3.4 m and were oriented from west to east. First, a vertical pit of circular shape was dug out for their construction, and then the vaulted burial chamber was erected; usually, the pits of the catacombs were filled with coble-stones, and burial chambers were plastered with mortar and surrounded by stone circles. Almost all the catacombs contained collective burials (2-10 individuals), except for four, where single burials were discovered. Grave goods in the catacombs consisted of numerous bracelets, earrings, beads, etc. It should be noted that attachments to clothing (belt buckles, fibulae, buttons, etc.) have not been found, which makes us think that the deceased were wrapped in a piece of cloth, remnants of which were discovered almost in every burial. Burial inventory is similar to the archaeological material found at numerous sites of East Georgia and dates to the 4th-5th centuries AD. In addition, a group of pit tombs which are covered with flat stone slabs dates to the period of transition from paganism to Christianity and have also been identify in the other cemeteries beyond Zhinvali, like Aragvispiri and Akhali Zhinvali.

The material obtained by the archaeological research clearly indicates the strength of the point in the place of Zhinvali. A joint analysis of archaeological materials shows that the settlement expanded throughout Late Antiquity and the Early Medieval periods. Special attention is paid to the ordinary tombs of the Late Antiquity period, which contain a lot of objects related to urban existence: ship seals, glass medallions with Sasanian scenes, and other jewelry. Overall, the Late Antiquity burial ground of Zhinvali has many analogies in terms of structure, burial rite, and the construction of the burial complexes to those of Armaziskhevi, Mtskheta, Ertso Valley, Urbnisi, and other cemeteries, all of which were part of Kartli at its the early stage of feudalization during the first century AD.

The two burial sites, Zhinvali and Aragvispiri, are closely related in terms of the level and manner of artistic decoration of individual objects, stylistic features, and the burial ritual. They also show structural and content similarities with sites including Armazi, Zghuderi, Ertso, Bori, the respective complexes of Ureki, Kldeeti, and others. Upon analyzing the obtained material, it is evident that a group of silver vessels from New Zhinvali and Aragvispiri share a stylistic unity and are closely related to the Mtskheta matches.

**Aragvispiri (AGV)**

**A) General Location and Chronology:**

Village Aragvispiri (42.075237, 44.752158) is located on the second terrace of the Aragvi River, along the Georgian military road, approximately 200 meters from the Dusheti turn. During the construction of the sewage line in the vineyards of the Soviet farm, remains of a settlement and a cemetery were found. Archeological research on Aragvispiri started in 1974. During two archaeological seasons, 46 burials were discovered, of which only six date to the Early Middle Ages. The rest belong to the Late Antiquity period.

**B) Excavation history and description of burials:**

Based on the composition of the burial goods (dishes, bronze or silver rings, and beads), most pit burials belong to ordinary members of society. The group of richly furnished tombs was relatively small. The goods in the richly furnished tombs included silverware, paintings with hunting scenes, rings, parts of belts, gold jewelry, earrings, rings, and gold and silver coins. This composition was similar to the one in richly furnished tombs from Armaziskhevi.

A comparative analysis of the tomb complexes found at Aragvispiri dates the cemetery to the 2nd-4th centuries, and, among them, to single out a large group of tombs of the Late Antiquity. Complexes found in luxurious tombs belong to the second half of the 3rd century and the beginning of the 4th century.

**Nedzikhi (NDZ)**

**A) General Location and Chronology:**

Nedzikhi (42.187288, 44.797053) is located on the right bank of the Aragvi River. Nedzikhi belongs to two chronological periods, the 3rd -4th and 4th -6th centuries AD.

**B) Excavation history and description of burials:**

Between 1985 and 1988, the Zhinvali and Mtianeti archeological expedition of Eastern Georgia, led by Ramin Ramishvili, conducted excavations at the Nedzikhi site. The lower layer of the site contains 223 pit burials of different types, such as burials with timber framing of logs, barrows, and slab-covered burials. In individual burials, the deceased were buried in a contracted position, lying on the right or left side with their head oriented towards the west or south. Grave finds include clay pitchers, drinking vessels, tumblers, pots, glass bowls, beads, bronze belts, fibulae, rings, earrings, iron knives, spearheads, and bronze and animal bone pins. The upper layer of the site consists of stone burials and some pit burials. All burials were Christian, with most individuals buried in a Christian posture, which means they were buried in a lying position. However, one individual was placed in a pre-Christian posture, buried in a contracted position. Artifacts found in the burials include clay pitchers, rings, earrings, bronze and silver Sasanian coins, and others.

NDZ001: Burial number is unknown. This individual was macroscopically and genetically defined as female. The age at death is estimated to 50-55 years. The skull was damaged. There are no apparent pathologies.

NDZ002: The burial number is unknown. This individual was macroscopically and genetically defined as female. The age at death is estimated to be 50-55. The skull indicated that he had artificial cranial deformation of the oblique type. The individual has wormian bone (Lambdoid suture) and metopic suture .

NDZ003: Burial number is unknown. This individual was macroscopically and genetically defined as male. The age at death is estimated to 60-65 years. The skull is damaged. There are no apparent pathologies.

NDZ004: The burial number is unknown. This individual was macroscopically and genetically defined as female. The age at death is estimated to be 45-50 years. The skull indicated that he had artificial cranial deformation- of tabular oblique type. The individual has a metopic suture, wormian bone, which is located on the left side of the lambdoid suture. The maxilla indicates that she had AMTL on the left maxilla (3,4).

NDZ005: The burial number is unknown. This individual was macroscopically and genetically defined as female. The age at death is estimated to 50-55 years. The skull indicated that he had artificial cranial deformation- of tabular oblique type. She had metopic suture.

NDZ006: The burial number is unknown. This individual was macroscopically and genetically defined as male. The age at death is estimated to be 45-50 years. The skull is poorly preserved. There are no apparent pathologies.

NDZ007: The burial number is N72. This individual was macroscopically and genetically defined as female. The age at death is estimated to 65-70 years. The skull indicated that he had artificial cranial deformation- of tabular type. On the left side of the maxilla, she had caries (8).

NDZ009: The burial number is N41. This individual was macroscopically and genetically defined as male. The age at death is estimated to 50-55 years. The skull indicated that he had artificial cranial deformation- of tabular oblique type. On the left side of the maxilla, he had AMTL (1,3,4). The individual had metopic suture.

NDZ010: The burial number is unknown. This individual was macroscopically and genetically defined as male. The age at death is estimated to 50-55 years. The skull indicated that he had artificial cranial deformation- of oblique type. The individual had metopic suture and wormian bone (Lambdoid suture).

Out of the 9 skulls studied (**Table 1**), 3 skulls were found to have WB and 5 skulls had metopic sutures, which were located in the sagittal sutures (4 cases) and squamosal sutures (1 case). 4 skulls were found to have AMTL and 6 skulls have artificial cranial deformation. All deformed skulls had metopic sutures, besides 1 skull (NDZ007).

**Table 1. Osteological context of Nedzikhi individuals**

| Individual N | Wormian Bone | Metopic suture | Caries | AMTL | ACD/Non-deformed | Sex | |
|---|---|---|---|---|---|---|---|
| NDZ001 | - | - | - | - | Non-deformed | F | |
| NDZ002 | + | + | - | + | ACD | | M |
| NDZ003 | - | - | - | - | Non-deformed | | M |
| NDZ004 | + | + | - | + | ACD | F | |
| NDZ005 | - | + | - | - | ACD | F | |
| NDZ006 | - | - | - | - | Non-deformed | M | |
| NDZ007 | - | - | + | - | ACD | F | |

| | | | | | | |
|---|---|---|---|---|---|---|
| NDZ009 | - | + | - | + | ACD | M |
| NDZ010 | + | + | - | + | ACD | M |

**Tchiatura (TCH)**

**A) General Location and Chronology:**

Tchiatura (42.286799, 43.276112) is located in the Municipality of Tsinsopeli Castle, about 20 km west of Sairkhe village. It is situated on the left bank of the river Kvirila. Previous archaeological works have identified two settlements in the area - one near Tsinsopeli Castle and another at Jieti. Both of these settlements are surrounded by a wide wall made of dry stone, with gravel filling the area between them. The Early Antique buildings have been identified on the southern slope of Tsinsopeli Castle, including a metal soldering workshop. The monuments of the Early Antique period at Jieti were destroyed due to earthworks, leaving only material from the first half of the 1st millennium BC well preserved. In addition to these monuments, cemeteries of the 2nd -4th centuries AD were discovered at Jieti.

**B) Excavation history and description of burials:**

The State Museum of Art of Georgia, under J. Nadiradze, conducted an archeological expedition in the Kvirila Valley of Tchiatura between 1976 and 1980. The excavated archaeological materials were quite diverse and spanned a long period of time. The team discovered a settlement at Jieti with six phases of occupation, with its chronology ranging from the Early Bronze Age to Early Antiquity. N. Tushabramishvili considered the Stone Age samples found at Jieti to be from a Paleolithic open-air site, with dates assigned to the Mousterian period. The cemetery at Jieti dates to the 2nd-4th centuries AD. The remains of the medieval level have also been revealed at the southeastern slope of the Tsinsopeli castle. Not far from the fortress of Tsinsopeli, the team also excavated a multi-layered settlement dating back from Early Bronze Age to Early Antique periods, a Paleolithic open-pit stone quarry, and a cemetery from the first half of the 1st millennium BC.

TCH001 was macroscopically and genetically defined as female. The age at death is estimated to 50-55 years. The skull indicated that he had artificial cranial deformation- of oblique type. The individual has hyperostosis.

TCH002 was macroscopically and genetically defined as male. The age at death is estimated to 20-25 years. There are no apparent pathologies.

TCH003 was macroscopically and genetically defined as male. The age at death is estimated to 40-45 years. On the left side of the maxilla, the individual had ante mortem tooth loss (1), and on the right maxilla, he had caries (7).

TCH004 was macroscopically and genetically defined as male. The age at death is estimated to 20-25 years. The skull indicated that he had artificial cranial deformation- of oblique type. The individual has metopic suture.  The individual had caries on the maxilla's right side (4).

TCH006- was macroscopically and genetically defined as male. The age at death is estimated to 25-30 years. There are no apparent pathologies.

TCH007 was macroscopically and genetically defined as male. The age at death is estimated to 50-55 years.  The skull indicated that he had artificial cranial deformation- of oblique type.

TCH008 was macroscopically and genetically defined as male. The age at death is estimated to 45-50 years. On the left side of the mandible, the individual had ante mortem tooth loss (5,6), and on the right side of the maxilla, he had caries (7).

TCH009 was macroscopically and genetically defined as male. The age at death is estimated to 25-30 years. The skull indicated that he had artificial cranial deformation- of oblique type. The individual have metopic suture and  wormian bone (lambdoid suture).

TCH010 was macroscopically and genetically defined as male. The age at death is estimated to 50-55 years. . On the left side of the mandible, the individual had caries (6).

Out of the 9 skulls studied, 5 skulls have artificial cranial deformation. 2 skulls were found to have AMTL, 3 skulls-metopic suture and 4 skulls have caries (**Table 2**).

**Table 2. Osteological context of Tchiatura individuals**

| Individual N | Wormian Bone | Metopic suture | Caries | AMTL | ACD/Non-deformed | Sex | |
|---|---|---|---|---|---|---|---|
| TCH001 | - | - | - | - | ACD | F | |
| TCH002 | - | - | - | - | Non-deformed | | M |
| TCH003 | - | + | + | + | Non-deformed | | M |
| TCH004 | - | + | + | - | ACD | | M |

| | | | | | | | |
|---|---|---|---|---|---|---|---|
| TCH006 | - | - | - | - | Non-deformed | | M |
| TCH007 | - | - | - | - | ACD | | M |
| TCH008 | - | - | + | + | Non-deformed | | M |
| TCH009 | - | + | - | _ | ACD | | M |
| TCH010 | - | - | + | - | Non-deformed | | M |

**Kamarakhevi (KMR)**

**A) General Location and Chronology:**

The Kamarakhevi cemetery (41.8569386, 44.7060885) is located to the south of Tsitsamuri village (Mtskheta district), in the plain on the left bank of the river Aragvi and its tributary Great Kamarakhevi. Archaeological excavations were carried out in 1953, and 1976-77 led by A. Afakidze, R. Ramishvili, and T. Zhgarkava. The tombs discovered on Kamarakhei are divided into three chronological groups: Group I - tombs of the 6th-5th centuries, Group II - tombs of the 5th-4th centuries, and Group III - tombs of the 4th-3th centuries BC.

**B) Excavation history and description of burials:**

In the 1953 excavation directed by T. Jgarkava, 39 tombs of the Kamarakhevi cemetery were foundOut of these, 34 were excavated, and 29 were inventoried. In some tombs, jewelry and pottery were found in large quantities. It should be noted here that most children were buried without burial goods.  The dominant burial type was pit burials, followed by cist graves (4/39; burials N6,11,12,32). In total, more than 100 tombs have been excavated at Kamarakhevi. Burial customs are almost the same in all cemeteries. The deceased was buried with their limbs folded on the right or left side with the skull placed on a cobblestone gravestone. Some of the burials were surrounded by pottery sherds and cattle bones.

An essential type of burial good of the Karamakhevi complex was clay vessels. In most cases, two or three vessels were placed in the tombs, rarely one, and in one case, four pieces. In a few graves, there are no clay vessels at all. Pottery was mainly used for domestic purposes, and jewelry was found in most of the burials. In the general characterisation of Kamarakhevi materials, attention is drawn to the fact that the weapons were made of iron. Among the jewelry, beads made of semi-precious stones and glass, mainly characteristic of "luxurious" complexes, are worth noting.

There are many similarities in the burial practices of Kamarakhevi with other sitesin the Caucasus, especially in Eastern Transcaucasia. Some of the bronze bracelets found in Kamarakhevi are similar to those found in Beshtasheni, Manglisi, Didube, and Tskneti. Equally important for the dating of Kamarakhevi is the rich archaeological material found in the Kazbegi cemetery, among which, in addition to bracelets, there are rings, beads, and others, which hold many similarities with the Akhalgori hoard.

The burial complex of Kamarakhevi can be divided into three chronological groups.The first group includes burial complexes that contain ceramic products, bracelets decorated with snake heads, massive head buckles, seals with images typical of the Achaemenid period, bent iron knives and various beads including sardine beads. This group of the material dates back to the 6th-5th centuries BC. The second group of Kamarakhevi burials is the largest. It includes the complete complex of Kamarakhevi pottery, finer-headed hinges, arc-shaped bracelets made of both bronze and iron, iron-tongued bow straps, curved iron knives, and unthreaded iron spearheads. The material in this group mainly dates to the 5th-4th centuries BC. The third and the latest phase (4th-3rd centuries BC) includes cist graves and an ancient burial rite, which means the deceased were buried on the right or left side with their limbs folded. Although there are fewer burials in this group, it is clear that bronze jewelry is greatly reduced, there are no sardine beads, and military and agricultural tools are made of iron. Ceramic products in this group are relatively late handiwork.

Tomb N23 - This tomb was located in the southern part of the trench, with stonework that sloped from the northeast to the southwest. It was made up of large boulders with cobblestones in between. Two individuals, an adult and a child, were found inside. In addition, the entire skeleton of a domestic animal was found under the same stone. The two individuals were buried simultaneously, and a common stone was placed in the tomb. The only difference was that the child was placed 10-15 centimeters higher than the adult. The adult’s skeleton was inclined from northwest to southeast, crouched on the right side. Meanwhile, the child was buried to the west of the adult, lying on the left side in a crouched position with the head oriented north. A clay jar, a pitcher, and a tumbler were found on the adult skeleton, while a jar, a pitcher, and a bronze statuette were discovered near the child’s skeleton.

KMR001 was macroscopically and genetically defined as female,40-45 years. Skull is poor preserved. There are no apparent pathologies.

KMR002 is a child. There are no apparent pathologies.

KMR003 was macroscopically and genetically defined as female, 66-70 years. On the left and right side of maxilla, he had AMTL (L4,5,6,7,8; R-5,6,7).

KMR004 was macroscopically and genetically defined as male, 45-50 years.

KMR005 was macroscopically and genetically defined as male, 50-55 years. The individual has wormian bones. On the left and right side of maxilla, he had AMTL (L 6; R-6,7,8).

KMR006 was macroscopically and genetically defined as female, 30-35 years. There are no apparent pathologies. Y-pestis signal

KMR007 was macroscopically and genetically defined as female, 45-50 years. The skull is poor preserved. There are no apparent pathologies.

KMR008 was macroscopically and genetically defined as female. The skull is poor condition. On the Left and right side of mandible, she had AMTL ( L 1,2, R 1,5,6).

KMR009 was macroscopically and genetically defined as female, 45-50 years. On the right side of maxilla this individual has AMTL (8).

KMR010 was anthropologically defined as female, 45-50 years. The skull is poor preserved. There are no apparent pathologies.

KMR011 was macroscopically and genetically defined as male, 30-35 years. The skull is poor preserved. There are no apparent pathologies.

KMR012 was macroscopically and genetically defined as female, 55-60 years. There are no apparent pathologies.

KMR013 was macroscopically and genetically defined as male, 30-35 years. There are no apparent pathologies.

KMR014 was macroscopically and genetically defined as male, 60-65 years. On the right side of maxilla this individual has AMTL (6,7,8).

KMR015-was macroscopically and genetically defined as male, 60-65 years. On the maxilla this individual has AMTL. There are no other pathologies.

KMR016 was macroscopically and genetically defined as female, 45-50 years. The skull is poor preserved. There are no apparent pathologies.

KMR017 was anthropologically defined as child. There are no apparent pathologies.

KMR018 was macroscopically and genetically defined as male, child. There are no apparent pathologies.

KMR019 was anthropologically defined as male. This individual has wormian bone. There are no apparent pathologies.

KMR020 was anthropologically defined as female, 50-55 years. The skull is poor preserved. There are no apparent pathologies.

KMR021 was macroscopically and genetically defined as male. The skull is poor preserved. On the left and right side of maxilla, the individual has AMTL (L-1,2; R-1,5,6).

KMR022 was macroscopically and genetically defined as female, 45-50 years. The skull is poor preserved. There are no apparent pathologies.

KMR023 was macroscopically and genetically defined as male, 35-40 years.  There are no apparent pathologies.

KMR024 was macroscopically and genetically defined as female, 45-50 years. Looks like cancer. The individual has wormian bone. Left side of mandible, the individual has carries (6).

KMR025 was macroscopically and genetically defined as female. There are no apparent pathologies.

KMR026 was macroscopically and genetically defined as male, 65-70 years. On the right and left side of maxilla, the individual has AMTL (R- 7,8; L- 5,6), also the individual has carries on $8^{th}$ teeth (right maxilla).

KMR027 was macroscopically and genetically defined as male, 45-50 years. There are no apparent pathologies.

**Nabagrebi (NBG)**

**A) General Location and Chronology:**

The cemetery (41.80549 , 44.57317) is located in the Nabakara village, to the east of Satovle-Nabagrebi, on the left bank of the Khekordze tributary called Patara Khevi. On the right bank of Patara Khevi, there is a hill settlement called Abuleti Gora, which is contemporary with the Nabagrebi cemetery. It stands at the confluence of Patara Khevi and Abuleti Khevi' on the cape. Pottery fragments from the Late Bronze to the Early Iron Age are exposed on the southern slope of the settlement mound. The settlement mound was traced in 1954 by the historical geography survey expedition of the Ivane Javakhishvili Institute of History and Ethnology, led by Acad. N. Berdzenishvili. Based on archaeological surveys and excavated materials, it is evident that the area of Satovle village was inhabited from the Bronze Age to the Middle Ages.

**B) Excavation history and description of burials:**

Six trial trenches were excavated in the possible location of the cemetery. Seven stone pit burials of the Early Iron Age were found in the first trench, while the fourth trench yielded two cist graves from the Middle Ages (6th-9th centuries AD). Four dead were buried in one cist grave and one in the other. Trenches revealed a replacement cultural layer consisting of red and straw-colored pottery fragments that belonged to the settlement of the early Middle Ages.

Only stone pit tombs from the Early Iron age were identified, with medium-sized crushed stones and cobblestones unsystematically laid on top of the graves. The dead are buried in a crouched position on the left or right side, with their heads facing west. Ceramics were the main archaeological material found in the tombs, followed byiron weapons and jewelry.

Based on material typology, the stone pit tombs of the Early Iron age excavated at Nabagrebi Cemetery should be relatively dated to the 8th-7th centuries BC. However, according to the bronze and iron hinges with tubular and fused glass heads, the cist graves could date earlier to the 6th-8th centuries AD.

Of the seven burials that were dated to the Iron Age, only six were identified as individual burials. One burial was unable to be sorted due to the lack of osteological material. During the Iron Age, the population demographic of the area was predominantly male with 61.95%, followed by females with 36.87%, and children with 1.18%. The average age of males during this time period was 52 years, while the average age of females was 31 years. In the Early Middle Ages, the average age of males was 56 years.

**Treli (TRL)**

**A) General Location and Chronology:**

Treli (41.766305, 44.7670064) is located in the Didube-Chugureti district of Tbilisi. It’s situated on the right bank of Mtkvari in the Digomi Valley.

The layers of the earliest occupation in the Digomi arrea and its adjacent territory belong to the Kura-Araxes culture. The settlement from this period is located on both banks of the Dighmura River, south of the Digomi Valley, at the foot of the eastern slopes of the hills of the Georgian Military Road. The remains of this settlement, spread over several hectares, suggest that it was quite large at the time.

**B) Excavation history and description of burials :**

The Treli settlement from the Kura-Araxes culture is multi-layered. At least three stratigraphic layers have been identified in some areas of the excavated region. Two burials of the same period (N26 and N54) were found within the Treli settlement.

Burial N26 was likely represented a kurgan, where a gold-painted skeleton was found. This type of ritual was widespread in the 4th-3rd millennia BC, from Western Siberia to Southeastern Europe.

Burial N54 was a pit burial mound. Therefore, the pottery found in the tomb, with its specific features, is believed to belong to the late stage of the Early Bronze Age.

Between 1968 and 1975, 129 tombs were excavated at the Treli Cemetery. Of these,two burials were from the Early Bronze Age, eight from the Middle Bronze Age, twelve from Antique Period, two from the Middle Ages, and the remaining burials belonged to the Late Bronze Age. All of the Middle Bronze Age burials in Treli Cemetery were individual burials. In most of the graves, inhumation of the deceased was confirmed, and in graves N53 and N104, cremation was probable. In those tombs where inhumation was confirmed, both females and males  were buried on the right side in a crouched position.

It is known that in the Late Bronze Age, the Samtavro Cemetery in Eastern Georgia had a custom of buring males on the right side and females on the left, individually. This same practice was also presented in Treli and other sites in the region synchronous to Samtavro. This custom of indiviudal burials was only established during the Late Bronze Age. The fact that instead of the collective burials common in Middle Bronze Age, only individual burials have been found in the cemetery should remind us of the advancement of the population of Treli compared to the contemporary tribes living in the mountain front of Eastern Georgia.

The type of tombs and associated materials discovered in the Treli Cemetery started to chang since the 13$^{th}$ century BC. However, starting from that time, the burial practice of Treli and Samtavro show the closest similarity to each other.

**Didnauri (DDN)**

**A) General Location and Chronology:**

Didnauri (41.46528, 46.10556) is located in the western part of the large Shirak Valley, on the 9th kilometer of Dedoplistskaro-Lower Ridge road. It was discovered thanks to satellite photo decoding. According to the ceramic material found during the excavation of the fence that belonged to its builders, the former city dates back to 12th-11th centuries BC. However, the city ceased to exist in the first centuries of the 1st millennium BC, as indicated by the ceramic material found in the defensive trench.

**B) Excavation history and description of burials:**

During the months of April to June 2015, the Kakheti archaeological expedition, which was financed by the National Agency for Cultural Heritage Preservation, carried out field works in Didnauri. The excavation was led by Davit Naskidashvili, with Konstantine (Kiazo) Pitskhelauri as a consultant. The excavation focused on the southern fence of the gate of the former city and the surronding cemetery. The expedition carried out works on the cemetery in three areas divided by vertical sections.

During the excavation of the cemetery, it was observed that water lines were present in the main trench, extending from west to east, with a large number of river stones and crushed stones. This suggests that the area was covered by cobblestone due to water flowing downstream. That is why the southern part of the cemetery should be significant and extensively used. The former city's main building for water supply was located approximately 300m away in the northwest of the cemetery, on a slight rise. Nearby, at the southern shore of the river Didnauri, a large water reservoir seems to have existed and was abutted by a large dam from the eastern side. Cobblestones were used for making the dam. The former Didnauri riverbed was located about 180m north of the cemetery. The alluvial layers found in the cemetery were not the result of river flooding. The provenance of the stones in the alluvial layers was from outside of Shiraki Valley, but were brought from the banks of Alasani River around 20 km to the northeast of Didnauri and was used to reinforce the dam of the main water supply building in the former city. As a result of dam disorder and flooding, the cemetery area was covered by crushed stones. Nevertheless, the expedition was able to excavate four tombs dating to Late Bronze Age. Also, it should be noted that many obsidian flakes were found in the excavation, indicating the existence of burials dating to the early stages of the Late Bronze Age and earlier.

**Fiqrisgora (FQR)**

**A) General Location and Chronology:**

Fiqrisgora Cemetery (41.85, 44.72) is situated in the valley of the Samtavro, which lies between Baiatkhevi, Monastery Gorge, Kodmani, and the river Aragvi River. This area includes Samtavro Monastery and cemetery, Baiatkhevi Monastery, Mtskheta Central Square, and Mtskheta Cloister. Thus, Fiqrisgora is in the buffer area of the world heritage monuments of Mtskheta and is part of its exceptional public value. Over time, 179 tombs have been excavated in Fiqrisgora, including 152 stone tombs, 17 pit tombs, one charnel house with stone blocks, one combined tomb, two clay plinth graves, and the remains of two tombs. Of these, 75 tombs are individual adults, 16 are of children, and the remaining 88 are collective. Based on the analysis of material and tomb architecture, the cemetery was in use during the 4th to 6th centuries.

**B) Excavation history and description of burials:**

Fiqrisgora is currently inhabited by the population of Mtskheta. In the 1970s, Mtskheta district hospital was constructed on the site, leading to the discovery of 12 graves that were excavated and studied in 1976 by the permanent expedition of Mtskheta, led by A.Afakidze and T.Jgharkava. More recently, in 2014, when a private owner started building a house in the southeastern part of Fiqrisgora, several burials were discovered, leading to the identification and study of 64 tombs by an expedition led by N. Maisurashvili and joined by other professionals. Unfortunately, soil erosion processes caused by heavy rains in January and February 2016 damaged eight cist graves, with one of them collapsing. To address this issue, the Great Mtskheta Archaeological State Museum-Reserve organized an expedition to work on the cemetery. As of 2022, 99 tombs were identified and studied.

**Saphar-Kharaba (SAP)**

**A) General Location and Chronology:**

Saphar-Kharaba (41.642448, 44.123947) burial ground with an area of 1500m x 500m, is situated in Trialeti, north of the village Saphar-Kharaba in the Tsalka district. This cemetery dates back to 15th-14th centuries BC.

**B) Excavation history and description of burials:**

Between 2003 and 2005, a total of 115 pit graves were excavated, which were surrounded by cromlechs made with large basalt stones. As a rule, one grave oriented from north to east was fixed in the center of the cromlech. However, in the case of graves 67 and 68, they were found inside a single cromlech. The deceased was found lying on their right or left side, oriented to the north. Four of the 115 unearthed graves (No. 10, 23, 30, 90) were distinguished by their size. In these graves, the deceased was

placed on wooden burial beds, or buried with chariots or some of their components. The burial goods are mainly represented by clay wares, with one group of pottery being thin, black burnished and decorated with polish ornament. The second group wass made from coarse-grained clay, brownish, and decorated by relief bands. Along with clay wares, the graves also contained daggers of Near Eastern style, scabbards, arrowheads, lancet-like weapons with bone handles, pyramidal stones, authoritative insignias, and Mitanian cylinder seals of the "Common Style".

**Kiketi(KKT)**

**A) General Location and Chronology:**

The Kiketi Cemetery (41.655494, 44.654936) is located in the Gardabani district, near Kiketi village, on the road from Kojori to Manglisi. It is situated in the historical Kvemo Kartli region. The site contains ceramic complexes that are typical of the Early Bronze Age culture. These belong to the initial stage of the Kura-Araxes culture and also reveal the characteristics of the initial elements of the Early Bronze Age of Kvemo Kartli.

**B) Excavation history and description of burials:**

Kiketi was discovered in 1920 and was first tested by E. Pchelina. In 1948 excavations were conducted by the State Museum of Georgia under the leadership of B. Kuftin. In 1961, a team from the Tetritskaro Archaeological Expedition of Javakhishvili Institute of History, Archeology, Ethnography, led by G. Pkhakadze, conducted further excavations at the same cemetery. The cemetery is located on a slope and extends over an area of 120 $m^2$.Excavations at Kiketi cemetery uncovered 21 tombs, including charnel houses and barrow pit graves. Underneath individual tombs or beside them, there were empty piles of stones and round pits filled with charcoal, ash, ceramic slag, fragments of stone grinders, and other products that predated the tombs.The burial rule had a certain regularity to it, where the charnel houses were oriented from north to south and the skeletons buried in a crouched position. In barrow pit graves, skeletons were sometimes found without skulls. The burial goods consisted mainly of clay vessels.

**Natakhtari (NAT)**

**A) General Location and Chronology:**

Natakhtari (41.919444, 44.727222) is a historical monument village located on the right side of the Aragvi River, 14 km away from Mtskheta. The village was previously known as Safurzle, named after the variety of silk mulberry trees that was used to grow there. It was mentioned in historical sources from the 2nd century, when King Amazasp of Kartli defeated the Alan army there.

During archaeological excavations led by A. Apakidze, the remains of the Middle and Late Bronze Age, Early Iron Age, and the middle ages settlements were discovered.

**B) Excavation history and description of burials:**

Layer I contained the remains of a stone-paved floor, as well as the bases of a pinkish-burnt thin-walled pitcher and fragments of clay vessels that were characteristic of the early Middle Ages.

Layer II contained burials from the Late Antiquity period. The dead were buried extended, with their heads oriented to the west and found with clay vessels, jewelry, and silver coins of Octavian Augustus.

Layer III revealed a Middle Bronze Age hill burial and catacomb-type burial structures, including two cenotaphs. Inside the tombs,black glossy relief-ornamented clay vessels, red englobed, chevron-decorated drugs, bronze hinges, and bronze stalked daggers were found.

Layer IV revealed Late Bronze to arly Iron Age barrow pit burials. The dead were buried in a crouched position. Black and gray clay pots, bronze and iron weapons, and jewelry were found in the tombs.

**Khimshiaantmitsebi (KIM)**

**A) General Location and Chronology:**

The site (41.7774711, 45.7671037)  is located in the southwest of the homonymous village, on the left bank of the Aragvi River. The cemetery found at this site is multi-layered and covers a wide range of time periods, from the 5th millennium BC to the 6th century AD.

**B) Excavation history and description of burials:**

In 1984-86, excavations were carried out by the archaeological expedition of Zhinvali under Ramin Ramishvili. The lower layer dates back to the 5th millennium BC and includes fragments of ceramics, flint, and obsidian, and other materials.

The next layer dates from the end of the 4th millennium BC and the beginning of the 2nd millennium BC. During excavation, five pit burials, twenty-six ritual pits, and six small hearth-altars, similar to the earlier so-called cylindrical hearths  were discovered. The pits and hearths contained fragments of pottery from the Kura-Araxi culture, bone and stone products. The skeletons were mostly flexed on the left or right side. Burial goods consist of a clay net, a bone crucible, and barrel-shaped paste beads.

Nine tombs of the Middle Bronze Age were excavated at a depth of 1.4-2.15 meters from the ground surface in the cemetery. These tombs date back to the second quarter of the 2nd millennium BC. Out of the nine tombs, four are of the catacomb type, and five are cist graves buried under barrows. Only one of the catacombs contained a deceased person, whose skeleton was found in a flexed position on the left side with the head oriented to the southeast. The other three burials were cenotaphs, in which one or two sheep skeletons were found with obsidian growths around their necks. The most abundant burial goods that were recovered from these catacombs were ceramics. Reddish brown ware decorated with combed geometric ornament and warty cups were found. The jewelry discovered consisted of bronze studs with sardine heads, sardines, and beads of various shapes and sizes. The skeletons found in the cist graves were tightly flexed, lying on the right or left side, with the head oriented towards the northwest and southeast. In these tombs, black-glossy clay vessels with pink handles, which were characteristic of the Trialeti culture, were found. Other items discovered included a bronze lancet with a stem, buckles, sardines, paste, and plaster beads.

Nineteen tombs date back to the 15th - 13th centuries BC. The tombs were made of small, semi-circular stones. The skeletons were on their right side, in a tightly flexed position. There was also a type of secondary burial where the skeletons were placed on any side. One of the tombs was found with  the deceased buried with a horse. Burial goods of this tomb mainly consisted of ceramics.

Three pit burials and several pits, dated back to the 3rd - 4th centuries AD, were found to contain a gold ring, a gold earring, a silver earring, a rosette, beads, a bronze ring, sardine and glass beads, among other objects.

Nine tombs from the 4th to 6th centuries were excavated in the cemetery. The tombs are oriented from east to west. Seven of these tombs contain the remains of individuals buried under Christian rule. In the other two tombs, the deceased were found lying on their sides in a flexed position, with their heads oriented to the west. Out of the nine burials, only four contained gold and silver earrings as burial goods, while the remaining five were without any burial goods.

#### Samshvilde (SVL)

#### A) General Location and Chronology:

Samshvilde (41.507222, 44.505556), a historical city of the Kvemo Kartli region in the southern part of Georgia, is a complex and multi-period archaeological site. The city occupies a strategic and defendable location on a long basalt plateau above ravines formed by the Khrami and Chivchava rivers. This distinctive landscape position and environmental conditions that include a mild climate and abundant natural resources, have attracted human occupation for millennia. While Samshvilde and its surroundings may have been inhabited since the Neolithic era, the urban complex mainly dates back to the medieval period, when it became the region’s principal fortress and political-economic centre. Its proximity to the northern branch of the Silk Road further increased its importance, making it a melting pot of various ethnic groups and cultures, which is reflected in the archaeological remains. Despite being an outstanding archaeological complex, the site has never been the subject of a full-scale archaeological investigation. Only small-scale fieldwork was carried out during the Soviet and post-Soviet periods, which did not provide details on the site's stratigraphy and chronology or the distribution of cultural features and monuments. In contrast, there has been a fairly extensive archaeological investigation of the surrounding Kvemo Kartli region. To address this gap, the Samshvilde Archaeological Expedition of the University of Georgia conducted five seasons of fieldwork from 2012 to 2016.

**B) Excavation history and description of burials:**

SVL005 was macroscopically and genetically defined as female. The age at death is estimated to 50-60 years. The skull indicated that he had artificial cranial deformation- of circular type. On the left side of the maxilla, the individual had ante mortem tooth loss, also on the left mandible, She had ante mortem tooth loss (5,6) and also on  right side (1,7). The individual has metopic suture, wormian bone (Lambdoid suture) and hyperostosis.

SVL006 was macroscopically and genetically defined as Male. The age at death is estimated to 7-9 years. On the right side of the maxilla, the individual had ante-mortem tooth loss (7,8).

SVL007 was macroscopically and genetically defined as male. The age at death is estimated to 40-45 years. The individual had metopic suture and wormian bone (lambdoid suture).

SVL008 was macroscopically and genetically defined as Female. The age at death is estimated to 10-12 years.

SVL011 was macroscopically and genetically defined as Female. The age at death is estimated to 30-35 years. The skull indicated that he had artificial cranial deformation- of Oblique type.

SVL012 was macroscopically and genetically defined as male. The age at death is estimated to 50-60 years. The skull indicates that he had wormian bones(os Inca bone and lambdoid suture)- abnormal ossicles that develop from extra ossification centers within the cranium. The individual also had metopic suture. On the left side of the maxilla, the individual had ante mortem tooth loss (6,7).

SVL013 was macroscopically and genetically defined as Female. The age at death is estimated to be 50-60 years. The individual had metopic suture and wormian bone.  On the left side of the maxilla, the individual had ante mortem tooth loss (1, 3,6,7), also on the right side of the maxilla (4).  The individual has osteoma, which means that reason for death was cancer.

SVL015 was macroscopically and genetically defined as male. The age at death is estimated to 20-30 years. The skull indicated that he had artificial cranial deformation- of fronto-occipital type.

SVL016 was macroscopically and genetically defined as Female. The age at death is estimated to 35-40 years. The skull indicated that he had artificial cranial deformation- of Oblique type. The skull had wormian bone (lambdoid suture). On the left side of the maxilla, the individual had ante mortem tooth loss (6,8), also on the right side of the maxilla (3,5,6,7,8).

Out of the 10 skulls studied, 5 skulls were found to have WB and metopic sutures, which were located in the sagittal sutures (4 cases) and squamosal sutures (1 case). 6 skulls were found to have AMTL and 4 skulls have artificial cranial deformation. All deformed skulls had wormian bones and metopic sutures, besides 2 skulls (SVL015, SVL011)(**Table 3**). Wormian bones are usually small irregular ossicles located within the cranial sutures. They are formed as a result of alterations in the normal formation of the flat bones of the skull and are usually regarded as normal variants. Pathological, mechanical, and genetic factors have been proposed as the primary causal mechanism in the occurrence of Wormian bones. The reported incidence is variable, ranging from around 10% (in Caucasian skulls), through 40% (in Indian skulls), to 80% (in Chinese skulls). In general, males are more frequently affected than females. According to some authors, the occurrence of WB is controlled by genetic factors. Some other studies suggest that the presence of WB is associated with cranial and central nervous system abnormalities (A.A. Khan et al. 2011). Metopic suture is a non-metric trait to have a genetic component that can be influenced or modified by epigenetic variables resulting from the external environment and internal physiology.

Out of the ten skulls studied, just one skull was found to have osteoma. An osteoma is an osteogenic tumor composed of well-organized mature bone. Osteomas of the cranial vault and mandible consist of dense mature lamellar bone and are less common clinically.

Permanent teeth may be lost prematurely through trauma, chronic pathology, or intentional ablation. This may be a sequel to gross caries, root caries, pulp chamber exposure and periapical infection, severe attrition with continuous eruption, or periodontal disease. An examination of the pattern on AMTL in individuals is required to investigate possible ablation as intentional removal will often be symmetrical or patterned. Most studies that observe AMTL in archaeological samples are typically associated with massive caries and/or severe macrowear with the subsequent exfoliation of the tooth.

Periodontal disease is initiated by polymicrobial plaque biofilm or abrasive effects of calculus, which are associated with the build-up of plaque. The build-up of plaque then causes an inflammatory response in the periodontal tissues of the teeth.

**Table 3. Osteological context of Samshvilde individuals**

| Individual N | Wormian Bone | Metopic suture | AMTL | ACD/Non-deformed | Sex | |
|---|---|---|---|---|---|---|
| 2869 | + | + | + | ACD | F | |
| 2867 | - | - | - | ACD | | M |
| 2879 | + | + | + | Non-deformed | | M |
| 2881 | + | + | - | Non-deformed | | M |
| 2882 | - | - | - | ACD | F | |
| 2883 | - | - | - | Non-deformed | F | |
| 2884 | + | + | + | ACD | F | |
| 2871 | + | + | + | Non-deformed | F | |
| 2874 | - | - | + | Non-deformed | F | |
| 2876 | - | - | + | Non-deformed | | M |

**Vani(VNI)**

**A) General Location and Chronology:**

Vani (42.0878157, 42.5123899) is an ancient city that existed between 7th -1st centuries BC. Materials from the Late Neolithic and Middle Ages have also been discovered in Vani. It is located on the western bank of the Sulori River at its confluence with the Rioni River, in the Colchis Lowlands. The city is situated on a hill bordered by two ravines and overlooks the plains through which the Rioni River flows and offers beguiling views of the Sulori River Valley with its surrounding hills and the Meskhetian Mountain Rangein the background. The strategic location made it the political, economic, and spiritual center for the ancient Colchians, with four distinct stages of uninterrupted occupation identified. Despite this, the name of this ancient city is still unknown.

**B) Excavation history and description of burials:**

The discoveries of ancient artifacts in Vani was first reported in the local press in 1876. Ekvtime Takaishvili conducted the first archaeological survey in Vani in 1896 and wrote the first archaeological publication about Vani.

In the 1930s, academician Niko Berdzenishvili, along with L. Muskhelishvili and N. Khoshtaria, conducted archaeological explorations in Vani and its surroundings.

The systematic archeological study of Vani settlement was started in 1947 under Nino Khoshtaria and continued until 1963 with minor gaps.

The archeological site of Vani comprises multiple cultural layers dating back from the Iron Age to the Hellenistic era. The settlement in Vani existed continuously for eight centuries, from the 8th to the 1st century BC. The earliest stage of ancient Vani dates back to the 8th - 7th centuries BC. However, the material of this period is relatively scarce, which may have been damaged by later construction activities. During this time, the city was supposed to be a cult center of the nearby areas, and the well-preserved cult complex on the city's central terrace is a testament to this. The second stage, which dates back to the 6th - 4th centuries BC, includes wooden buildings, a ritual square, a rock-cut complex, luxurious tombs, and other monuments. This was the time when ancient Vani was one of the political-administrative centers of the Kingdom of Colchis. Archaeological excavations have uncovered specimens of Colchian goldsmithery of this period. In the third stage, which lasted from the second half of the 4th century BC to the beginning of the 3rd century BC, the rulers of the Vani region gained some political independence under the weakening of the Kingdom of Colchis. This period included a series of luxurious tombs and chapels built with groups of tombs. There were innovations in the craft industry, like the production of amphoras and tiles and changes in burial rules, such as burying grave goods like coins and amphoras for the deceased.The fourth stage, lasting from the second half of the 3rd century BC to the mid-1st century BC, witnessed fundamental changes. The burial of the upper strata of society within the city was stopped, and it became a temple city. All buildings of this period were for religious purposes. Ancient Vani was destroyed in the middle of the 1st century BC.

**Bazaleti (BZT)**

**A) General Location and Chronology:**

Bazaleti (42.07333,44.64861) is a village located on the left bank of the Tinishkevi River, on the Bazalet plateau. The area has been inhabited since the Stone Age, with remains of Neolithic stalls found here. Bazaleti is mentioned in “The Georgian Chronicles” dating back to the 2nd century BC and was a separate administrative unit. Bazaleti cemetery dates back to the 8th - 3rd centuries BC.

**B) Excavation history and description of burials:**

Archaeological excavations were carried out in 1976-77 by L. Tsitlanadze after the chance discovery of an ancient artifact. In 1987-89, the cemetery was excavated by the Mtianeti archaeological expedition of Eastern Georgia, headed by R. Ramishvili. About a hundred pit burials were excavated, most of which were located in the middle lane of the cemetery. The burials followed a uniform pattern: the deceased was laid at the bottom of an oval or rectangular pit, in a sharply bent position, predominantly on the right side, with their head towards the west or northwest. During the early period (7th - 4th centuries BC), black-burnt clay vessels were found in the tombs located in the eastern and northern parts of the slope. Iron weapons and bronze jewelry were also abundant in the tombs. In the group of burials dating back to the 4th - 3rd centuries BC, red-burnt and red-painted vessels were found. These types of vessels had first appeared in Georgia from the end of the 5th century BC.

**Nastakisi (NSK)**

**A) General Location and Chronology:**

Site Nastakisi (41.863,44.5742) is located on the left bank of Mtkvari, at the mouth of the Ksani River, near the railway station of the village of Ksani. The site of Nastakisi is bordered on the north by the Savaneti ridge, on the last peak of which the Ksani fortress is erected, on the south by the Mtkvari river. The Ksani river lies to the west, and the Naparsevi ravine is to the east. The archaeological findings at Nastakisi date from the Late Bronze Age to the Middle Ages.

**B) Excavation history and description of burials:**

Archaeological excavations on Nastakisi were carried out in 1978-80 under the leadership of A. Bokhochadze.

The oldest settlement found on the site is located on the mountain slope in the northwestern corner, dating back to the Late Bronze - Early Iron Age, which spans from the end of the 2nd millennium to the beginning of the 1st millennium BC. The excavation unearthed various grey clay vessels, flint sickle inserts, hand grinders, and other objects.

At the end of the 4th century BC, a large city-type settlement appeared on Nastakisi, which existed until the end of the 7th century.

In the Hellenistic period (4th - 2nd centuries BC), the Nastakisi vestry was connected directly to Nastakisi, with the settlement of Samadlo discovered on Samadlo Hill, and together with it constituted one vestry unit. This fortified outpost controlled important crossroads - one of these roads followed the Mtkvari river, while a north-south road connected Shida Kartli with Trialeti. Some believe that Samadlo-Nastakisi village is the "old Mtskheta" mentioned in the chronicle of "The Conversion of Kartli". In the 4th-3rd century BC, a 6-km-long river was drawn from the Great Ruskhmuli of Mukhran to irrigate the fields of Nastakisi.

On Nastakisi hill, there were capital buildings and buildings covered with tiles. The fragments of Doric order capital and architrave found on the site date back to the 4th-3rd centuries BC. Nastakisi clothing reached exceptional prosperity in the 4th-3rd centuries BC. Excavations revealed several clothing complexes, cellars, and other buildings of this period. Of particular note is the discovery of Christian churches built on a cobblestone foundation, as they are still the oldest Christian cult monuments in Georgia, dating back to the 2nd century AD according to archaeological data.

Archaeological finds reveal fragments of clay vessels painted with writing from the Hellenistic period, whited-painted pottery of the same period, dark-burnished vessels imported from Greece, and fish-ashets from Asia Minor dating back to 2nd century BC. Additionally, bronze arrowheads of the Scythian type were also found.

In addition to the settlement, on the field of Nastakisi site various types of burials were found at the Nastakisi site. A total of 145 tombs were excavated, which can be classified into five main types: 1) pit tombs, 2) urn burials, 3) cobblestone-built tombs, 4) stone tombs, 5) combined tombs. Of the 145 tombs excavated, 95 are pit burials. The pagan customs of burial prevail in the cemetery. Deads lie on their right or left side in a crouched position. Most of the tombs have been cataloged.

In the eastern part of the Nastakisi site, a medieval settlement and a church called "Nasparsevi" have been preserved.

**Nokalakevi (NOK)**

**A) General Location and Chronology:**

Village Nokalakevi (42.35722, 42.19389) is located in Senaki municipality, on the left bank of river Tekhuri, 16 km from Senaki.

**B) Excavation history and description of burials:**

Archaeological excavations were conducted from 1930 to 31, under the leadership of Alfons Maria Schneider, with the assistence of L. Muskhelishvili and G. Gozalishvili. This work lasted four months and uncovered part of the wall, several towers, and the eastern area of the Pentecostal Church. No further archaeological work was carried out at the site until 1950. A pit burial in the central part of the citadel was discovered in 1971.

In 1973, the medieval archaeological department of the Janashia Museum of Georgia conducted archaeological works at the site. In 1974, the southern part of the palace, which was built with large rocks, was cleaned. Research on the eastern defensive fence continued from 1975 to 1976. In 1974, excavations were carried out on the lower terrace located to the east of the eastern walls of Nokalakevi. The excavations in this area continued until 1977 and revealed 24 burials, including a pit burial that dates back to 4th-3rd centuries BC, and a  urn burial from the 3rd-2nd centuries BC.

Archaeological works were continued on the citadel and the lower terrace of Nokalakevi from 1978 to 1989. During this period, the remains of another church dating back to the middle of the 4th century were discovered on the lower terrace. Excavations of the eastern and central parts of the lower terrace revealed a large amount of archaeological material from different periods.

The excavations from 1990 to 1998 were limited in  scale and duration. Political processes in the country affected the expedition as well. Nokalakevi was one of the epicenters of the civil conflict. In 1990, only three areas were excavated.

In 1996, the expedition carried out excavations outside the settlement in the territory of the cemetery, which dates back to the Hellenistic period. No graves were discovered this year. At the same time, the cleaning of the circular building near the northern wall of the Royal Bath began and continued in the following years. The building was of the so-called two-chamber oven type.

Further archaeological expeditions have been conducted in Nokalakevi since the 21st century. A Georgian-Swiss expedition worked there briefly, while a Georgian-English expedition has been ongoing since 2001 untill 2022.

Several trenches have been excavated since 2001, including trenches A, B, C, D, E, F, and G. Trench A was excavated for the longest duration of 15 years, during which a multi-layered archaeological settlement was discovered. The archaeological study of Trench E was completed during the 2020 season, while active work is currently underway on Trench F and G. Nokalakevi is a multi-layered site with the oldest layers dating back to the 7th century BC. It is confirmed from the 8th century when zoomorphic figures with bipartite proteomes and a production population have been identified. Layers from the 6th to 4th centuries BC, 3rd to 1st centuries BC, and 4th to 6th centuries AD, along with a small number of developed and late medieval layers, have been recorded and studied.

In trench A a massive cultural layer dating back to the 3rd-1st centuries BC was discovered along with a settlement, and a cemetery. The layers contained an abundance of ceramic and building materials, construction debris in the form of piles of stone, foundation levels of the most important dwellings, and ceramic material from theinteriors of these dwellings, dating back to the 3rd-2nd centuries BC. Trench F was excavated in 2016, and it recorded layers dating back to the 5th-4th centuries and the 3rd-2nd centuries BC. The trench revealed a terraced outcrop where the Colchian houses were located. The excavation recorded a stone foundation and a layer of burnt clay plasters, which must have been created by a strong fire. The pottery found in this trench belongs to 3rd-2nd centuries BC in terms of its shape, color, and degree of firing.

**Khuntsi(KHT)**

**A) General Location and Chronology:**

Village Khuntsi (42.40308,42.43439) is located in Martvil Municipality, on the right bank of Tshnisskali. The Khuntsi castle complex dates back to the Early Middle Ages.

**B) Excavation history and description of burials**

In 2014, the Georgian-English archaeological expedition of Nokalakevi visited the Natsikhari area in the village of Khuntsi. This visit marked the beginning of an archaeological research project that continues to this day with the funding and support of the National Agency for the Protection of Cultural Heritage of Georgia.

In 2015, the archaeological expedition of Khuntsi cut 4 trial trenches in the territory of the castle. The layers revealed in the N1 and N3 trenches were particularly remarkable.

The N1 trench, located near the citadel tower, revealed a part of the floor covered with hydraulic solution about 20 cm beneath the surface. This area was temporarily preserved.

The N3 trench was dug near the northern wall of the castle, where a large amount of archaeological material dating back to the 4th-6th centuries AD was found in a 30-cm-thick layer covering an area of 1x3 m. Among the uncovered artifacts, there are both local and imported ceramic dishes.

In 2016, the expedition carried out work on five sections of the castle's territory, resulting in some interesting findings. In the southwestern part of the castle, a part of a building of unknown purpose was discovered, with unique architectural and construction details. Additionally, a 25-meter-long section of the northern wall of the castle was identified. The team also found a significant number of ceramic materials, such as amphorae, pitchers, luteriums, bricks, tiles, tiles. Most of these materials date from the 4th-6th centuries, indicating that the castle's existence should also be related to this period.

In 2017, work was carried out in five areas, including the expansion of trench N3 towards the south and east. In addition, four trial trenches were cut in the central, northern and eastern parts of the fort area.

In 2018, the excavation was carried out in three areas, namely trenches N1, N6 and N7. The structure and northern glass crypt discovered in previous years in trench N1 were entirely cleared, except for the crypt located in the center of the temple. A 20-meter section of a probably disintegrated wall fragment or stone wall was cleared in trench N6. And in trench N7, a quadrangular tower covered with a layer of earth was unearthed, which was the only visible structure on the ground surface until 2015.

In 2019, the walls that were attached to the tower located in the north-west of the hill, as well as its southern and northern extensions, were cleared. Additionally, a trial trench N9, measuring 3×4 meters, was cut, 20-30 m to the east of the structure that was identified in 2016. In this trench, a fragment of the fence wall was cleared. Another trench, N10, was dug on the north-eastern slope, where a fragment of the rampart wall was also exposed.

In 2020, archaeological works were carried out in four sections of the castle.

In trench N1, the stepped entrance to the crypt, which was discovered in 2016, and the crypt itself were excavated. This resulted in the discovery of the mixed skeletons of twelve individuals, which could only be distinguished after anthropological research.

In the N7 trench, the eastern section of the wall exposed last season was excavated, along with the stone pit located at 21 meters. Furthermore, the grave of a teenager near the alleged gate was also unearthed.

In the N8 trench, the northern section of the fence exposed in the N10 trench during the previous season was unearthed for a length of 10 meters.

Finally, in the N9 trench, the southern, zigzag-shaped section of the fence exposed in the N9 trench during the previous season was excateed for10 meters long.

In 2021, excavation works were carried out in three areas of the Khuntsi Castle. The N8 trench, which was initially cut in 2020, was extended to the northwest by 13 meters. The N9 trench, also cut in 2020, was expanded by 10 meters to the south, revealing a broken section of the fence.

Additionally, a new N10 trench was cut on the western edge of the Khuntsi Castle plateau. In trench N7, the south-eastern extension of the rampart was exposed and cleared. This section of the wall was fortified with pylons.In 2022, excavation work was carried out in two areas within the Khuntsi fortress. The N8 trench, which was initially dug in 2020, was combined with the N5 trench dug in the previous years and expanded to the west by 8 meters. Additionally, the study of trench N10, dug in 2021 and located on the western edge of the Khuntsi castle plateau, was continued.

**Atskuri (ATK)**

**A) General Location and Chronology:**

The site of Atsquri (41.72917,43.15944) is situated in the valley of the same name, on the left bank of the Mtkvari (Kura) River in the Akhaltsikhe district. The site is home to a stone mound, which contained a mass burial in a pit grave, dating to the end of the Middle Bronze Age. The discovery was made when the mound was damaged during work on the farmstead owned by N. and G. Gasitashvilis. The Atsquri burial belongs to the Trialeti culture and is estimated to date back to around 1600-1500 BC.

**B) Excavation history and description of burials:**

The Atsquri expedition, directed by V. Licheli, carried out archaeological excavations at the site in July 2001. The anatomical analysis of the human bones indicated that a total of 63 individuals were buried in the grave. Among them, there were 21 adult males, 38 adult females, and 4 juveniles. It is believed that this burial site belonged to an extended family.

During the early stage of the excavation, a mushroom-headed pin and fragments of black pottery were found. Owing to considerable damage to the central part of the barrow, the bones were concentrated along the central and western sides of the trench. Five human skulls were discernible in the mixed mass. Parts of broken bronze daggers and disc-headed pins were also found at this level. Bones of the lower limbs, lying above two severely deformed skulls, were found beneath the western section of the mound.

In the southern section of the mound, under the bones of the deceased, a one-handled cup, a bronze disc-headed pin, and an ornamented gold disc were found. The deceased individuals appear to have been laid on a bed that was placed over the pottery. Although no trace of wooden materials survived, the loose brown soil suggests that the funerary bier was made from wood or some form of matting. The skeleton of an ox was found mixed with human bones. In addition, in Square A's north corner at the same level, two human skulls were discovered facing southeast. Three pyramidal-headed bronze pins were found in situ on one of the skulls, clearly pointing to their use as hairpins. Two square and perforated bronze plaques lay in front of the face and were presumably attached to the clothes. At the eastern edge of the trench (Square B2), a dagger was found along with a bronze bracelet lying above it. A large disc-headed pin, a torque, and a deer skull with antlers were found at the boundary. Small carnelian beads were found under the torque.

A part of a human skeleton was discovered at the center of Square A2. Two daggers, not in their original position, were also found here. Furthermore, four bronze disc-headed pins were recovered from the chest area.

The Atsquri burial contained a total of 54 handmade clay vessels, including jars, pots, mugs, cups, and other drinking vessels.

A large number of metal artifacts, including 221 bronze items, were found in the burial. Among these were 11 weapons, including a spearhead, nine daggers, and a double-pronged fork. In addition, 29 bronze personal ornaments in the form of bracelets were also found.

Some of the deceased were buried with gold. There were 286 beads of various shapes, mostly carved from carnelian, and they came in a range of colors, including dark and light hues, transparent and opaque.

**Alaverdi(AVR)**

**A) General Location and Chronology:**

There are two archaeological sites situated near the village of Alaverdi (42.03246, 45.37717) , named "Alaverdi I" and "Alaverdi II."

"Alaverdi I" is a settlement dating back to the Early Bronze Age.  The archaeological layer of this site comprises 2cm-thick floor fragments of burnt clay, black-glazed pottery fragments, and flint sickle blades. On the other hand, "Alaverdi II" is a Middle Bronze Age cemetery, buried under three meters of sediments. The cemetery overlies and sometimes cuts through an Early Bronze Age layer.

**B) Excavation history and description of burials:**

Between 1969 and 1971, the Kakheti expedition led by K. Fitkhelauri of the Iv. Javakhishvili Institute of History, Archeology, and Ethnography conducted a study of the Alaverdi cemetery and settlement. The cemetery had two tombs with stone circles in the middle, surrounded by several other tombs. All the burials were pits filled with small stones and inhumations, with the deceased buried on either the right or left side and their heads facing west. Most of the burials contained clay vessels. Two samples included in this study, AVR001 and AVR002, were obtained from Alaverdi II.

**Skhalta (SKH)**

**A) General Location and Chronology**:

The Skhalta complex is located inside the village of Kinchauri, in the valley of the Skhalta (Khikhani) river, in the autonomous republic of Adjara, Georgia. The site is bordered by an unnamed dry gorge to the east.

The site consists of a settlement dating back to the 4th-3rd centuries BC, which coincides with the formation of the kingdom of Kartli (Iberia), and a contemporary cemetery.

**B) Excavation history and description of burials:**

The Skhalta Archaeological expedition, led by Z. Shatberashvili, partially investigated the Skhalta archaeological site in 2004-2005. The cemetery was excavated across an area of 150 square meters, extending to the unexcavated north. The burials were closely packed in a 15-meter strip, consisting mainly of cist burials with five or six rectangular rough-hewn basalt slabs. Additionally, five oval pit burials were found covered with slabs. These burial types were common in the Classical period and were widespread throughout Georgia. The burials were individual, with the head of the deceased towards the northwest, lying on the right or the left side. Burial N52 was an exception where the deceased was buried with the head to the east.

According to the paleoanthropological data, 46.7% of those buried in the Skhalta cemetery were men, 19.8% were women, and 23.4% were children. The average life expectancy among men was 44.4 years, while among women, it was 39.7 years. The average height of adult males wa 167.2 cm, whereas that of females 157.9 cm. The height of the population of Skhalta was above average for the period.

Burial N25 contained a jug with a tubular handle, which is unique at Shkalta. It was discovered alongside circular iron bracelets, small bronze temple rings, bi-conical glass beads and a bronze finger ring with a twisted wire bezel. The jug's origins can be traced back to the Persian world and it is believed to be an imitation of the metal vessels of Luristan. This type of jug became common in Georgia from the 5th-4th centuries BC, making this burial possibly the earliest in the cemetery. It is noteworthy that this burial was deeper than the others and was constructed using well-cut slabs of white sandstone instead of basalt. It is difficult to determine its exact date, but it is presumed to date back to the end of the 5th-4rd centuries BC.

Skhalta cemetery has been dated back to the 4th-3rd centuries BC based on the burial goods. Iron weapons, notably spearheads and axes, were among the items recovered. There are two types of spearheads found at the site:the first type has an elongated shape with a narrow head and a ridge, which is common in Georgia during the 6th-4rd centuries BC. The second type has a rhomboid head and a ridge, with the head and the butt being of the same size, while the middle part of the head being wide. Such spearheads have been found at Etso in Kvemo Kartli, and are also known from sites of the 4th-3rd centuries BC in other regions of Georgia and in western Azerbaijan.

The iron axes found at the cemetery have a similar shape with an oval hole for the shaft, a four-faceted butt, a slightly waisted body, and a narrow oval blade. These axes date back to the 4th-3rd centuries BC.

Iron tools such as knives and adzes were discovered at Skhalta. The knives come in three types: straight, bent and sickle-shaped. An iron adze, which was used for woodwork, was found in one damaged burial. It is an extremely rare type, unparalleled any other contemporary sites in Kvemo Kartli. Its shape is similar to those used in the pre-Classical period. It is worth noting that there was little innovation among the iron weapons and tools and that they were similar to those used in the previous period.

**Samadlo (SDL)**

**A) General Location and Chronology**:

Samadlo (42.465833,44.31944) is situated in the Mtskheta municipality, adjacent to Ksani railway station, on the right bank of the Mtkvari River, atop two prominent hills. A settlement dating to the Late Bronze Age and Early Iron Age, along with burials dating to the 4th to 3rd centuries BC, was identified at Samadlo.

**B) Excavation history and description of burials:**

In 1912, an accidental discovery led to the unearthing of a remarkably opulent burial assemblage. Between 1962 and 1964, the archaeological expedition conducted by the S. Janashia State Museum of Georgia, under the leadership of I. Gagoshidze undertook reconnaissance work at the site. From 1966 to 1975, extensive excavations were carried out on both hills of the settlement. In 1984, the Mtskheta archaeological expedition of the Archaeological Research Center at the Javakhishvili Institute of History, Archeology, and Ethnography, led by A. Apakidze, successfully excavated a cult building dating back to the 2nd century BC, located in the southwestern part of the settlement. The initial habitation of Samadlo dates to the middle of the 2nd millennium BC. During the Late Bronze to Early Iron Age, the eastern hill of the settlement featured clay-plastered pillar buildings, revealing a diverse range of artifacts such as miniature clay vessels, prints, sacrificial wheels, and other objects. The oldest layer unveiled lustrous black and gray ceramics adorned with amorous designs, suggesting a possible connection to religious or cult practices. Among the notable discoveries at Samadlo, one of the terraces revealed a charnel house from the 4th to 3rd century BC. The walls of this charnel house, reaching heights of up to 2 m, were skillfully constructed using cobbles and crushed stone, reinforced with wooden posts. The upper part was built with an adobe, and the roof was composed of wood and tiles. Within the confines of charnel-house, two wooden sarcophagi were uncovered. In one sarcophagus, the remains of seven individuals were found, accompanied by a limited inventory including a silver earring, finger rings, a painted clay jar, and a bronze arrowhead. Unfortunately, the second sarcophagus had already been subjected to looting.

**Klde (KLD)**

**A) General Location and Chronology:**

The Klde settlement (41.674618, 43.034928) is situated on a terraced slope at the confluence of the Mtkvari and Potskhovi Rivers near the Turkish border in southwestern Georgia, along a major trade route that once linked the South Caucasus and eastern Anatolia. The site, encompassing a large multi-layer settlement and a cemetery, extends over 3,486 square meters and includes structures, graves, and storage pits. The excavations yielded excellent and extensive cultural material from the first millennium AD. The settlement appears to have been destroyed by fire and rebuilt several times. The last fire in the 7th century AD, possibly during the campaign of Byzantine Emperor Flavius Heraclius or during an Arab invasion, led to the abandonment of the site.

**B) Excavation history and description of burials:**

The AGT (Azerbaijan, Georgia and Turkey) Pipelines Archaeology Program excavated the site of Klde was excavated in the early 2000s during the construction of the Baku-Tbilisi-Ceyhan (BTC) crude oil and adjacent South Caucasus (SCP) natural gas pipelines. The structures excavated during the pipeline project appear to have been domestic and were constructed from stone with tile roofs. All the dwellings possessed hearths for cooking, generally located either in the center or corner of the structure. The settlement's layout leads archaeologists to believe that the structures also had a defensive purpose. Several stone sling bullets of different shapes and sizes may have been a means of defense against attackers.

Interment at some of the burial sites at Klde, which were concentrated in three separate areas, occurred in stone-lined pit graves, some of them edged with stone, while others were in wine jars. Many of the skeletons were lying on their backs, but others were on their sides in crouched positions. These differences mean the burials took place in at least three cultural periods and may reflect broad religious and other cultural changes over time. Indeed, in the region under the Kartli (Iberia) Kingdom, differences between pre-Christian and Christian funerary cultures shed light on the shift to Christianity, with some graves manifesting both Christian and pre-Christian funerary traditions.

**Table 4. Osteological context of Klde individuals**

| ID | AGE | Genetic Sex | Burial Number | C14 Date | Archaeological date | Pathology |
|---|---|---|---|---|---|---|
| **KLD002** | | F | Burial N91 | 435-478 | | |
| **KLD003** | | F | Burial N87 | | Late Antique, I-III AD | |
| **KLD004** | | U | Grave No64 (III) | | | |
| **KLD005** | | U | Grave No86 | | | |
| **KLD006** | **60-65** | M | Grave No64 | | 3-4 Century CE. | AMTL-18,19,30 |
| **KLD007** | **45-50** | M | Grave No73 (III) | | 3-4 Century CE. | |
| **KLD008** | **50-55** | M | Grave No73 (I) | | 3-4 Century CE. | |
| **KLD009** | **60-65** | F | Grave No64 (I) | | 3-4 Century CE. | |
| **KLD010** | **40-45** | M | 2742, Grave No64(II) | | 3-4 Century CE. | |
| **KLD011** | **65-70** | M | 2748, Grave No83 (III) | 421-540 | 3-4 Century CE. | |

| | | | | | | |
|---|---|---|---|---|---|---|
| **KLD012** | **60-65** | M | 2747, Grave No83 (II) | 445-602 | 3-4 Century CE. | |
| **KLD013** | **30-35** | M | 2749, Grave No83 (II) | 550-638 | 3-4 Century CE. | |
| **KLD014** | | M | 2746, Grave No83 | | | |
| **KLD015** | | F | Grave2 | | Late Antiquity (according to Georgian periods) | ACD,metopic suture, WB |
| **KLD016** | | F | Grave No83 (III) | | 3-4 Century CE. | AMTL:5,18,19,29 |

**Mukhatgverdi (MUK)**

**A) General Location and Chronology**:

The village of Mukhatgverdi (41.822222, 44.730833), in the Mtskheta municipality and community, is located on the right bank of Mtkvari River, at an elevation of 560 meters above sea level, and 6 kilometers away from Mtskheta. The cemetery Mukhatgverdi I dates back to the 4th to 8th centuries AD, with a single cenotaph dating back to the Middle Bronze Age.

**B) Excavation history and description of burials:**

In 1937, the Iv. Javakhishvili Institute of History conducted an archaeological expedition in the area of Mukhatgverdi. From 1975 to 1976, the Mtskheta expedition of the Archaeological Research Center, led by A. Apakidze, studied the early medieval cemetery site of Mukhatgverdi I near Mukhatgverdi. Additionally, between 1979 and 1983, an expedition near the village of Mukhatgverdi revealed and studied a multi-layered settlement known as Mukhatgverdi II.

During the excavations at Mukhatgverdi I, the team explored an area of 360 square meters and revealed a total of 56 tombs made of stone slabs, including one pit tomb covered with stone slabs. These elongated rectangular tombs featured flat roofs. Typically, each burial accommodated one to three deceased individuals, who were interred on their backs with their heads facing west. The tombs yielded a diverse array of artifacts, such as iron and bronze hinges and studs that were adorned with acorns, coral, or lamb heads. Additionally, various beads made of glass, paste, sardine, acorn, briquettes, amber, and other materials were found. Notable discoveries in the cemetery include bronze awls, bow ties, printing tools, Artemisia, fragments of iron belts, and other objects. The cemetery dates back to the 4th to 8th centuries AD.

Besides, the cemetery excavation uncovered a stone-built cenotaph from the Middle Bronze Age. The cenotaph yielded three brownish-gray burnt pots, two jars, salt, and an obsidian fragment.

**Narekvavi (NRK)**

**A) General Location and Chronology:**

Narekvavi Cemetery (41.875638, 44.720319) is situated at the 543rd-kilometer mark along the Baku-Sufsa oil pipeline route. Narekvavi Cemetery has revealed over 200 tombs spanning from the Late Bronze Age to the Early Iron Age. The predominant burial style is pit burials, characterized by rounded corners and circular barrows. Most individuals were interred in a contracted position on wooden beds.

**B) Excavation history and description of burials:**

The site has undergone extensive archaeological investigations by the Mtskheta expedition at various periods. In 1960, Kalandadze led the initial archaeological surveys. Subsequently, from 1989 to 1992, R. Davlianidze spearheaded the excavation efforts, followed by A. Afakidze's research in 1998-1999. The final phase of excavation, from 2000 to 2001, was under the guidance of V. Nikolaishvili.

Notably, in 1989, archaeological excavations were conducted during the construction of the Mtskheta-Gori road branch, leading to the discovery of 17 pit burials within a 100-square-meter area. These burials featured elongated rectangular pits with rounded corners, with the heads facing toward the northeast. Some tombs displayed evidence of superimposition, suggesting multiple interments over time. Two tombs contained cobblestones, which served as supports for the wooden beds.The burial complex yielded a diverse range of archaeological artifacts, including a bronze quiver, "nest" arrowheads, round-headed buckles, bent bracelets, an iron spearhead, single-edged blades, swords, compound-handled daggers, sardines, beads made of paste and agate, as well as various clay vessels.

Further excavation efforts designated an 800-square-meter area for exploration. Within this region, nine pit burials were uncovered, with four displaying cobblestones and the other five lacking such features. The most notable burial among them was a collective grave with a diameter of nine meters, containing the remains of ten individuals. This burial yielded nine clay vessels, an obsidian arrowhead, a bronze shaft, and a bone object.

By 1998, the Mtskheta Institute of Archaeology had excavated a total of 42 burials, with all but two consisting of individual interments. In these burials, the deceased were laid to rest on their right or left side, with their heads oriented towards the north or northeast.

In 2000, an unexpolred tomb was discovered at Narekvavi confirming the existence of graves associated with distinguished warriors. These burials contained archaeological materials commonly found in similar contexts, including engraved belts related to ritual, components of horse harnesses such as bridles and umbons, swords, sates, axes, spearheads, arrowheads, standard heads, deer statues, and ritual pipes. Of particular interest was an engraved belt in the ninth tomb, depicting horses tethered to a plow, with a warrior astride one of them. The ceramic assemblage recovered from the Narekvavi tombs displayed a range of colors, including grayish and black, and showcased intricate incised relief and pressure-stamped ornamentation.

Unfortunately, we don’t know exactly which of these burials indicate individual, which make aDNA data.

**Mogvtakari (MGV)**

**A) General Location and Chronology:**

Mogvtakari (41.8415244, 44.7075154) is located opposite the Mtskheta railway station, nestled at the base of the northern slope of Mtkvari. Historical records and archaeological evidence suggest that Mogvtakari was likely one of the densely populated districts during Mtskheta's time as the capital. According to Leonti Mroveli's "Life of the Kings," the district's origin is connected to Farnajomi, the fourth king of Georgians. Farnajomi relocated servants of fire and Mogun from Persia, establishing their settlement in Mtskheta, which is now known as Moguta. Archaeological excavations in Mogvtakari commenced in the 1940s. Over the years, various archaeological expeditions have explored a cemetery dating from the 1st to 3rd centuries and a settlement from the 3rd to 4th centuries.

**B) Excavation history and description of burials:**

The cemetery at Mogvtakari reveals a predominant presence of elongated quadrangular tombs constructed with flat, side-folded, and grooved tiles. The tomb roofs can be either flat or two-colored, depending on the tomb's size. The burial floors were typically covered with flat-sided tiles, although some tombs had clay floors. Notably, this distribution of burial types and concentration of tiles is distinct from other Mtskheta burial mounds. While burial rituals varied, a consistent pattern emerged with the deceased commonly being laid to rest with their hands and feet folded to the side.

Inside the tombs, archaeologists discovered a diverse array of pottery vessels, including thin-faced, pink, or straw-colored clay goblets, plain and cup-decorated clay goblets, fluted three-lipped goblets, and anvils. Additional artifacts found include gold earrings, three-pointed arrowheads, bronze quills, pendants with deer figures, a bronze mirror, a colorless saltwater vessel with a bone spoon, iron and bronze rings (including one iron ring bearing the Greek inscription "Magnificent Queen" or "Beautiful Lady"), beads and pendants made of various materials, as well as coins of Augustus and Gottarze.

The tomb types and artifacts suggest that the Mogvtakari tombs belonged to the affluent class, with access to luxurious tiles and imported items. The presence of non-local tile tombs, foreign objects or those closely resembling foreign origin, and the frequent discovery of "Charon's coin" placed in the mouths or hands of the deceased raise intriguing questions about the ethnic identity of those interred in the Mogvtakari tombs. Researchers believe they may have been foreigners, either Greeks or Hellenized Iberians.

South of the cemetery, archaeological excavations uncovered a settlement, revealing the remains of buildings constructed with large-sized broken stones and mud for walls and fences. To the east of the burial site, the remnants of a house emerged, exhibiting crushed and burnt clay pots and vessels, alongside charred skeletal remains. Additionally, the excavation team unearthed four adult pitchers outside the house, and they identified a hearth on the northeast side of the dwelling.

**Tsaghvli (TSG)**

**A) General Location and Chronology:**

Tsaghvli (42.12083, 43.69722), located in the municipality of Khashuri, lies in the Lopanistskali valley within the present-day village of Tsaghvli. Archaeologists excavated the cemetery located 2 km southeast of Tsaghvli village, on the left bank of the Tsaghvlura River, in the area known as "Mikelaant Verkhevi." Out of the tombs found in the "Mikelaant Virkhevi" cemetery, 94 can be dated to the end of the Middle Bronze Age, 52 to the transitional period between the Middle Bronze Age and the Late Bronze Age, and 4 to the Late Bronze Age.

**B) Excavation history and description of burials:**

Since 1977, the Archaeological Research Center of Khashuri municipality, led by Al. Ramishvili has conducted archaeological excavations in Tsaghvli village. The Archaeological Research

Center of Khashuri municipality, under the leadership of Al. Ramishvili successfully identified several cemeteries and settlements through extensive reconnaissance. These include the Satsikhurisgori settlement and cemetery in the village of "Mikelaant Virkhevi," as well as the multi-layered settlement and cemetery in Natsargora village.

At the Mikelaant Virkhevi cemetery, archaeologists discovered 150 stone pit graves characterized by rectangular shapes and rounded corners. Among the excavated tombs, archaeologists identified 100 as inhumation graves, 46 as cenotaphs, and they found four in a destroyed state. Most of the tombs served as individual burial sites, while two tombs contained the remains of two individuals each. Most of the tombs functioned as individual burial sites, although two tombs contained the remains of two individuals each.

Moreover, in five tombs, archaeologists verified the recumbent burial posture of the deceased. At the cemetery, archaeologists discovered approximately 450 different types of clay vessels. The findings also included weapons such as bronze lances and spearheads without threads, obsidian arrowheads, bronze jewelry, various types of fasteners (112 pieces), bracelets (21 pieces), and pendants of different shapes (90 pieces) made from materials such as sardine, agate, antimony, and paste beads (900 pieces).

**Arboshiki (AHI)**

**A) General Location and Chronology:**

Arboshiki(41.551111, 45.963611) is a village in Dedoplistsqaro Municipality, Kakheti region. A Middle Bronze Age cemetery near the village dates back to the 13th - 12th centuries BCE.

**B) Excavation History and Description of Burials:**

During the construction of the reservoir near the cemetery, it was discovered that the "Black puddle" had been damaged by a bulldozer. Seven heavily damaged pit graves were excavated at the cemetery, dating back to the 13th-12th BC period. Within these graves, the deceased were found resting with their limbs bent, either on their left or right side, with their heads positioned to the north. Archaeological artifacts including clay pots, bracelets, rings, needles, flint, and beads were unearthed. Pit N4, relatively better preserved, was covered with poles. Additionally, bones of small cattle were discovered within the same tomb alongside the other artifacts.

**Taribana (TRB)**

**A) General Location and Chronology:**

Taribana (41.43420806, 46.13162666)is situated in the municipality of Dedoflitskaro in Kakheti. A multiple burial was found at Taribana dating back to the Early Bronze Age.

**B) Excavation History and Description of Burials:**

A burial pit with gently rounded corners was unearthed, sloping from the southeast to the northwest. Within the tomb, the remains of 63 individuals were discovered, ranging in age from 4-5 to 60 years. These individuals were laid to rest with bent limbs, reclining on either the right or left side, with their heads facing various directions.

Among the artifacts found in the burial were a fragmentary vessel, an obsidian arrowhead, and a bone spindle whorl. The clay varied in texture, ranging from fine to coarse. The assemblage of vessels included tubs, mugs, drinking vessels, pots, and bowls. The tomb's entrance is connected by a dromos from the southeast, where a majority of the skeletons, predominantly children, were buried.

Adjacent to the northern wall of the chamber, a skull of either a goat or pig was uncovered. In the center of the tomb, it appears that several skeletons were likely wrapped in animal skins. The artifacts suggest a timeline placing the tomb in the Early Bronze Age.

**Manavi (MVI)**

**A) General Location and Chronology:**

Manavi (41.71833, 45.46278) is a village situated in the Sagarejo Municipality of Kakheti, Georgia. Nestled at the foot of the southwestern slope of the Gombor range, it acts as a central hub for surrounding villages, including Burdiani. A cemetery dating back to the Middle Ages was found near Manavi village.

**B) Excavation History and Description of Burials:**

In August 2021, approximately 0.5 kilometers northwest of Manavi village, at a location called 'Turisubani' (which is southwest of Manavi Castle and approximately 0.58 kilometers away), a tomb was discovered during construction work. The territory is owned by "Vine Art" LLC, with the specified tax code (S/C: 55.09.67.304). The plot, designated as agricultural, covers an area of 4718 square meters. Earthworks were halted immediately and reported to the National Agency for Cultural Heritage Protection of Georgia. A joint inspection of the site where the archaeological object was discovered was conducted, and plans for rescue archaeological works were formulated. At that time, reinforced concrete walls, covering several hundred square meters, had already been half-built for the purpose of arranging various buildings and structures. During earthworks in one of the sections, a tomb was unearthed and partially damaged (GPS coordinates: 536448.00 m E, 4619354.00 m N). This burial structure is of the stone box type, a common style from ancient times to the late Middle Ages. According to the initial inspection, the tomb was determined to be from the Middle Ages, based on the monolithic, large size of its roof tiles, characteristic of that period. Additionally, fragments of ceramic products were found scattered near the tomb, including developed and late medieval periods, as well as early medieval ceramic products.

In August 2021, rescue archaeological work commenced on the discovered cemetery. Burial N1 is of the stone box type. Its western side was displaced during heavy equipment work before the start of archaeological excavation. The tomb, constructed of large sandstone slabs with stone used in the southeast corner, measures 162x70 cm in interior dimensions, with a height of 65 cm. Osteological material of an individual was discovered within, lying on their back with the head facing west. This individual was accidentally covered by earth during the lifting of the roof slab. Visible were pelvis, femur, and tibia bones, but no burial inventory was found. The tomb, located in the northwestern part of the cemetery, contained the remains of three individuals, two of whom were buried Gulag-style, oriented on a west-east axis, while the skulls of the second and third individuals were placed in the western part of the tomb, south of the main skull. The remaining bone material was buried in the southeast corner. The skeleton of one exhumed individual had the upper limbs stacked at the waist, with the head tilted towards the south. Based on pelvic bones, this skeleton likely belonged to an adult male. The stone box, measuring 0.9 m wide and 1.7 m long (external dimensions), consists of four sandstone slabs and one 0.35 m long stone. Burial N2, also of the stone box type, comprised rock fragments and was discovered in the northeastern part of the cemetery, containing three individuals. The first individual was positioned on their left side in a crouched position, oriented on a west-east axis and facing north. The second individual was buried sprawled out, with their upper limbs stacked on their shoulders, with a skull found near the left shoulder. The rest of the skeletal remains were buried in the southeast edge of the tomb. Individuals #1 and #2 likely were adults, while #3 was a juvenile. Disintegrated fragments of a human skull were found at the western edge of the stone box, possibly damaged during their burial process.

Tomb N3, made of stone with correctly processed frames, is also of the stone box type. The covering of sandstone slabs was pushed into the tomb due to the weight of the bulk earth. The dimensions are 185X72 cm, with a depth of 55 cm. Four individuals were buried in the tomb, facing west-east on the line. Two dead bodies were laid side by side, in a distended position, facing north, with the skulls placed on the north side of the tomb. The bones were wrapped in the eastern part of the tomb, located in the western corner. According to the pelvic bones, the skeleton in the north probably belonged to a female, and the one in the south must have been a male.

Burial N4 was identified and cleaned in the center of the cemetery, with only the northern stones of the retaining wall of the tomb preserved, and the rest, including the osteological material, not identified.

Burial N5 was discovered and cleaned in the southwestern part of the cemetery. During cleaning, it was found that only one flat sandstone from the side and roof of the tomb remained, with the osteological material found in a small, highly fragmented state. The burial inventory consists of bronze and iron objects, including bronze bracelets, bronze and iron rings, bronze pins, iron buckles, bone ornaments (hairbands), and iron pins with serdolite heads.

**Tsaishi (TSH)**

**A) General Location and Chronology:**

The Tsaishi cemetery is located within the Tsatskhvi administrative unit of the Zugdidi municipality in Georgia. It lies approximately 12 km from Zugdidi, situated on the right side of the Khobi-Zugdidi highway. The geographic coordinates of the site are 42° 24I 55II N – 41° 47I 42II E(42.415278, 41.795000). The excavation and study of the site have revealed its occupation from the Late Bronze Age to the Early Iron Age. The site has been under systematic investigation since the 1970s, led by Professor Teimuraz Mikeladze.

**B) Excavation History and Description of Burials:**

The Tsaishi cemetery was discovered during a surface inspection, which led to the assumption that it was associated with an ancient cemetery culture due to the presence of human bone fragments and clay vessels. In 2001, a surface reconnaissance clarified the topography and exact location of the site. Subsequent small-scale excavation works in 2003 revealed the cemetery's structure, area, and distribution, indicating collective pit burials and individual graves.

The cemetery consists of surface cult squares and large collective burial pits or rock dams encircling them. These rock-cut pits, reaching depths of 1-2 meters, cover an area ranging from 20 m² to 80 m² and are often connected to cult squares by a dromos.

Two main pit-graves, N1 and N2, were excavated. Pit-grave N1 contained the remains of approximately 800 individuals and around 1300 items, including various artifacts made of stone, clay, copper, bronze, iron, silver, and gold. Adjacent to Pit-grave N1, Pit-grave N2 potentially held the remains of up to 2000 individuals. The composition of items recovered from Pit-grave N2 closely resembles that of Pit-grave N1, suggesting a similar ritualized arrangement.

In terms of dating, Pit-grave N1 is associated with the early utilization of iron resources, likely operating from the second half of the 8th century BC to the first half of the 7th century BC. Pit-grave N2 predates Pit-grave N1 and is chronologically linked to it, possibly dating back to the 1st century BC or earlier, indicated by the presence of archaic weapon forms and the proportion of bronze to iron artifacts.

**Igoeti (IGT)**

**A) General Location and Chronology:**

The Grakliani settlement site and burial ground are situated in the Caspi region, within the territory of Igoeti village, on a hill located on the right bank of the Lekhura River, in close proximity to the Tbilisi-Senaki-Leselidze highway. The site covers a significant portion of the territory and is positioned between two small rivers – Lekhura and Tortla. The discovery of the site occurred in 2008 during a salvage archaeological excavation conducted due to the expansion of the highway. Subsequent excavations have revealed the presence of a multi-layered settlement and graves dating from stone age to high Middle Ages. In 2021, further study of the burial site west of the III-IV terrace of the settlement was initiated.

**B) Excavation History and Description of Burials:**

Burial 19-3 is a pit burial that has suffered damage to its southern part. The tomb is oriented on a west-east axis. The skeletal remains within the tomb are extensively damaged. Two individuals are interred in the tomb, facing each other, with one atop the other. The upper individual's skeleton is particularly damaged, complicating identification, and they are buried with their head to the west on the right side, while the lower individual is positioned with their head to the east on the left side. Despite the severe damage, several iron objects were found in the chest area of the upper individual, though their identification is challenging due to damage. Additionally, a badly burnt jug was discovered in the western part of the tomb, along with two narrow bronze rings, potentially earrings, near the skull of the lower individual. The dimensions of the tomb measure 120 cm in length and 50 cm in width.

Burial No. 19-I-5, located beneath Burial No. 19-I-4, is another pit burial tilted on a west-east axis and enclosed by a stone wall. Only partial skeletal remains, including the skull, pelvis, and lower limbs, were identified, with the bones arranged in a non-standard manner. Notably, the lower limbs are positioned near the skull. Various artifacts were found within the tomb, including black-fired clay plates, pots, and vessels, suggesting ritual significance. Iron bars covering the lower limbs and a bronze hinge found nearby add to the burial's complexity. The dimensions of this tomb measure 120 cm in length and 60 cm in width.

**Varsimaantkari (VRS)**

**A) General Location and Chronology:**

Archaeological sites of Varsimaantkari(42.0830129, 44.6951935), settlement, and cemetery are located in the village of Varsimaan, at the site of Varsimaantkhedi (Dusheti municipality), dating back to the 5th-4th centuries BC.

**B) Excavation History and Description of Burials:**

Archaeological excavations were conducted in 1972 and 1981–1986, led by R. Ramishvili. Individuals with stone and earthen tombs have been identified, featuring pit burials. The cemetery is single-layered and was operational for a long period. The deceased were buried on their right side  or left side , with paired burials being rare. Four tombs contained a horse buried with its rider. Various artifacts were discovered in the burials, including clay vessels (bowls, cups, jars, ladles, platters), iron weapons (spearheads, quivers, arrowheads, knives), bronze items (shields, helmets, armor, bracelets, rings, buckles), as well as silver and gold jewelry. A contemporary settlement was found approximately 10 meters northwest of the cemetery, featuring stone foundation walls with raised platforms and a sunken hearth. The material obtained from the excavation is similar to that found at other archaeological sites in Georgia.

**Erkneti (ERK)**

**A) General Location and Chronology:**

Erkneti is located (42.223333, 43.847778) in the municipality of Znauri, on the left bank of the East Frone River, within the territory of the modern village of Erkneti.

**B) Excavation History and Description of Burials:**

Around 500 meters away from the village at "Vasastskaro," eight graves were excavated, comprising six pit graves, one pit grave, and one derg grave. The oldest among them is the pit burial, found on the lower horizon, which contained two skeletons with bent limbs buried horizontally in a red-painted pitcher. The burial inventory included a black-burnt pot, a cone-bottomed jug, back-shaped bracelets, a blue glass scaraboid, and multi-faceted prints, dating the tomb to the 2nd-1st centuries BC.

**Okrokana (OGR)**

**A) General Location and Chronology:**

Okrokana is located (41.36, 43.25) in the Samtskhe-Javakheti region, Aspindza municipality. Several hill burials and remains of hill burials have been excavated in Okrokana, Vardzia. Some of them were located under the Vardzia-Nyala highway. These burials date back to the 2nd millennium BC.

**B) Excavation History and Description of Burials:**

Burial chambers were built with stone, among which the N3 hill burial is outstanding. Under its stonework, a megalithic burial chamber was discovered, built with dry pile. The floor of the chamber is compacted with gravel, and the roof was false. The dimensions of the burial chamber are: length 4.5 m, width 2.2 m, height 3 m. The anthropological material in the tombs was poorly preserved; in some cases, the remains of two bodies were distinguishable. The burial material mainly consisted of clay vessels and bronze products. Most of the clay vessels are decorated with various kinds of ornaments. There are also small vessels for different purposes, all of which are sculpted by hand. Among the bronze items, it is worth noting the hinges and stemmed and unstemmed leaf-shaped dagger.

**Kistauri (KTR)**

**A) General Location and Chronology:**

Kistauri Cemetery (41.9925, 45.2875) is located south of the village of Kistauri, in the place called Kaklebi. It dates back to the end of the 2nd-1st millennium BCE.

**B) Excavation History and Description of Burials:**

In 1965, the Kakheti archaeological expedition of the Institute of History, Archaeology, and Ethnography named after Ivane Javakhishvili (led by Kiazo Fitskhelauri) studied 19 tombs. In pit graves sloping from west to east, the dead were buried with their hands and feet bent, men on the right side and women on the left side. Earthenware (jar, bowl, pots, tray), a bronze spearhead, bronze bracelets, and sardine beads were found in the burials.

**Bodbe (BBD)**

**A) General Location and Chronology:**

Bodbe Cemetery (41.6064581, 45.9334379) is located in Signaghi Municipality and dates back to the High Middle Ages.

**Bulachauri (BLK)**

**A) General Location and Chronology:**

Bulachauri Cemetery (42.050278, 44.762778) is located in Dusheti Municipality and dates back to the transition period of Late Antique to Early Middle Ages.

**Mtsketijvari (MKV)**

**A) General Location and Chronology:**

Mtsketijvari Cemetery (41.9896647, 43.5805375) is located in Khashuri Municipality and dates back to the Early Middle Ages.

**Rustavi (RTV)**

**A) General Location and Chronology:**

Rustavi Cemetery (41.533333, 45) dates back from the Late Bronze Age to the Early Middle Ages.

**Telavi (TEL)**

**A) General Location and Chronology:**

Telavi Cemetery (41.9275971, 45.482239) is located in Kakheti region and dates back to the Early Iron Age.

**Tetritskaro (TKA)**

**A) General Location and Chronology:**

Tetritskaro Cemetery (41.701289, 43.169381) dates back to the Middle Bronze Age.

**Lafanaantkari (IAF)**

**A) General Location and Chronology:**

Lafanaantkari Cemetery (42.0874479, 44.6946425) is located in Dusheti Municipality and dates back to the Early Middle Ages.

**Grakliani (GKL)**

**A) General Location and Chronology:**

Grakliani Cemetery (41.9964, 44.5742) is located in Kaspi Municipality and dates back to the Hellenistic period.

**Khtsisi (KHS)**

**A) General Location and Chronology:**

Khtsisi Cemetery (41.983898, 43.6732292) dates back to the Late Antique period.

**Murakebi (MRA)**

**A) General Location and Chronology:**

Murakebi Cemetery (41.7929991, 45.7569288) is located in Kakheti region and dates back to the Late Bronze Age.

**Ota (OTA)**

**A) General Location and Chronology:**

Ota Cemetery (41.61944, 43.301389) dates back to the Middle Bronze Age.

**Sagvarjile (SGV)**

**A) General Location and Chronology:**

Sagvarjile Cemetery (42.1801251, 42.9668848) is located in Imereti region and dates back to the Middle Bronze Age.
